# METTL3 drives telomere targeting of TERRA lncRNA through m^6^A-dependent R-loop formation: a therapeutic target for ALT-positive neuroblastoma

**DOI:** 10.1101/2022.12.09.519591

**Authors:** Roshan Vaid, Ketan Thombare, Akram Mendez, Rebeca Burgos-Panadero, Anna Djos, Daniel Jachimowicz, Kristina Ihrmark Lundberg, Christoph Bartenhagen, Navinder Kumar, Conny Tümmler, Carina Sihlbom, Susanne Fransson, John Inge Johnsen, Per Kogner, Tommy Martinsson, Matthias Fischer, Tanmoy Mondal

**Author notes:** **Equal contribution** (Roshan Vaid, Ketan Thombare). Corresponding author,. Department of Clinical Chemistry, Bruna Stra□ket 16, Sahlgrenska University Hospital, Gothenburg University, Gothenburg-41345, Sweden.

## Abstract

Telomerase-negative tumors maintain telomere length by alternative lengthening of telomeres (ALT), but the underlying mechanism behind ALT remains poorly understood. A proportion of aggressive neuroblastoma (NB), particularly relapsed tumors, are positive for ALT (ALT+), suggesting that a better dissection of the ALT mechanism could lead to novel therapeutic opportunities. TERRA, a long non-coding RNA (lncRNA) derived from telomere ends, localizes to telomeres in a R-loop-dependent manner and plays a crucial role in telomere maintenance. Here we present evidence that RNA modification at the *N*^6^ position of internal adenosine (m^6^A) in TERRA by the methyltransferase METTL3 is essential for telomere maintenance in ALT+ cells, and the loss of TERRA m^6^A/METTL3 results in telomere damage. We observed that m^6^A modification is abundant in R-loop enriched TERRA, and the m^6^A-mediated recruitment of hnRNPA2B1 to TERRA is critical for R-loop formation. Our findings suggest that m^6^A drives telomere targeting of TERRA via R-loops, and this m^6^A-mediated R-loop formation could be a widespread mechanism employed by other chromatin-interacting lncRNAs. Furthermore, treatment of ALT+ NB cells with a METTL3 inhibitor resulted in compromised telomere targeting of TERRA and accumulation of DNA damage at telomeres, indicating that METTL3 inhibition may represent a therapeutic approach for ALT+ NB.

## Introduction

RNA modification at the *N*^6^ position of internal adenosine (m^6^A) is a crucial regulatory mechanism involved in RNA splicing and various steps of RNA metabolism. m^6^A modification in RNA contributes to a wide range of cellular functions, including stress response, DNA damage, cellular differentiation, and the regulation of oncogenes and tumor suppressor gene functions (1). METTL3, in complex with METTL14 and WTAP, serves as the key enzyme responsible for depositing m^6^A in RNA. The deposition of m^6^A is guided by active chromatin marks such as H3K36me3, along with adaptor proteins RBM15 and VIRMA (also known as KIAA1429), which facilitate the recruitment of the METTL3 complex to RNA substrates for m^6^A modification (2-4). While the "DRACH"-like motif is the preferred site for m^6^A modification, METTL3 can also utilize other non-canonical motifs for m^6^A deposition (5). The dynamic regulation of m^6^A modification involves the activity of m^6^A demethylases such as FTO and ALKBH5, which can remove m^6^A from transcripts (6,7). Once m^6^A is deposited in RNA, reader proteins in the nucleus (e.g., YTHDC1, RBMX, hnRNPA2B1) and cytoplasm (YTHDF1/2/3) recognize m^6^A residues and guide functional complexes to the modified RNA (1,8,9). Recent studies have demonstrated that the chromatin binding of METTL3 and METTL14 to promoter and distal enhancer elements plays a regulatory role in the transcriptional process (10). Additionally, m^6^A modification in retrotransposon-derived transcripts has been found to modulate local chromatin structure, thereby influencing transcriptional regulation (5,11). The RNA strand of the RNA:DNA hybrid containing an R-loop structure is enriched with m^6^A modification, and depletion of METTL3 affects the overall level of R-loops (12,13). However, further investigation is needed to understand how m^6^A modification in RNA modulates the level of R-loops and the functional significance of the presence of m^6^A in the R-loop structure.

The impact of m^6^A modification on the function of long non-coding RNAs (lncRNAs) has been described in several studies (14,15). For instance, the *cis*-acting lncRNA Xist, which localizes to the inactive X-chromosome, undergoes m^6^A modification at multiple residues in both the "DRACH" and non-"DRACH" motif context. The adaptor protein RBM15 recruits the METTL3/METTL14 complex to deposit m^6^A on Xist RNA. The reader protein YTHDC1 has been shown to recognize m^6^A in the UCG tetra-loop of Repeat A within Xist RNA and is required for Xist-mediated gene silencing (3,16). Additionally, m^6^A modification has been detected in the non-canonical GGAAU motif of the HSATIII lncRNA, playing a critical role in the sequestration of splicing factors in nuclear stress bodies (17). Widespread m^6^A modification has also been detected in several chromatin-bound RNAs, including enhancer RNAs and promoter-associated RNAs (18). The m^6^A modification of enhancer RNAs is necessary for YTHDC1 and BRD4-mediated transcriptional activation of neighboring genes (19). However, the mechanisms through which m^6^A modification influences the chromatin localization of lncRNAs to specific genomic loci remain poorly understood.

TERRA (telomeric repeat-containing RNA) is a lncRNA derived from the subtelomeric ends that contain telomeric repeats in its mature transcript (20). TERRA promoters can be found within 1-2 kilobases (kb) from telomere ends or several kb away from the telomere end. TERRA lncRNAs are transcribed by RNA polymerase II (RNA Pol II), which moves towards the telomere ends and incorporates telomeric repeats containing UUAGGG sequences (21,22). TERRA has been shown to localize to telomeres in both *cis*- and *trans*-acting manners, and the formation of RAD51/BRCA1-driven R-loops is required for this process (20).

TERRA transcription is upregulated in tumor cells that are positive for alternative lengthening of telomeres (ALT+), and it is believed to contribute to the maintenance of long telomeres. However, the additional factors involved in R-loop-mediated TERRA targeting in ALT+ cells are still unknown. Understanding the chromatin targeting of TERRA could provide insights into the mechanisms utilized by other *cis*- and *trans*-acting lncRNAs for their chromatin recruitment. Since R-loop formation can be regulated in an m^6^A-dependent manner, we aimed to investigate whether m^6^A modification plays a role in TERRA function. It is also worth noting that the telomere capping protein TRF2 has been found to interact with m^6^A-related proteins (RBM15 and ALKBH5), although the significance of this interaction remains unknown (23,24). Additionally, recent studies have shown that G-quadruplexes derived from TERRA repeats and other G-quadruplex-forming sequences can bind METTL14, but the role of such sequences in driving m^6^A modification in TERRA is not yet understood (25).

Neuroblastoma (NB) is the most common extracranial solid tumor in children and exhibits high clinical and biological heterogeneity (26). The presence of telomere maintenance mechanisms in NB has been proposed as a factor contributing to the heterogeneity of NB tumors and disease outcomes. Telomere maintenance mechanisms is strongly associated with high-risk NB, whereas they are absent in spontaneously regressing low-risk tumors (27). In the majority of high-risk cases, telomere maintenance is achieved through the telomerase pathway, which is transcriptionally induced by oncogenes such as *MYCN* or by genomic translocations that lead to enhancer hijacking events (28). However, in approximately one-third of high-risk NB cases, telomeres are maintained through the ALT pathway, characterized by long telomeres and ALT-associated promyelocytic leukemia nuclear bodies (APBs) (27,29). Despite recent advancements, the molecular mechanisms underlying ALT are still poorly understood (30). Therefore, gaining a deeper understanding of ALT+ tumor biology and systematically investigating the molecular mechanisms of ALT may help identify potential therapeutic targets for these clinically unfavorable high-risk NB tumors, which urgently require improved treatment options (29,30).

In this study, we investigated the role of m^6^A RNA modification in TERRA and its significance in R-loop formation in ALT. Further, we demonstrate that METTL3-mediated m^6^A modification of TERRA is essential for telomere maintenance in ALT+ NB cells and propose that inhibiting METTL3 could be a potential therapeutic option for ALT+ NB tumors.

## Materials and methods

### Cell culture

U-2 OS cells (Sigma-Aldrich) were cultured in Dulbecco’s modified Eagle’s medium (DMEM) supplemented with 10% fetal bovine serum (FBS) and penicillin/streptomycin. NB cell lines SK-N-FI (Sigma-Aldrich), SK-N-AS (ATCC), SH-SY5Y (CLS Cell Lines Service), CHLA-90 (CCOG), IMR-32 (CLS Cell Lines Service), and SK-N-BE(2) (DSMZ) cells were cultured in specific media. SK-N-FI, SK-N-AS, and SH-SY5Y cells were cultured in DMEM supplemented with 10% FBS, 0.1 mM Non-Essential Amino Acids (NEAA), and penicillin/streptomycin. CHLA-90 cells were cultured in DMEM supplemented with 20% FBS, 1x ITS (Gibco), and penicillin/streptomycin. IMR-32 cells were cultured in Minimum Essential Medium (MEM) supplemented with 10% FBS, 1 mM sodium pyruvate, 1x GlutaMAX (Gibco), and penicillin/streptomycin. SK-N-BE(2) cells were cultured in DMEM/F-12 media supplemented with 10% FBS, 1x GlutaMAX, and penicillin/streptomycin. All cell lines were routinely tested for mycoplasma contamination and confirmed to be mycoplasma negative.

### Plasmids, siRNAs, and molecular cloning

METTL3 (sh1 and sh2) and Control shRNA in pLKO.1-TRC cloning vector were purchased from Sigma-Aldrich to generate stable METTL3-KD cells. shRNA sequences are provided in Supplementary Table S1. To generate DOX inducible METTL3-KD cells in ALT+ NB METTL3- sh1 sequences were cloned in the Tet-pLKO-puro vector (21915, Addgene). As a control for inducible shRNA KD, we used pLKO-Tet-On-shRNA-Control plasmid (98398, Addgene). dCasRx-FTO^WT^-HA plasmid was received as a generous gift from Dr. Wenbo Lís laboratory (University of Texas Health Science Center, Houston, USA). The dCasRx-FTO^MUT^, dCasRx-METTL3^WT^, dCasRx-METTL3^MUT^, and dCasRx-hnRNPA2B1 plasmids were cloned using dCasRx-FTO^WT^-HA as the backbone vector. The plasmid was linearized using PCR, and the primers used were specifically designed to amplify the vector backbone (dCasRx-HA), excluding the FTO^WT^. The coding sequences of FTO^MUT^ (FTO catalytically dead mutant - H231A and D233A) METTL3^WT^ (METTL3 shRNA resistant wild-type), METTL3^MUT^ (METTL3 shRNA resistant catalytically dead mutant - APPA), and hnRNPA2B1 were custom synthesized (sequence information provided in Supplementary Table S1) as gene fragments from IDT (gBlocks™). The dCasRx-HA vector backbone and the insert were assembled using NEBuilder® HiFi DNA Assembly (E2621, NEB). Active CasRx was obtained from Addgene (138149). The TERRA and non-template control (NTC) gRNAs were cloned in pLentiRNAGuide_002 (138151, Addgene). Sequences of guide RNAs are provided in Supplementary Table S1. TRF1-mCherry plasmid was received as a generous gift from Dr. Emilio Cusanelli (University of Trento, Italy). Pre-designed siRNAs were purchased from either Sigma-Aldrich or Thermo Fisher Scientific and details are provided in Supplementary Table S1.

#### cDNA synthesis and Quantitative PCR (qPCR)

RNA was converted to cDNA using the High-Capacity RNA-to-cDNA Kit (4387406, Thermo Fisher Scientific). qPCR was performed with diluted cDNA (10-fold dilution) as a template and gene-specific PCR primers (Supplementary Table S1) mixed with Power SYBR Green Master Mix (4367659, Thermo Fisher Scientific) using a Quant Studio 3 thermocycler (Thermo Fisher Scientific). The expression values presented for each gene are normalized to *GAPDH* using the delta-delta Ct method.

#### Slot blot assay

Nucleic acids (RNA or DNA) samples were loaded in Bio-Dot SF Microfiltration Apparatus (Bio-Rad) and allowed to bind onto the membrane. Nucleic acids were UV-crosslinked to the membrane and incubated with prehybridization buffer DIG Easy Hyb (11603558001, Roche,) in a hybridization tube at 37° C for 30 min with rotation. Prehybridization buffer was replaced by hybridization buffer (prehybridization buffer containing 20-50 nM DIG-labeled probe [TERRA/Alu/LINE1]) and the membrane was hybridized at 37°C overnight with rotation. After hybridization, the membrane was washed with 2x SSC containing 01% SDS for 4×10 min at 37°C and equilibrated with washing buffer (0.1 M Maleic acid, 0.15 M NaCl, pH 7.5, 0.3% Tween 20) for 2 min at RT. Next, the membrane was incubated in a blocking buffer (1% (w / v) Roche Blocking Reagent, 0.1 M Maleic Acid, 0.15 M NaCl, pH 7.5) for 30 min at RT. The blocking buffer was replaced with antibody solution (1: 20,000 dilution of anti-DIG antibody conjugated with alkaline phosphatase in blocking buffer) for 30 min at RT. The membrane was washed twice 2×15 min with washing buffer. The membrane was equilibrated in a detection buffer (0.1 M Tris-HCl, 0.1 M NaCl, pH 9.5) and incubated with CDP-Star Chemiluminescent Substrate (Sigma-Aldrich) inside a hybridization bag. ChemiDoc XRS+ system (Bio-Rad) and ImageLab software were used to detect the signal and quantify the bands.

When detecting telomeric DNA we used DIG-labeled TERRA probe [5´-(TAACCC)_5_-DIG], which in the figure legend is indicated as telomeric probe.

#### TERRA RNA levels using slot blot assay

To check the level of TERRA in different cell lines, tumors, and conditions, isolated total RNA from different samples were first quantified on NanoDrop™ 2000 Spectrophotometer (Thermo Fisher Scientific) and diluted to 30 ng/μl concentration. The diluted RNA was again quantified on the Nanodrop to verify the concentration. The diluted RNA was then loaded onto slot blot in amounts as mentioned in the figures, and 1 µl of the diluted RNA which corresponds to 30 ng was used as a loading control. The slot blots were probed with a TERRA probe, developed as mentioned above, and normalized to the 28s rRNA signal from the loading control. To obtain the 28s rRNA signal the RNA samples were subjected to electrophoresis on a chip (ScreenTapes) using TapeStation 4200 (Agilent) according to manufacturer instructions.

#### m^6^A RIP followed by sequencing (m^6^A RIP-seq)/ slot blot/ qPCR

m^6^A RIP was performed on RNA as previously described by (31). In brief, for the m^6^A RIP-seq experiments the cellular (15 μg/IP) and tumor RNA (4-5 μg/IP) were supplemented with 10 ng or 3 ng of bacterial RNA respectively as a spike-in control. For m^6^A RIP followed by slot blot bacterial spike-in RNA was not added. The RNA was then subjected to fragmentation using RNA fragmentation reagents (AM8740, Thermo Fisher Scientific). m^6^A RIP was also performed in one instance without the fragmentation step. RNA immunoprecipitation (RIP) was performed using antibodies against m^6^A modification (202003, Synaptic systems) or IgG (sc-2027, Santa Cruz) in RIP buffer (150 mM NaCl, 10 mM Tris-HCl pH 7.5, 0.1% IGEPAL CA-630 and supplemented with RNase inhibitor just before use) for 4 h at 4°C. IgG antibodies served as a negative control for the immunoprecipitation experiments. The immunoprecipitated RNA-antibody complex was captured by incubating with protein A and G magnetic beads (10002D, 10004D, Thermo Fisher Scientific) which were washed twice in RIP buffer. After 2 h incubation, the magnetic beads were washed thrice with RIP buffer for 10 min. Immunoprecipitated RNA was eluted either by competitive elution (for RIP-seq) as previously described (31) or by direct extraction using TRIzol (for slot blot).

For the m^6^A RIP slot blot, the input RNA and the immunoprecipitated RNA were directly subjected to the slot blot assay described above.

For the m^6^A RIP qPCR, the input RNA and the immunoprecipitated RNA were converted to cDNA, and qPCR was performed with primers (Supplementary Table S1) as described above.

For m^6^A RIP and RNA-seq, sequencing libraries for total RNA-seq/input for m^6^A RIP-seq were prepared from 10 ng of the fragmented RNA parallelly along with m^6^A RIP samples using SMARTer Stranded Total RNA-seq Kit V2, Pico Input Mammalian (Takara Bio) according to manufactures instructions. All the libraries were single-end sequenced (1×88 bp) on an Illumina NextSeq 2000 platform at the BEA core facility, in Stockholm, Sweden.

#### Direct RNA sequencing using Oxford Nanopore Technologies (ONT) long-read sequencing

Total RNA isolated from Control-sh and METTL3-KD cells, 20-25 µg was directly sequenced on the Oxford nanopore platform. In brief total RNA quantification and quality control were performed on the Fragment Analyzer 5300 system and showed RNA integrity number (RIN) of 10 for both samples. Poly(A) RNA Selection Kit V1.5 (Lexogen) was used to enrich polyadenylated transcripts with minor changes to the protocol. All the processing volumes were doubled, and 4 µg of Control-sh or METTL3-KD RNA was used as an input in a single reaction. Each sample was batch-processed in 5 separate reactions that were combined at the final elution step. Control-sh and METTL3-KD poly(A)-enriched RNA was analyzed on Fragment Analyzer showing remaining rRNA contamination to be 11% and 21% of mass for each sample, respectively. Enriched poly(A)-RNA was then purified using 1x of RNAClean XP (Beckman Coulter) beads to concentrate the sample and remove very short RNA fragments and eluted into 11 μl of water. Qubit™ RNA High Sensitivity kit (Invitrogen) was used to quantify the RNA before direct RNA sequencing. SQK-RNA002 (ONT) kit was used for library preparation and followed according to the manufacturer’s protocol. Briefly, 400 ng of poly(A)-enriched RNA was used as an input, and recommended reverse-transcription step was performed. Control-sh and METTL3-KD libraries were loaded on individual FLO-MIN106 (ONT) flow-cells and sequenced on GridION Mk1 device (ONT) for 72 h yielding 1.65 and 1.35 million long-reads, respectively.

## Metagene analysis

Genome-wide distribution of m^6^A sites from Illumina and ONT data was analyzed using the plyranges package (32) to map the m^6^A coordinates to the genomic feature coordinates (5’ UTR, CDS, 3’UTR) obtained from the T2T-CHM13 v1.1 GTF annotation files. Mapped coordinates were further translated to their relative metagene positions. The peak density distribution for all m^6^A sites along metagene coordinates was calculated using the density function in R.

### Mappability at chromosome ends

To account for the repetitive nature of chromosome ends, we calculated the mappability of short and long reads using GenMap (33). This tool estimates the occurrence of k-mer sequences of a given length across the genome and computes a probability score proportional to its occurrence at different parts of the genome. A mappability score of 1 indicates that a sequence could align to a unique genomic location, while a low mappability value indicates that a sequence can map to one or many sites in the genome. The computed mappability scores for Illumina (75 bp) and Nanopore (600 bp) reads were averaged over fixed-size windows using BEDTools v2.30.0 (34) and further used to normalize the number of reads aligned at each chromosome end to the likelihood of mapping at one or multiple locations.

#### Analysis of TERRA expression

To analyze the expression towards chromosome ends in a strand-specific manner, the reads from Illumina alignments mapping to subtelomeric 30 kb regions from chromosome q and p arms were split by strand and filtered from all T2T alignments after duplicate removal. Alignments from direct RNA ONT data were filtered using positive strand information. The number of reads mapping to subtelomeric regions on each chromosome was quantified using the multicov function from BEDTools v2.30.0(34). Expression values were further converted to counts per million (CPM) to normalize for differences in library sizes across samples and normalized to the mappability scores computed using GenMap to adjust for the likelihood of short and long reads to align at one or multiple repetitive sites at each chromosome end. Chromosome ends were classified into two categories (active/inactive) according to their relative CPM expression values, being classified as ‘active’ if their expression was greater than the median normalized expression value across all chromosome ends at the control condition and ‘inactive’ otherwise. Enrichment of telomeric repeats in m^6^A RIP-seq from Control-sh, METTL3-KD, and IgG data was calculated using TelomereHunter v1.0.4 using a default repeat threshold of at least 10 non-consecutive repeats to search for TTAGGG, TGAGGG, TCAGGG, and TTGGGG repeats (35). Filtered TelomereHunter read counts were normalized using CPM values.

#### Immunofluorescence (IF) and Proximity ligation assay (PLA)

U-2 OS and NB cell lines were seeded on a coverslip in a 24-well plate at a density of 100,000 and 200,000, respectively. In case of NB cells, coverslips (MENZCB00120RAC20, Epredia) were coated with 50 μg/ml collagen (5005, PureCol, Type I Collagen Solution) at RT for 1 h. Cells were fixed using 4% formaldehyde for 10 min followed by 2x washes with PBS and stored in PBS at 4°C until further use. IF was performed using a standard protocol. Briefly, cells were permeabilized for 10 min with 0.25% Triton X-100 followed by 3x washes with PBS-T. Cells were blocked with 3% BSA in PBS-T for 30 min and left with primary antibody overnight at 4°C. IF was continued the next day starting with 3x PBS-T washes followed by incubation with respective secondary antibodies for 1 h at RT, and then 3x PBS-T washes. Coverslips were dried and mounted on a slide using ProLong Gold Antifade Mountant with DAPI (P36931, Thermo Fisher Scientific). Images were taken using EVOS M7000 microscope (Thermo Fisher Scientific). Information on the antibodies used is provided in Supplementary Table S1.

For PLA cell fixation and permeabilization are the same as above. PLA was performed using a Duolink® PLA kit (DUO92014, Sigma-Aldrich) according to the manufacturer’s protocol. As a background control, a single antibody was used in this assay. For R-loop PLA with S9.6 and TRF2 antibodies, cells were pre-treated with RNase A after permeabilization for 30 min at 37°C to reduce the backgrounds. Information on the antibodies used is provided in Supplementary Table S1. Images were taken using EVOS M7000 microscope (Thermo Fisher Scientific).

#### RNA-FISH

U-2 OS and NB cell lines were seeded on a coverslip in a 24-well plate at a density of 100,000 and 200,000, respectively. In case of NB cells, coverslips (MENZCB00120RAC20, Epredia) were coated with 50 μg/ml collagen (5005, PureCol, Type I Collagen Solution) at RT for 1 h. Cells were fixed using 4% formaldehyde for 10 min followed by 2x washes with PBS and stored in 70% ethanol at 4°C until further use. RNA-FISH was performed to detect TERRA and Stellaris RNA-FISH protocol for adherent cells was used with some modifications. Briefly, cells were permeabilized for 10 min with 0.25% Triton X-100 followed by a wash with PBS. Cells were then incubated for 2-5 min with wash buffer A (Stellaris RNA-FISH Wash Buffer A, SMF-WA1-60, Biosearch Technologies). Hybridization was carried out at 37°C for 4 h by incubating cells with hybridization buffer (Stellaris RNA-FISH Hybridization Buffer, SMF-HB1-10, Biosearch Technologies) containing TERRA probe (TAACCC)_7_-Atto488 in a humidified dark chamber. Cells were then washed 2x with wash buffer A at 37°C for 15 min, followed by 2-3 min incubation at RT with wash buffer B (Stellaris RNA-FISH Wash Buffer B, SMF-WB1-20, Biosearch Technologies). Coverslips were dried and mounted on a slide using ProLong Gold Antifade Mountant with DAPI (P36931, Thermo Fisher Scientific). Images were taken using EVOS M7000 microscope (Thermo Fisher Scientific).

### Drug treatments

U-2 OS cells were treated with Bleomycin 10 µg/ml containing media for 4 h. U-2 OS cells were treated with Actinomycin D (10 µg/ml) for the indicated time points and RNA was isolated from the treated cells to perform TERRA slot blot assay. METTL3 inhibitor STM2457 (10 µM, HY-134836), UZH1A (50 µM, HY-134673A), and UZH1B (50 µM, HY-134673B) were treated for the different time points as indicated in the figure legends. METTL3 inhibiuted cells were used for TERRA RNA-FISH, Telomere DNA-FISH/IF or Chromatin Immuno precipitation (ChIP). For detailed method related to ChIP see supplementary methods.

### Purification of TERRA and RNA-MS

#### TERRA pulldown

Capturing a specific RNA from total RNA was performed as described previously (36). In brief, ∼300 μg of DNase I treated RNA isolated from U2O-S cells was mixed with 400 pmoles of biotinylated DNA probe (Supplementary Table S1) against TERRA or Luciferase probe (negative control) in 5x SSC buffer and denatured at 90°C for 3 min. This mixture was immediately cooled down on the ice and again heated at 65°C for 10 min after which the samples were left to cool down at RT to allow hybridization of the probe. To pulldown the RNA hybridized with biotinylated DNA probes, MyOne Streptavidin C1 magnetic Dynabeads (Thermo Fisher Scientific) were added and incubated for 30 min at room temperature. The magnetic beads were washed once with 1x SSC buffer and thrice with 0.1x SSC buffer. To elute the captured RNA, the magnetic beads were resuspended in nuclease-free water, and heated at 75°C for 3 min. The eluted RNA was treated with RNase-free DNase I for 30 min at 37°C. Finally, the RNA was precipitated with 3 volumes of 100% ethanol, 0.5 M ammonium acetate, and GlycoBlue (Thermo Fisher Scientific) at -80°C for 2 h. The precipitated RNA was washed with 80% ethanol and resuspended in nuclease-free water.

The RNA captured using TERRA or Luciferase probe (Luciferase) was first checked for specificity by either sequencing or slot blot assay using TERRA probe (see slot blot assay). Probes against Alu and LINE-1 RNA were used as control. Probe sequences are provided in Supplementary Table S1.

Further, the captured TERRA along with synthetic RNA oligo (Supplementary Table S1) were digested with RNase T1 and analyzed on a mass spectrometer (MS).

#### RNA-MS analysis

RNA-MS was performed using an Easy-nLC 1200 LC system (Thermo Fisher Scientific) coupled to Orbitrap Fusion™ Eclipse™ Tribrid™ mass spectrometer. The analytes were trapped on a 2□cm x 100□µm Acclaim PepMap C18 precolumn (particle size 5□µm; Thermo Fisher Scientific) and separated on a 30□cm x 75□µm analytical column packed in-house with 3 μm Reprosil-Pur C18 material (Dr. Maisch, Germany) at 300 nL/min flow using a stepwise elution profile during 45 min. Solvent A was 5□mM di-n-butylamine and 8□mM acetic acid in H_2_O, and solvent B was 70% methanol, 5□mM di-n-butylamine, and 8□mM acetic acid. The column oven temperature was 50°C. Nano-Flex ion source (Thermo Fisher Scientific) was operated in negative ionization mode at 1.8□kV with the ion transfer capillary temperature 275°C. The MS instrument was run in a targeted MS/MS setting with switching quadrupole isolation between the two tetramers of interest. Each precursor was isolated with a 1.1 Da window and followed by HCD 30% recorded at 15,000 resolutions in the Orbitrap with the first m/z 130. The monoisotopic mass of UUAGp is 1304.1 Da equivalent to doubly charged m/z 651.07 and UUm^6^AGp has a monoisotopic mass of UUAGp is 1318.2 Da equivalent to doubly charged m/z 658.08. The ratio of UUm^6^AGp over UUAGp was calculated from the average intensities of the respective fragment ions. The MS2 spectra over the targeted precursors for 30 seconds were averaged and the total fragment intensities were calculated and compared. The intensities for UUm^6^AGp fragment ions were divided by the total sum of UUAGp and UUm^6^AGp fragment ions intensities.

#### Telomere DNA-FISH and IF

The cells were fixed with 4% formaldehyde for 10□min, followed by two washes with PBS. Then, cells were permeabilized using 0.25% Triton X-100 in PBS for 10 min, followed by two washes with 0.1% PBS-Tween (PBS-T). Blocking step and RNase A treatment were performed by adding RNase A (100 µg/ml, Sigma-Aldrich) in 3% bovine serum albumin (BSA) for 1 h at RT, followed by 2 washes with PBS-T. The cells were incubated overnight with primary antibody γ-H2AX (1:500, Thermo Fisher Scientific) at 4°C, washed three times 5 min in PBS-T, and subsequently incubated with secondary antibody conjugated with Alexa Fluor 555 (1:800, A32727, Thermo Fisher Scientific) for a 1 h in the dark at room temperature. After incubation with the antibody, cells were washed three times for 5 min in PBS-T. Following IF staining, the cells were refixed using 4% formaldehyde for 10 min. After washing twice with PBS, the slides were dehydrated with ethanol gradient for 3 min and air dried for 5 min at RT. For hybridization, denature slide with probe (1:200, (TAACCC)_7_-Atto488) in hybridization buffer (20 mM Na_2_HPO_4_, pH 7.4, 20 mM Tris, pH 7.4, 60% formamide, 2x SSC and 0.1 µg/ml salmon sperm DNA) for 10 min at 85°C, and incubated with the probe in a humidified chamber for 2 h at RT. Following hybridization, wash twice in washing solution (2x SCC/0.1% Tween) for 15 min each and MQ water twice for 2 min. Prolong Gold with DAPI (Thermo Fisher Scientific) was added to each coverslip and air-dried in the dark before imaging using 60X objectives in EVOS M7000 microscope.

Telomere fluorescence intensity (TFI) was measured using telomere-specific DNA-FISH signals, which are directly proportional to telomere length (37). The analysis involved calculating the ratio of total telomeric signal intensity to DAPI stained nuclear DNA signal on a per-cell basis. The (TAACCC)_7_-Atto488 was utilized to assess the telomeric spots, with their fluorescence intensities and corresponding areas used to determine the TFI.

#### Mouse xenograft experiments

TetO Control and METTL3-KD SK-N-FI and CHLA-90 cells were subcutaneously injected into the right dorsal flank (2 x 10^6^ and 5 × 10^6^ cells, respectively) of 5-week-old female nude mice (Charles River) in 100 µL and 200 µL mixture of (1:3) Matrigel/PBS, (n=4 and 3 per group, respectively). METTL3-KD was induced by adding 2 mg/mL DOX and 2% sucrose to drinking water, after 4-5 days of cell injection. Mice weight was monitored weekly and tumor volume was measured every 2–3 days using a digital caliper and calculated according to the formula Volume (mm^3^) = (w^2^ x l x π)/6 (where w is width, shortest diameter, and l is length, longest diameter). Mice were euthanized when tumors reached 1000 mm^3^ or had weight loss ≥ 10% of initial weight. At the end of the experiment, tumors were collected, weighed, and processed for further analysis. All experiments were carried out as per the standards approved by the Institutional Ethical Committee of Animal Experimentation, Gothenburg, Sweden (ethical permit no 3722/21). METTL3-KD was verified before and after cell injection by Western blot.

#### DNA-RNA immunoprecipitation (DRIP) and DRIP followed by m^6^A RIP (DRImR)

DRIP was performed as described previously (38) with few alterations. In brief, cells were lysed in lysis buffer (1xTE supplemented with 0.5% SDS) for 15 min at RT. Brief RNase A treatment at 37°C for 15 min was done to remove free RNA from the subsequent Proteinase K treatment and genomic DNA isolation by Phenol chloroform isolation. The genomic DNA before S9.6 IP were digested with a cocktail of restriction enzymes (*Bsr*GI, *Eco*RI, *Hind*III, *Ssp*I, and *Xba*I) and RNase III which is shown to remove non-specific enrichment of double-stranded RNA to identify genuine R-loop enriched RNAs. IP reaction was carried out with fragmented genomic DNA that was either pre-treated/non-pre-treated with RNase H to cross-check the specificity of S9.6 IP. Following 9.6 IP we performed extensive DNase I treatment to obtain R-loop associated RNA. For DRImR, DRIP enriched RNA was used as input for m^6^A RIP as described above without fragmentation.

#### Image quantification

Western and slot blots were quantified using ImageLab software (Bio-Rad). Images were quantified using ImageJ software (39). To investigate the potential influence of stochastic cluster overlap, we utilized the Interaction Factor package in ImageJ to randomize the signals (40). DAPI was used to mark the nuclear border and TERRA foci, PLA foci, or colocalization signals including intensity values within the nucleus were obtained using particle analysis. Graphs were prepared, and statistical analysis was performed using GraphPad Prism (version 9.4.1 for Windows, GraphPad Software, San Diego, California, USA, www.graphpad.com) and *p-*values were reported in the figures and the figure legends. Each box-and-whisker plot presents the median value as a line, a box around the lower and higher quartiles, and whiskers extending to the maximum and lowest values.

## Data availability

The data underlying this article are available in NCBI Gene Expression Omnibus (GEO) at https://www.ncbi.nlm.nih.gov/geo/ and can be accessed with the GSE219146 and GSE219212 accession numbers.

## Results

### TERRA RNA is enriched with m^6^A modification in ALT+ cells

To determine whether TERRA RNA undergoes m^6^A modification, we conducted m^6^A RNA immunoprecipitation (m^6^A RIP) using an m^6^A antibody in ALT+ U-2 OS cells, followed by slot blot analysis with a probe targeting telomeric repeats. Our results revealed the enrichment of TERRA, and this m^6^A enrichment was sensitive to RNase A treatment (Figure 1A). As a positive control for m^6^A RIP, we performed m^6^A RIP qPCR with *TUG1* and *HPRT1* served as a negative control (41) (Figure 1A). To investigate the specific location of m^6^A modification within TERRA, we performed m^6^A RIP using both fragmented and non-fragmented RNA. We hypothesized that if m^6^A modifications are present in the telomeric repeat sequences, extensive fragmentation (<150 nt) would have a minimal impact on TERRA m^6^A enrichment when detected with a probe against telomeric repeats in the slot blot assay. Conversely, if major m^6^A modifications are located in non-repetitive regions of TERRA, m^6^A enrichment would be more significantly affected after fragmentation. To assess the specificity of m^6^A RIP in fragmented RNA, we employed primers targeting the m^6^A peak regions of *TUG1* and 300 nucleotides upstream of the m^6^A peak in a qPCR assay to measure the enrichment. We observed that while m^6^A RIP performed on non-fragmented RNA could not differentiate between the enrichment of *TUG1* RNA sequences at the m^6^A peak and those 300 nucleotides apart, fragmented RNA allowed for differentiation (Supplementary Figure S1A). Moreover, the specific enrichment of *TUG1* m^6^A was sensitive to METTL3 knockdown (METTL3-KD) (Supplementary Figure S1A). We detected no major difference in TERRA m^6^A enrichment between fragmented and non-fragmented RNA, this indicated that a major fraction of m^6^A modifications in TERRA is located in the telomeric repeat sequences (Figure 1B, C; Supplementary Figure S1A). The m^6^A enrichment of TERRA was compromised in METTL3-KD cells for both fragmented and non-fragmented RNA (Figure 1C).

**Figure 1:**
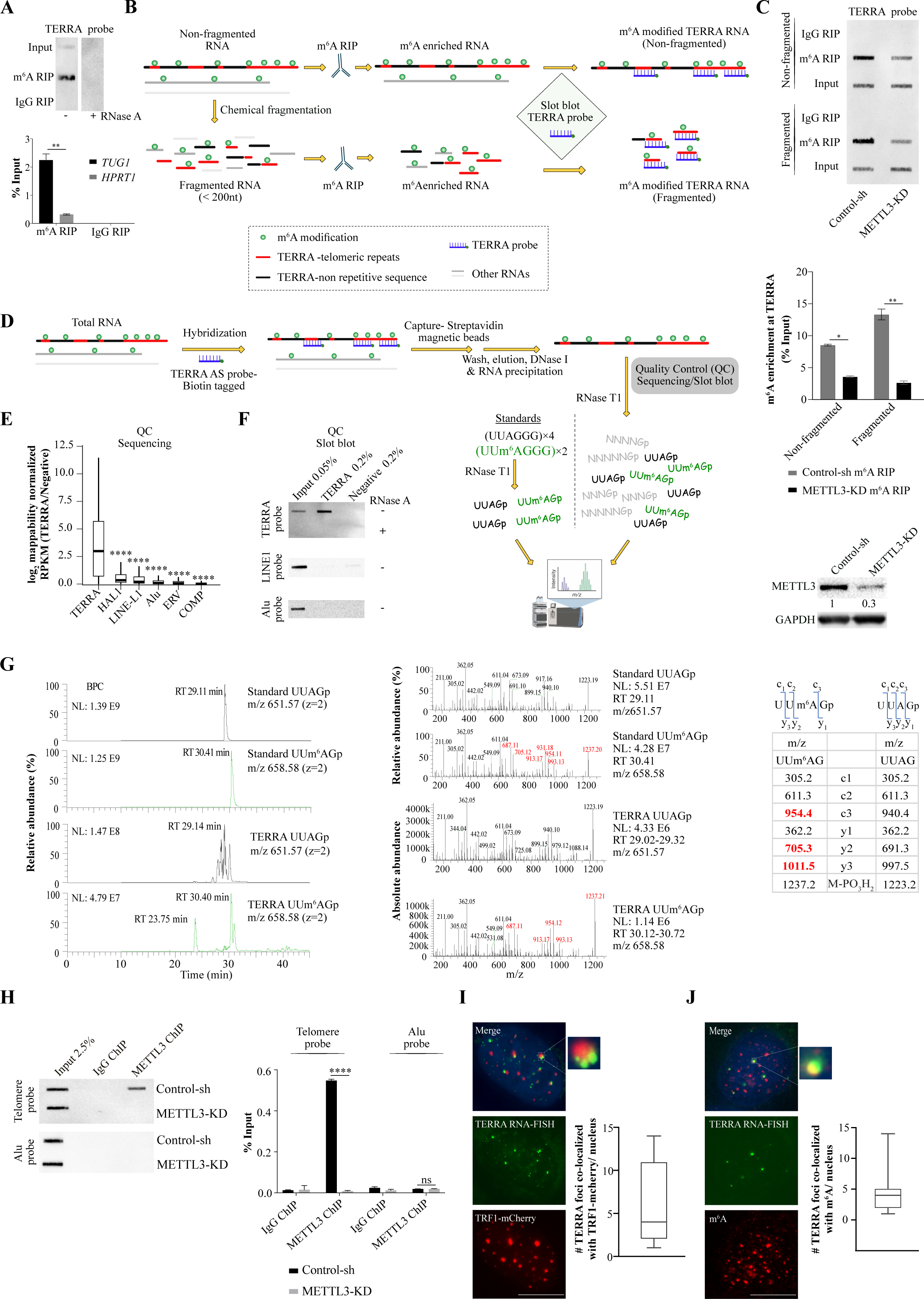
(**A, top panel**) m^6^A RNA Immunoprecipitation (m^6^A RIP) on RNA isolated from U-2 OS cells followed by slot blot, probed with DIG-labeled TERRA probe. RNase A treatment was used as a control. (**Bottom panel**) m^6^A RIP-qPCR validation of samples used in slot blot with *TUG1* and *HPRT1* primers. Data is represented as a percentage of Input. Data are shown as mean ± SD from three biological replicates. Two-way ANOVA with Sidak’s *post hoc* test was used, ** *p <* 0.01, * *p <* 0.05. m^6^A RIP in fragmented and non-fragmented RNA (**B**) Schematic diagram describing key steps of m^6^A RIP performed on either fragmented or non-fragmented RNA isolated from Control-sh or METTL3-KD U-2 OS cells. Keys for the schematic are indicated in the box. (**C, top panel**) m^6^A RIP performed with either fragmented or non-fragmented RNA isolated from Control-sh or METTL3-KD U-2 OS cells, followed by slot blot with DIG-labeled TERRA repeat probe. (**Middle panel)** Quantification of the slot blot. Data are shown as mean ± SD from two biological replicates. Two-way ANOVA with Sidak’s *post hoc* test was used, ** *p <* 0.01, * *p <* 0.05. (**Lower panel**) A representative Western blot showing METTL3- KD, and GAPDH was used as a loading control. The values below indicate the fold change in levels of METTL3. Detection of UUAG tetramer in TERRA RNA by RNA-MS analysis (**D**) Schematic diagram describing key steps of TERRA pulldown using biotin tagged TERRA antisense probe and RNA-MS analysis. Keys for the schematic are same as in (**B**). (**E, F**) Verification of TERRA pulldown by either RNA sequencing or slot blot. (**E**) Boxplot demonstrating log_2_ fold enrichment of mappability normalized RPKM values of TERRA pulldown over negative (luciferase) pulldown at different genomic repeat elements and TERRA repeats. Statistical significance between TERRA and other genomic repeat elements was calculated using the Wilcoxon test, **** *p <* 0.0001. (**F**) Slot blot assay of input, TERRA, and negative pulldown, probed with either TERRA, LINE1, or Alu probe, and RNase A treatment was used as a control. (**G**, **left panel**) Base peak chromatogram (BPC): first and second row show the BPC of Standard UUAGp, isolated m/z 651.07 (doubly charged), and Standard UUm^6^AGp, isolated m/z 658.08 (doubly charged) respectively. Third and fourth rows show the BPC of the TERRA pulldown sample, isolated m/z 651.07 (doubly charged) and isolated m/z 658.08 (doubly charged) respectively. **(Middle panel**) MS2 spectrum: first and second rows show the MS2 spectrum of Standard UUAGp, isolated m/z 651.07, and Standard UUm^6^AGp, isolated m/z 658.08 respectively with ions of mass difference 14 (methylation) starting from m/z 673 and up (highlighted in red). Y-axis represents relative abundance in percentage. Third and fourth rows show the MS2 spectrum of the sample, isolated m/z 651.07 and isolated m/z 658.08 respectively with ions of mass difference 14 (methylation) starting from m/z 673 and up (highlighted in red). Y-axis represents absolute abundance. (**Right panel**) Fragment ion series for the MS2 spectra of UUm^6^AGp and UUAGp with ions of mass difference 14 (methylation) are highlighted in red. (**H**) Chromatin Immunoprecipitation (ChIP) using METTL3 antibody for Control-sh or METTL3-KD U-2 OS cells, followed by slot blot. Blot probed with a DIG-labeled Telomere probe or Alu probe. Quantification of the METTL3 ChIP slot blot represented as percentage input in bar graph. IgG antibody was used as a negative control. Data are shown as mean ± SD from two biological replicates. Two-way ANOVA with Sidak’s *post hoc* test was used, *** *p <* 0.001. (**I**) TERRA foci (green) in U-2 OS cells expressing TRF1-mCherry (red). Box plot shows the number of co-localizations between TRF1-mCherry and TERRA. 52 cells were counted from two independent biological replicates. (**J**) m^6^A IF (red) was performed in U-2 OS along with TERRA foci (green). Box plot shows the number of co-localizations between TERRA and m^6^A. Scale bar is 10 μm. 86 cells were counted from three independent biological replicates.

To further confirm the presence of m^6^A in TERRA repeats, we purified TERRA from total RNA using a biotin-labelled antisense probe specific to TERRA, together with a negative control probe targeting Luciferase RNA (Figure 1D). The specific enrichment of TERRA was verified compared to the negative control using RNA-seq and slot blot assays, and no enrichment was observed for other repeat-containing RNAs such as LINE-1 and Alu (Figure 1E, F). The enrichment of TERRA was also sensitive to RNase A treatment (Figure 1F). Furthermore, we conducted RNA mass spectrometry (RNA-MS) analysis on purified TERRA RNA and synthesized RNA oligo standards containing TERRA repeats with or without m^6^A modification after digestion with ribonuclease T1 (RNase T1) (Figure 1D, G). Using the RNA-MS we confirmed the presence of UUAGp and UUm^6^AGp fragments in our oligo standards using higher-energy collisional dissociation (HCD) tandem MS (MS2) data (Figure 1G). Leveraging this MS2 information from the standards, we identified UUAGp tetramers within the complex mixture of other RNA fragments generated by RNase T1 cleavage. HCD MS2 analysis further confirmed m^6^A modification in the UUAGp tetramer (UUm^6^AGp) with approximately 21% of UUAGp fragments being m^6^A modified (Figure 1G).

To validate whether METTL3 could m^6^A modify the telomeric repeat sequences in TERRA, we performed an *in vitro* m^6^A methyltransferase assay using a purified METTL3/METTL14 enzyme complex. As substrates, we utilized synthesized TERRA oligo (UUAGGG)_3_ or DRACH oligo (GGACU)_3_, followed by treatment with Nuclease P1 and liquid chromatography-tandem mass spectrometry (LC-MS/MS) analysis to detect m^6^A/A (Supplementary Figure S1B). We conducted parallel digestion of the synthesized standard oligos with or without m^6^A modification using Nuclease P1 and included them in the LC-MS/MS analysis (Supplementary Figure S1B). Our results demonstrated *in vitro* m^6^A modification of both the TERRA and DRACH oligos by the METTL3/METTL14 complex (Supplementary Figure S1B). Although the efficiency of m^6^A modification in the TERRA oligo was lower compared to the canonical "DRACH" oligo, as indicated by the m^6^A/A ratio (Supplementary Figure S1B), our data support the notion that the METTL3/METTL14 complex can modify telomeric repeats in TERRA RNA.

Immunofluorescence (IF) analysis of U-2 OS cells revealed that METTL3 exhibits nuclear localization with distinct punctate structures, consistent with earlier observations (42), and these METTL3 puncta frequently co-localize with telomeres (detected by TRF1-mCherry). The observed degree of co-localization between METTL3 and TRF1-mCherry was much higher than stochastic overlap (Supplementary Figure S1C). To determine whether METTL3 is localized to telomeres in ALT+ U-2 OS cells, we performed Chromatin Immunoprecipitation (ChIP) and observed that METTL3 is recruited to ALT telomeres, with this recruitment being compromised in METTL3-KD cells (Figure 1H). Additionally, a proximity ligation assay (PLA) using TRF2 in combination with either METTL3 or METTL14 revealed an overlap of METTL3/TRF2 and METTL14/TRF2 PLA signals with telomeres detected by TRF1-mCherry. The negative control PLA did not yield any TRF1-mCherry overlapping signal (Supplementary Figure S1D). Collectively, our data suggest that METTL3 is the m^6^A methyltransferase for TERRA.

We conducted RNA *in situ* hybridization (RNA-FISH) to validate the localization of TERRA at telomeres and found that TERRA foci frequently overlapped with the TRF1-mCherry signal (Figure 1I). We confirmed the specificity of TERRA foci using locked-nucleic acid (LNA)-mediated knockdown of TERRA (Supplementary Figure S1E). Using a modified RNA-FISH protocol that reduces background staining in m^6^A IF (43), we detected m^6^A foci in the nucleus of control U-2 OS cells, which were reduced following METTL3-KD (Supplementary Figure S1F). We also observed a frequent overlap between m^6^A and TERRA foci, further suggesting the presence of m^6^A modification on TERRA (Figure 1J). We performed TERRA RNA-FISH combined with m^6^A IF on metaphase chromosomes and observed that TERRA foci at the telomere ends frequently overlapped with m^6^A signals (Supplementary Figure S1G).

To further investigate TERRA expression and m^6^A modification, we performed short-read (Illumina) and long-read (ONT: Oxford Nanopore Technologies) RNA sequencing (RNA-seq) and m^6^A RIP sequencing (m^6^A RIP-seq) in Control-sh and METTL3-KD U-2 OS cells (Figure 2A). For accurate mapping of sequencing reads to telomere ends, we utilized the latest telomere-to-telomere (T2T) human genome assembly (44), specifically focusing on subtelomeric regions in our RNA-seq data analysis. By combining strand-specific Illumina short reads and ONT long reads, we identified both transcriptionally active and inactive subtelomeres from both the p and q arms of chromosomes (Figure 2B). Since TERRA RNAs are transcribed toward the telomere ends, the strand-specific Illumina short reads were mainly derived from one of the strands, depending on the chromosome arms (p and q) from which they originated (Figure 2B; Supplementary Figure S2A). We remapped available CTCF ChIP-seq data to the T2T assembly and observed that both proximal and distal CpG islands located over subtelomeres are flanked by CTCF sites, consistent with previous reports (Figure 2B) (45). We also found occupancy of RNA Pol II at actively TERRA transcribing chromosome ends using available RNA Pol II ChIP-seq data (Figure 2B). Considering the repetitive nature of chromosome ends, we employed mappability normalization to the RNA-seq data. We used mappability-normalized short and long-read RNA-seq data to define high and low TERRA-expressing subtelomeres based on the reads covering 30 kb (distance from the last consecutive telomeric repeats) of subtelomeric regions, including both proximal and distal CpG islands. We consistently observed activity of high TERRA-expressing subtelomeres in both short and long-read RNA-seq datasets (Figure 2C; Supplementary Figure S2B, C). Although METTL3 showed a trend of subtelomere-specific decrease in TERRA expression from some chromosome ends, the overall difference in TERRA levels between METTL3-KD and control cells was not statistically significant (Supplementary Figure S2B, C). This is consistent with the minor (∼15%) decrease in TERRA expression observed in METTL3-KD cells compared to Control-sh cells using the slot blot assay (Supplementary Figure S2D).

**Figure 2:**
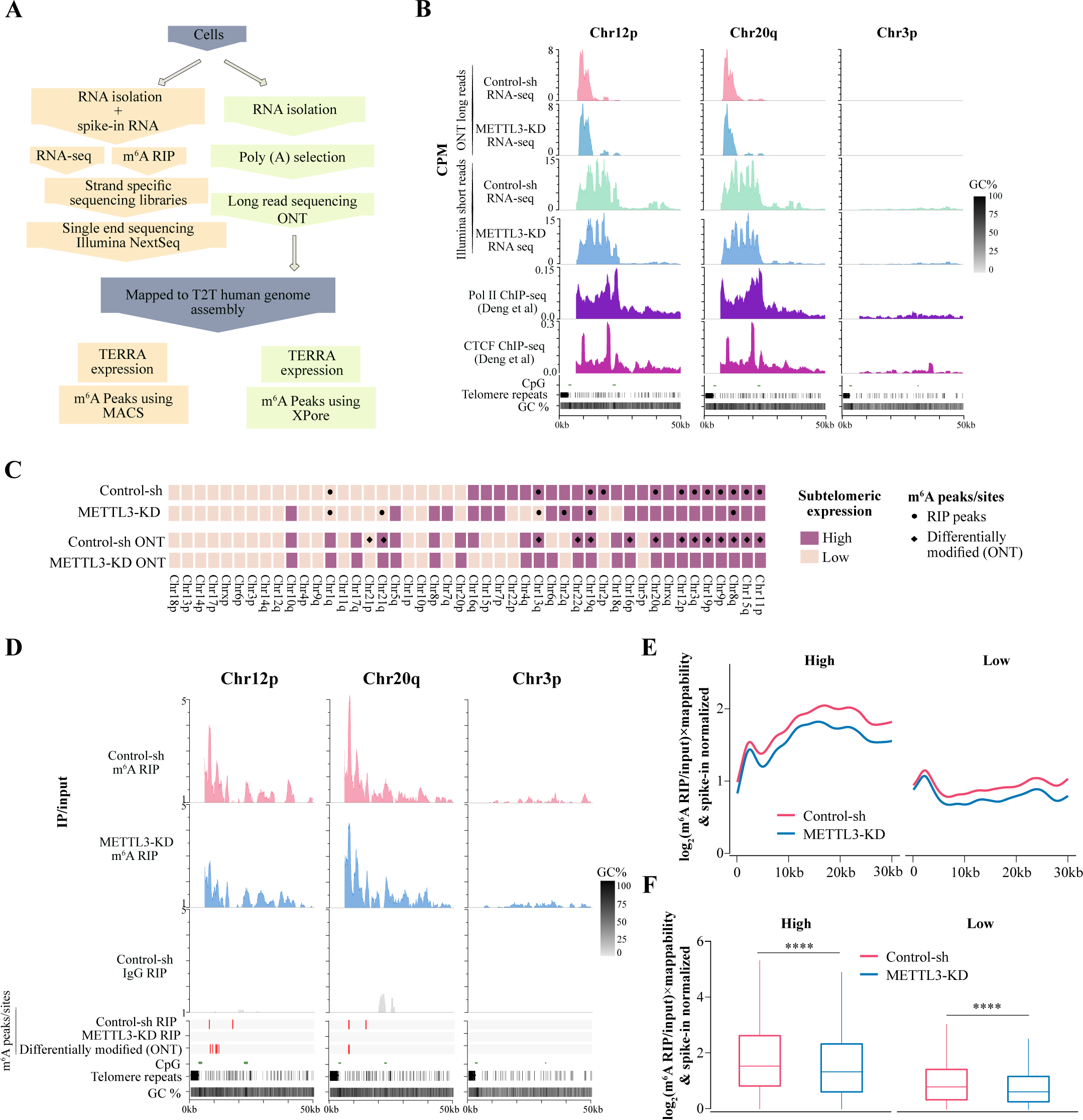
(**A**) Flow chart depicting the major step of m^6^A RIP-seq along with RNA-seq performed either on Illumina platform or on Oxford Nanopore Technologies (ONT), to generate short and long RNA-seq reads respectively. (**B**) Browser screenshot of 50 kb region around the telomere ends in T2T assembly, showing TERRA RNA expression (CPM) from two active chromosome ends (Chr12p, Chr20q) and one less active chromosome end (Chr3p) using ONT long read and Illumina short-read RNA-seq. RNA Pol II (purple) and CTCF (pink) enrichment at 50 kb region around the telomere ends from publicly available RNA Pol II and CTCF ChIP-seq data which were re-analyzed with T2T human genome assembly. CpG islands are marked with green bars and the telomeric repeats in this region are marked with black bars. (**C**) Heatmap summarizing RNA-seq and m^6^A RIP-seq data. Chromosomes are sorted based on high (dark violet-pink) and low (light violet-pink) subtelomeric TERRA transcription. Black dots denote m^6^A RIP-seq peaks and black diamonds denote the differentially modified m^6^A sites detected by xPore from ONT long reads at subtelomeres. (**D**) Genome browser screenshots showing IP/input ratio tracks for m^6^A RIP (Control-sh and METTL3-KD) or IgG RIP (Control-sh) at two active chromosome ends (Chr12p, Chr20q) and one inactive chromosome end (Chr3p). Red bars mark m^6^A RIP peaks identified using MACS (for short reads) or the differential m^6^A sites identified using xPore (for long reads) for the samples as denoted. (**E**) The m^6^A enrichment profiles across all subtelomeres in T2T assembly in Control-sh and METTL3-KD. The m^6^A RIP/input signals were normalized to spike-in and to the mappability likelihood of each chromosome ends. Subtelomeres (high and low TERRA transcription) were classified according to the normalized median expression shown. (**F**) Box plot of spike-in, mappability-normalized m^6^A RIP/input enrichment profiles of all chromosome ends. Statistical significance was calculated using the Wilcoxon test, **** *p <* 0.0001.

After we had established the subtelomeric TERRA expression profile in U-2 OS cells, we investigated whether telomeric repeats and subtelomeric sequences are enriched with m^6^A modifications. We made certain modifications to the m^6^A RIP-seq protocol to profile m^6^A with a smaller amount of RNA (31) while preserving the strand information of the RNA-seq data. We also used bacterial RNA as a spike-in control in our m^6^A RIP-seq protocol, allowing us to accurately normalize the data (Figure 2A). The m^6^A distribution from RIP-seq performed on U-2 OS RNA was found to be remarkably similar as described earlier (5), with major m^6^A peaks observed at the 3’ UTR and enrichment of the "DRACH" motif (Supplementary Figure S2E, F). In the m^6^A RIP-seq data from METTL3-KD cells, we observed an overall decrease in m^6^A peaks compared to the control, indicating the specificity of the m^6^A RIP-seq (Supplementary Figure S2G).

We quantified the amount of telomeric repeat reads present in the m^6^A RIP-seq data and found higher enrichment of telomeric repeat reads in Control-sh cells compared to METTL3- KD and IgG control (Supplementary Figure S2H). Furthermore, we observed that TERRA from several subtelomeric regions are enriched with m^6^A modifications (m^6^A/input ratio), and this enrichment was affected in METTL3-KD cells (Figure 2D). We detected m^6^A peaks in the subtelomeres, and there was an overall decrease in subtelomere-associated peaks in METTL3-KD cells (Figure 2C). To visualize m^6^A enrichment across several active and inactive subtelomeres together, we plotted mappability and spike-in normalized m^6^A RIP-seq signal over input (log_2_ m^6^A/input) obtained from Control-sh and METTL3-KD cells over the 30 kb subtelomeric regions as metagene plots. We observed m^6^A enrichment over subtelomeres in Control-sh cells, and this enrichment was significantly decreased after METTL3-KD (Figure 2E, F).

A recent report suggests that "DRACH"-like sequences in TERRA can also contribute to m^6^A modification (46). We observed the presence of the DRACH sequence in the subtelomeric m^6^A peaks, and m^6^A enrichment (log_2_ m^6^A/input) over the DRACH sequence was modestly but significantly decreased in METTL3-KD compared to Control-sh (Supplementary Figure S2I).

We also used long-read RNA-seq data from Control-sh and METTL3-KD cells to identify differentially modified m^6^A sites. Among the 27,125 significantly modified NNANN sites identified by xPore (47) genome-wide, 26,952 could be successfully translated to annotated gene coordinates. The differentially modified m^6^A sites showed a similar 3’ UTR enrichment as observed in the m6A RIP-seq peaks, with considerable overlap between the two m^6^A detection methods (Supplementary Figure S2E, J). We also observed enrichment of the "DRACH"-like motif in the long-read-detected m^6^A sites (Supplementary Figure 2F). Next, we focused on the non-repeated part of subtelomeres to search for possible m^6^A sites, considering inherent base calling errors in Nanopore long-read sequencing at telomeric repeats (48). We performed a transcriptome assembly based on the long reads mapped to the T2T reference genome and identified 92 NNANN sites. The NNANN sites that were differentially m^6^A modified between Control-sh and METTL3-KD, were located within the last 30 kb regions of subtelomere ends (as defined above), with 7 of them having a DRACH motif (Supplementary Figure S2K). Subtelomeres with high expression of TERRA also contained more differentially detected m^6^A sites in ONT reads compared to low-expressed ones, and these sites frequently overlapped with m^6^A RIP-seq detected peaks (Figure 2C, D).

In summary, our findings demonstrate that TERRA undergoes m^6^A modification in both telomeric repeats and subtelomeric non-repeated regions, which frequently contain a "DRACH"-like motif.

### METTL3 mediated m^6^A modification of TERRA RNA is required for telomere maintenance in ALT+ cells

We next investigated the importance of m^6^A modification on TERRA RNA. As evident from the results above, TERRA forms foci over telomeres in U-2 OS cells, and these foci are sensitive to TERRA KD (Figure 1I; Supplementary Figure S1E). We first tested the role of m^6^A modification on TERRA foci formation by performing RNA-FISH on U-2 OS cells with transient KD of METTL3 using siRNA. We observed that TERRA foci formation was compromised in the METTL3 siRNA-treated cells compared to the control (Supplementary Figure S3A). Since we observed co-localization of METTL3/METTL14 over ALT telomeres (Supplementary Figure S1D), we also tested the effect of siRNA-mediated depletion of METTL14 on TERRA foci formation. Similar to METTL3, METTL14 depletion led to compromised TERRA foci (Supplementary Figure S3B). METTL3 siRNA-treated U-2 OS cells showed reduced proliferation (Supplementary Figure S3C), which is consistent with previous observations (46). The loss of TERRA foci was further validated in stable KD of METTL3 in U-2 OS cells (Figure 3A). TERRA is known to contribute to the maintenance of telomeres; therefore, we sought to address the significance of TERRA m^6^A modification in telomere maintenance. To investigate this, we performed IF staining with γ-H2AX, a DNA damage response marker, and TRF2 (to mark the telomeres) in control and METTL3-KD cells. METTL3-KD cells showed an increased level of γ-H2AX, consistent with the role of METTL3-mediated m^6^A modification in DNA repair (49), while the number of TRF2 foci did not significantly change following METTL3-KD (Supplementary Figure S3D). METTL3-KD resulted in γ-H2AX accumulation at telomeres, indicating telomere damage in these cells (Figure 3B). Despite the overall increase in γ-H2AX in METTL3-KD cells, the observed overlap between γ-H2AX and TRF2 was higher than expected by chance alone (Supplementary Figure S3E). Using ChIP-qPCR, we further confirmed increased enrichment of γ-H2AX at telomeres in METTL3-KD cells (Supplementary Figure S3F). We also observed higher γ-H2AX deposition at telomeres in TERRA-depleted cells (Supplementary Figure S3G), suggesting that TERRA is required for telomere maintenance in ALT cells (49,50). We hypothesized that if we create telomere DNA damage exogenously using bleomycin, a well-characterized DNA damaging agent, in U-2 OS cells (51), it will allow us to dissect the role of TERRA m^6^A modification in such repair processes. U-2 OS cells were treated with bleomycin for 4 h followed by recovery in bleomycin-free media for 20 h. Bleomycin treatment led to an overall increase in γ-H2AX levels in METTL3-KD cells after both 4 h treatment and recovery phases (Supplementary Figure S3H). In control U-2 OS cells, the accumulation of γ-H2AX at telomeres ceased during the post-bleomycin recovery phase. However, in METTL3-KD cells, the accumulation of γ-H2AX at telomeres did not cease but rather continued to increase during the post-bleomycin recovery (Figure 3C). We also detected increased γ-H2AX accumulation in METTL3-KD cells over telomeres during bleomycin recovery using ChIP-qPCR (Supplementary Figure S3I). Furthermore, we observed a higher number of TERRA and m^6^A foci during bleomycin recovery compared to untreated cells (Supplementary Figure S3J). The overlap between TERRA and m^6^A also increased significantly during bleomycin recovery (Figure 3D). These data collectively suggest a role for METTL3-mediated TERRA m^6^A modification in telomere damage repair.

**Figure 3:**
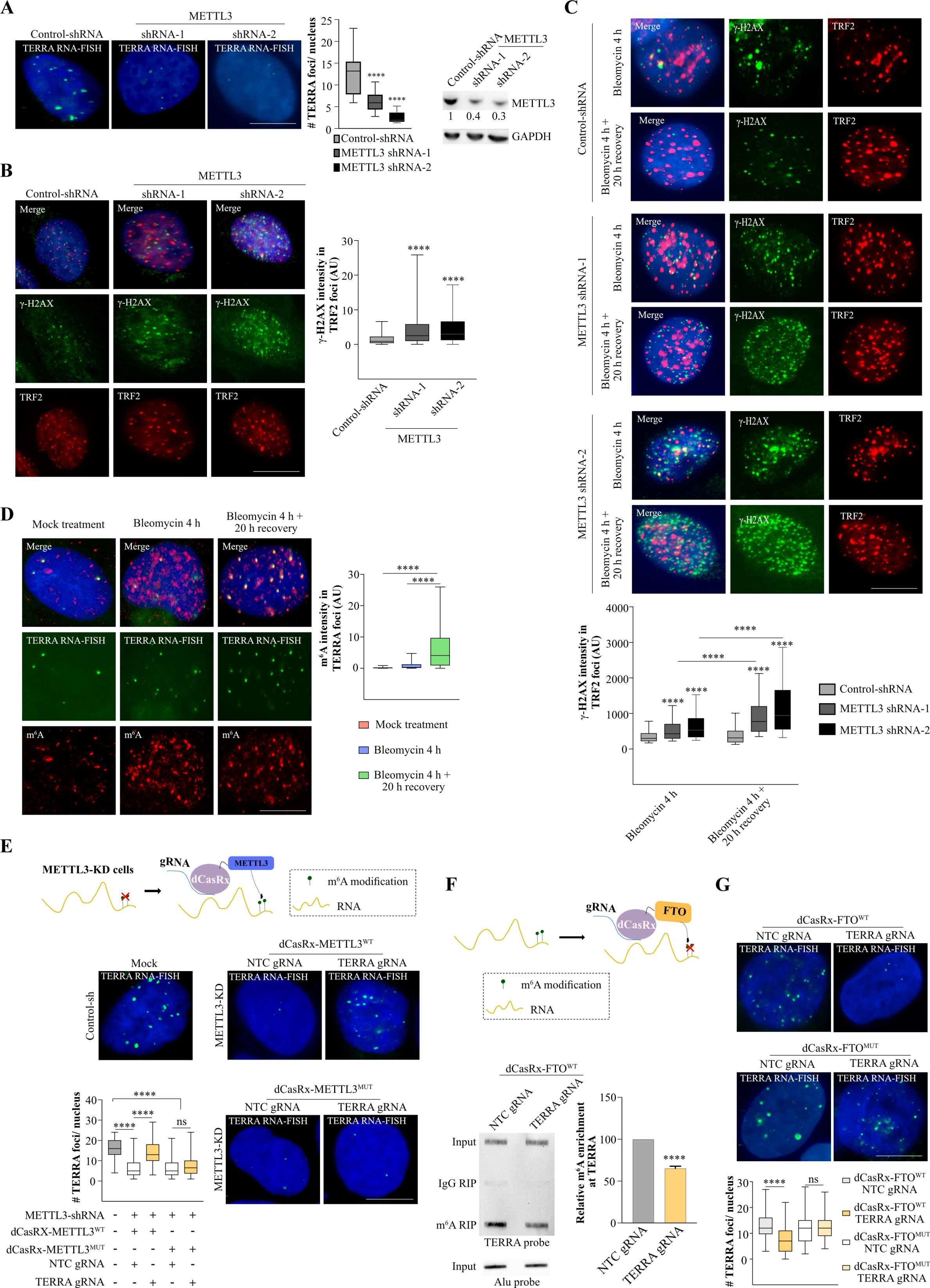
(**A, left panel**) TERRA foci (green) in U-2 OS cells with stable KD of METTL3. (**Middle panel**) Box plot shows the number of TERRA foci per nucleus. Dunnett’s multiple comparisons test was used, **** *p <* 0.0001. At least 100 cells were counted from three independent biological replicates. (**Right panel**) Western blot showing METTL3 level in stable shRNA-depleted U-2 OS cells. GAPDH was used as a loading control. The values below indicate the fold change in levels of METTL3. (**B**) Localization of the γ-H2AX (green) over telomere in Control-sh and METTL3-KD cells. Telomere is detected by TRF2 staining (red). Box plot shows the γ-H2AX intensity in TRF2 foci (Arbitrary units, AU). Dunnett’s multiple comparisons test was used, **** *p <* 0.0001. At least 70 cells were counted from three independent biological replicates. (**C**) Localization of γ-H2AX (green) over telomere (marked by TRF2) in bleomycin-treated Control and METTL3-KD cells 4 h post-treatment and followed by 20 h of recovery without bleomycin. Box plot shows the γ-H2AX intensity in TRF2 foci (Arbitrary units, AU). Two-way ANOVA with Tukey’s *post hoc* test was used, **** *p <* 0.0001. At least 70 cells were counted from three independent biological replicates. (**D**) m^6^A IF (red) along with TERRA RNA-FISH (green) was performed in U-2 OS cells 4 h post-treatment and followed by 20 h of recovery without bleomycin. Box plot shows the m^6^A intensity in TERRA foci (Arbitrary units, AU). One-way ANOVA with Tukey’s *post hoc* test was used, **** *p <* 0.0001. At least 70 cells were counted from three independent biological replicates. (**E**) Cartoon demonstrating recruitment of dCasRx-METTL3 to target RNA. TERRA RNA-FISH detecting TERRA foci (green) was performed on METTL3-KD U-2 OS cells expressing dCasRx-METTL3^WT^ (wild-type-shRNA resistant) / dCasRx-METTL3^MUT^ (catalytically dead APPA mutant-shRNA resistant) with either NTC (non-target control) or TERRA guide RNA. Control-sh cells were used as a positive control for TERRA RNA-FISH. Box plot shows the quantification of the number of TERRA foci per nucleus in the conditions as indicated. One-way ANOVA with Tukey’s *post hoc* test was used, **** *p <* 0.0001. ns - nonsignificant *p* > 0.05. At least 100 cells were counted from three independent biological replicates. (**F**) Illustration demonstrating recruitment of dCasRx-FTO to target RNA. m^6^A RIP followed by TERRA slot blot on RNA isolated from U-2 OS cells expressing dCasRx-FTO^WT^ with either NTC or TERRA guide RNA. IgG antibody was used as a negative control for the RIP experiment. Input samples were probed with Alu probe to serve as a loading control. Bar graph shows the quantification of slot blot. Data are shown as mean ± SD from two biological replicates. Unpaired *t*-test was used, **** *p <* 0.0001 (**G**) TERRA RNA-FISH detecting TERRA foci (green) was performed on U-2 OS cells expressing dCasRx-FTO^WT^ (wild-type) / dCasRx-FTO^MUT^ (catalytically dead-H231A and D233A mutant) with either NTC or TERRA guide RNA. Box plot shows the quantification of TERRA foci per nucleus. At least 70 cells were counted from three independent biological replicates. One-way ANOVA with Tukey’s *post hoc* test was used, **** *p <* 0.0001. ns - nonsignificant *p* > 0.05.

METTL3-depleted cells showed compromised ALT activity as measured by the C-Circle assay (Supplementary Figure S3K). Long-term culture of METTL3-KD cells (5 weeks) showed signs of decreased telomere content and frequent detection of telomere-free ends in metaphase spreads, further suggesting a role for METTL3-mediated m^6^A modification in telomere maintenance in ALT cells (Supplementary Figure S3L, M).

To further elucidate the direct role of METTL3-mediated m^6^A modification on TERRA, we used two approaches. Firstly, we recruited METTL3 to TERRA using the RNA-guided CRISPR/dCasRx system (52). Secondly, we specifically removed m^6^A modification from TERRA by recruiting the m^6^A demethylase FTO using CRISPR/dCasRx. To investigate the direct role of METTL3, we recruited shRNA-resistant wild-type METTL3 (METTL3^WT^) or catalytically inactive APPA mutant METTL3 (METTL3^MUT^) (53), fused with dCasRx, in the endogenous METTL3-KD cells (Figure 3E; Supplementary Figure S3N). METTL3-KD cells showed reduced TERRA foci compared to control cells, as described above (Figure 3A), but we were able to recover TERRA foci formation by recruiting shRNA-resistant METTL3^WT^ at TERRA (Figure 3E). Recruitment of METTL3^MUT^ could not recover the loss of TERRA foci phenotype in METTL3-KD cells, suggesting that the enzymatic activity of METTL3 is essential for TERRA foci formation (Figure 3E; Supplementary Figure S3N).

To test the effect of specific removal of m^6^A from TERRA, we fused wild-type (WT) or mutant (MUT: catalytically inactive; H231A and D233A) m^6^A demethylase FTO with dCasRx and recruited them to TERRA in U-2 OS cells (54) (Figure 3F; Supplementary Figure S3O). We verified the removal of m^6^A from TERRA by performing m^6^A RIP followed by TERRA slot blot. Recruiting FTO^WT^ to TERRA resulted in lower enrichment of m^6^A compared to the non-template control (NTC) (Figure 3F). Recruitment of FTO^WT^ did not result in any change in TERRA RNA level (Supplementary Figure S3P). Next, we evaluated the effect of this specific removal of m^6^A on the ability of TERRA foci formation by performing TERRA RNA-FISH. We observed a loss of TERRA foci in cells where FTO^WT^ was recruited at TERRA (Figure 3G). Recruitment of FTO^MUT^ at TERRA did not alter TERRA foci in U-2 OS cells, suggesting that the loss of TERRA foci is dependent on the m^6^A demethylase activity of FTO (Figure 3G; Supplementary Figure S3O). We tested the specificity of the TERRA guide RNA by deploying catalytically active CasRx and found that TERRA RNA was downregulated. CasRx-mediated TERRA downregulated cells showed a reduced level of C-Circles suggesting a direct role of TERRA in ALT activity (Supplementary Figure S3Q, R).

Given the growing importance of METTL3-mediated m^6^A modification in diseases, two independent studies have developed METTL3 inhibitors (55,56). Notably, Yankova et al. extensively tested a small molecule METTL3 inhibitor in various AML mouse models and concluded that METTL3 inhibition is a potential therapeutic strategy in AML (55). We wanted to test the effect of these available METTL3 inhibitors on TERRA foci formation. Considering the TERRA half-life obtained from the ActD data (Supplementary Figure S3S), we treated U-2 OS cells with two METTL3 inhibitors (STM2457 and UZH1A) for 6 h, which would result in compromised deposition of m^6^A in newly formed TERRA molecules. We observed a pronounced reduction in TERRA foci following 6 h of METTL3 inhibition compared to the DMSO control (Supplementary Figure S3T). Treatment with an inefficient METTL3 inhibitor (UZH1B) (56) had a less pronounced effect on TERRA foci (Supplementary Figure S3T). METTL3 inhibition with STM2457 showed the highest effect on TERRA foci without affecting TERRA levels (Supplementary Figure S3U). STM2457 treatment did not alter the cell cycle profile of the U-2 OS cells, consistent with earlier observations (Supplementary Figure S3V) (50).

Collectively, through the modulation of m^6^A using either the dCasRx-METTL3/FTO system or METTL3 inhibitor treatment, we have unraveled the critical role of m^6^A modification in the functionality of TERRA. Our data suggest that METTL3-mediated m^6^A modification of TERRA is vital for TERRA foci formation, and m^6^A modification in TERRA RNA is required for the repair of damaged telomeres in ALT+ cells.

### hnRNPA2B1 binds TERRA RNA in m^6^A-dependent manner and is required for TERRA function

Our results so far suggest that METTL3-mediated m^6^A modification of TERRA RNA is required for TERRA foci formation. Next, we aimed to evaluate if m^6^A reader proteins are involved in this process. We conducted a small siRNA screen to deplete known m^6^A reader proteins, including YTHDC1, hnRNPA2B1, RBMX (nuclear m^6^A readers), and YTHDF2 (cytoplasmic reader) (Figure 4A; Supplementary Figure S4A). Among these, hnRNPA2B1 and YTHDC1 have previously been shown to interact with TERRA (46,57). We show that KD of hnRNPA2B1 had a major effect on TERRA foci formation compared to the other m^6^A reader proteins (Figure 4A). Although we observed a loss of TERRA foci in hnRNPA2B1-depleted cells, the TERRA levels remained unchanged (Supplementary Figure S4B). We also validated the interaction between hnRNPA2B1 and TERRA in the nuclear fraction of U-2 OS cells using RNA RIP followed by slot blot (Figure 4B).

**Figure 4:**
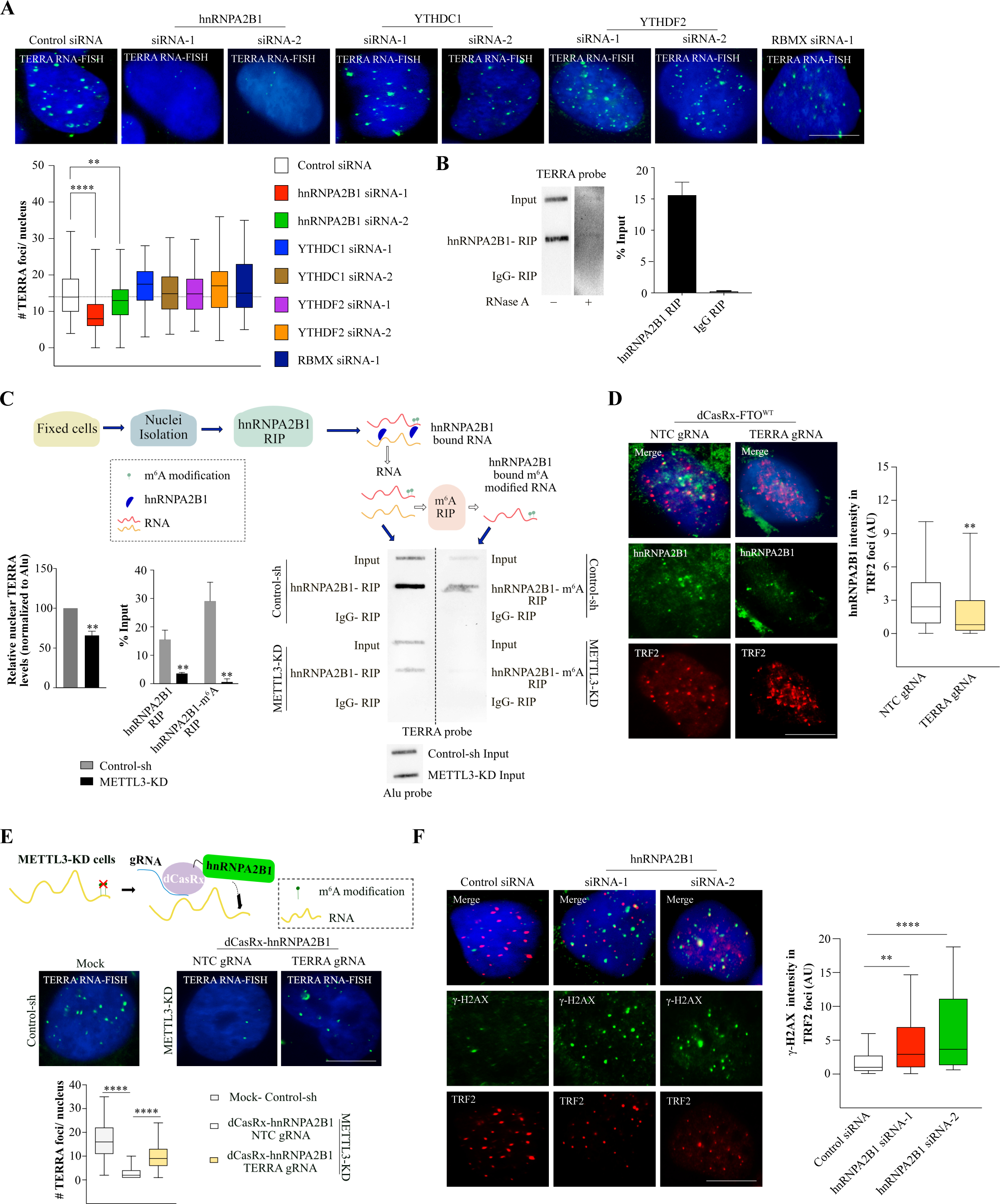
(**A**) TERRA foci (green) in U-2 OS cells with siRNA-mediated transient KD of m^6^A reader proteins as denoted. Box plot shows the number of TERRA foci per nucleus. At least 100 cells were counted from three independent biological replicates. Dunnett’s multiple comparisons test was used ** *p <* 0.01; **** *p <* 0.0001. (**B**) hnRNPA2B1 RIP followed by TERRA slot blot demonstrates the interaction of hnRNPA2B1 with TERRA. Quantification of the hnRNPA2B1 RIP slot blot represented as percentage input in bar graph. IgG RIP and RNase A treatment were used as control. Data are shown as mean ± SD from two biological replicates. (**C**) Scheme and results of the re-RIP (hnRNPA2B1-m^6^A) experiment in Control-sh or METTL3-KD U-2 OS cells. First RIP was performed with hnRNPA2B1/IgG antibody, the precipitated RNA was further subjected to RIP with m^6^A antibody. The RNA after hnRNPA2B1 RIP and re-RIP were blotted and probed with the TERRA probe. Input samples were probed with Alu probe to serve as a loading control. RIP with IgG served as a negative control. Bar graph shows the quantification of TERRA in the nuclear input (Control-sh Vs. METTL3-KD) normalized to Alu (loading control) and for RIP/re-RIP experiment presented as percentage of input. Data are shown as mean ± SD from two biological replicates. Unpaired *t*-test was used, ** *p <* 0.01. (**D**) IF with hnRNPA2B1(green) and telomeres (TRF2, red) in U-2 OS cells recruited with dCasRx-FTO^WT^ at TERRA or NTC. Box plot shows the hnRNPA2B1 intensity in TRF2 foci (Arbitrary units, AU). At least 70 cells were counted from two independent biological replicates. Unpaired *t*-test was used, ** *p <* 0.01. (**E**) Cartoon demonstrating recruitment of dCasRx-hnRNPA2B1 to target RNA in METTL3-KD U-2 OS cells. TERRA RNA-FISH detecting TERRA foci (green) was performed on METTL3-KD U-2 OS cells expressing dCasRx-hnRNPA2B1 with either NTC or TERRA guide RNA. Control-sh cells were used as a positive control for TERRA RNA-FISH. Box plot shows the quantification of TERRA foci per nucleus in the conditions as indicated. At least 100 cells were counted from three independent biological replicates. One-way ANOVA with Tukey’s *post hoc* test was used, **** *p <* 0.0001. (**F**) γ-H2AX (green) accumulation at telomeres (TRF2, red) in U-2 OS cells with siRNA-mediated KD of hnRNPA2B1. Box plot shows the γ-H2AX intensity in TRF2 foci. At least 55 cells were counted from two independent biological replicates. Dunnett’s multiple comparisons test was used, ** *p <* 0.01, **** *p <* 0.0001. Scale bar is 10 μm.

Next, we investigated how changes in TERRA m^6^A levels contribute to hnRNPA2B1 interaction. We performed hnRNPA2B1 RIP in Control-sh and METTL3-KD cells, followed by TERRA slot blot. We observed reduced hnRNPA2B1 interaction with TERRA in METTL3-KD cells, suggesting a role for m^6^A modification in TERRA and hnRNPA2B1 interaction (Figure 4C). To further confirm if hnRNPA2B1-enriched TERRA was modified by m^6^A, we performed m^6^A RIP using hnRNPA2B1-bound RNA and observed that hnRNPA2B1-interacting TERRA was indeed modified with m^6^A (Figure 4C). Consistent with our RIP experiment, purified hnRNPA2B1 efficiently interacted with TERRA derived from control U-2 OS cells, but with reduced efficiency with TERRA derived from METTL3- KD cells (Supplementary Figure S4C). Finally, when we specifically demethylated TERRA using the dCasRx-FTO^WT^, we observed reduced occupancy of hnRNPA2B1 at the telomere (Figure 4D).

It is important to note that the nuclear input from METTL3-KD cells in the hnRNPA2B1 RIP experiment showed a reduced level of TERRA (36% decrease compared to control) (Figure 4C), which was greater than the decrease observed at the level of total RNA (15% decrease in METTL3-KD) (Supplementary Figure S2D). This indicates that in METTL3-KD cells, a fraction of TERRA might be mislocalized to the cytoplasm in the absence of TERRA foci formation over the telomere. Fractionation of nuclear and cytoplasmic RNA showed that both METTL3 and hnRNPA2B1-depleted cells had a higher fraction of TERRA present in the cytoplasm compared to their respective controls, although the distribution of nuclear and cytoplasmic RNA markers was not affected (Supplementary Figure S4D).

To further verify the importance of hnRNPA2B1 in TERRA foci formation, we utilized the dCasRx system to guide hnRNPA2B1 to TERRA RNA in METTL3-KD cells. Our rationale was that if m^6^A-dependent recruitment of hnRNPA2B1 plays a critical role in TERRA foci formation, then in METTL3-KD cells lacking TERRA m^6^A, dCasRx-mediated guidance of hnRNPA2B1 to TERRA should rescue the loss of the TERRA foci phenotype. Indeed, we were able to recover TERRA foci in METTL3-KD cells when hnRNPA2B1 was guided to TERRA, suggesting that forceful recruitment of hnRNPA2B1 could bypass the requirement for m^6^A in TERRA (Figure 4E; Supplementary Figure S4E).

To determine if depletion of hnRNPA2B1 mimicked the increased DNA damage phenotype observed in METTL3-KD cells at telomeres, we performed IF for γ-H2AX and TRF2 in U-2 OS cells. Depletion of hnRNPA2B1 resulted in an increased accumulation of γ-H2AX at telomeres (Figure 4F). Additionally, depletion of hnRNPA2B1 led to a reduced level of C-Circles in U-2 OS cells, indicating compromised ALT activity (Supplementary Figure S4F). Our results demonstrate that the binding of hnRNPA2B1 and m^6^A cooperate for TERRA-dependent maintenance of telomeres in ALT+ cells.

### TERRA m^6^A modification promotes R-loop formation

Recent studies have reported the presence of m^6^A modification in the R-loop, which may contribute to R-loop formation (12,13). Additionally, R-loop formation by TERRA is critical for the targeting of TERRA to the telomere (20). In our study, we observed that the lack of m^6^A modification in TERRA RNA leads to a decrease in TERRA foci, which could be attributed to compromised telomere targeting of TERRA. We hypothesized that the decrease in TERRA R-loop formation in the absence of m^6^A may result in inefficient targeting of TERRA to the telomere. To investigate this, we modified the published DNA-RNA immunoprecipitation (DRIP) protocol, aiming to accurately measure TERRA present in R-loop structures and identify m^6^A-modified TERRA in the R-loop context (Figure 5A).

**Figure 5:**
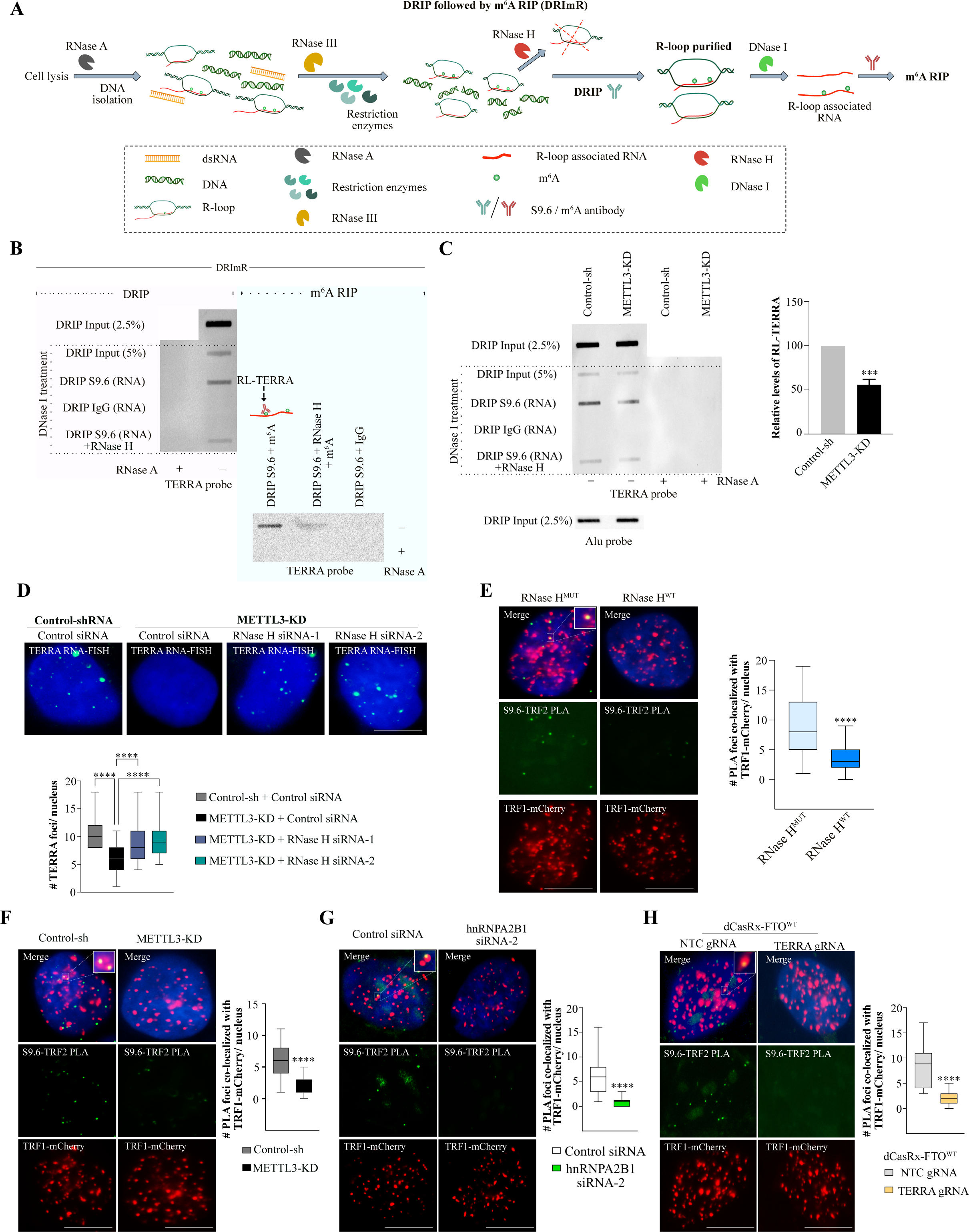
(**A**) Schematic diagram describing the DRIP and DRImR method to purify R-loop bound RNA and R-loop bound m^6^A modified RNA respectively. (**B**) Blot on the left is from RNA purified in the DRIP experiment and the blot on the right is from the DRImR experiment performed on U-2 OS cells. Experiments performed with IgG antibody or nucleic acid pre-treated with RNase H served as a control. RNase A treatment was done to affirm the presence of only RNA in the final elution in the lanes indicated. Blots were probed with a DIG-labeled TERRA probe. (**C**) RNA enriched in R-Loops from Control-sh or METTL3-KD U-2 OS cells were isolated by the DRIP method followed by slot blot assay. Blot was probed with a DIG-labeled TERRA probe. Input samples probed with Alu probe served as a loading control. DRIP performed with IgG antibody or pre-treatment with RNase H served as a control. RNase A treatment was done to affirm the presence of only RNA in the final elution. Bar graph shows the quantification of the blots. Data are shown as mean ± SD from two biological replicates. Unpaired *t*-test was used, **** *p <* 0.0001. (**D**) TERRA RNA-FISH detecting TERRA foci (green) in Control-sh or METTL3-KD U-2 OS cells that were transfected with either Control siRNA or siRNAs against RNase H. Box plot shows the quantification of TERRA foci per nucleus. At least 100 cells were counted from three independent biological replicates. One-way ANOVA with Tukey’s *post hoc* test was used, **** *p <* 0.0001. R-loop PLA assay (**E**) PLA in U-2 OS TRF1-mCherry cells with overexpression of either RNase H^MUT^ (**left**) or RNase H^WT^ (**right**) depicting the interaction of TRF2 with R-Loop (S9.6 antibody). TRF1-mCherry (Red), PLA foci (green) in the nucleus (marked by DAPI). Box plot shows the number of overlaps of PLA signal and TRF1-mCherry per nucleus. At least 100 cells were counted from three independent biological replicates. Unpaired *t*-test was used, **** *p <* 0.0001. (**F-H**) PLA depicting the interaction of TRF2 with R-Loop (S9.6 antibody) in U-2 OS TRF1-mCherry cells with (**F**) Control-sh or METTL3-KD (**G**) Control siRNA or hnRNPA2B1 siRNA2. TRF1-mCherry (Red), PLA foci (green) in the nucleus (marked by DAPI). Box plot shows the number of overlaps of PLA signal and TRF1- mCherry per nucleus. At least 70 cells were counted from three independent biological replicates. Unpaired *t*-test was used, **** *p <* 0.0001. (**H**) U-2 OS TRF1-mCherry cells expressing dCasRx-FTO with either NTC (**left**) or TERRA guide RNA (**right**). Box plot shows the quantification of the overlapping PLA and TRF1- mCherry signal. At least 105 cells were counted from three independent biological replicates. Unpaired *t*-test was used, **** *p <* 0.0001. Scale bar is 10 μm.

To obtain specific enrichment of R-loop-associated TERRA RNA, we briefly treated the cell lysate with RNase A and the isolated genomic DNA with RNase III (58), to identify genuine R-loop-enriched RNAs (Figure 5A). Following the S9.6 IP, we performed extensive DNase I treatment to obtain R-loop-associated RNA (Figure 5A). Subsequently, we used the R-loop-associated RNA for slot blot assays and successfully detected enrichment of R-loop-associated TERRA (referred to as RL-TERRA) (Figure 5B). The enrichment of RL-TERRA decreased following RNase H treatment, and no enrichment was observed with the IgG control, confirming the specificity of the obtained RL-TERRA using our modified DRIP method. Furthermore, the enriched RL-TERRA was sensitive to RNase A treatment as well (Figure 5B). To investigate m^6^A modification present in R-loop structures, we used the R-loop-associated RNA obtained from our modified DRIP method as input for m^6^A RIP, which we refer to as DRImR (DRIP followed by m^6^A RIP) (Figure 5A, B). DRImR allowed us to detect abundant m^6^A modification in RL-TERRA (Figure 5B). Additionally, METTL3-KD led to compromised enrichment of RL-TERRA in U-2 OS cells, suggesting that TERRA m^6^A contributes to R-loop formation (Figure 5C).

To investigate the effect of R-loop removal on TERRA foci formation, we treated U-2 OS cells with RNase A alone or in combination with RNase H. Only RNase A had a moderate effect on TERRA foci in U-2 OS cells but combining RNase A and RNase H led to the disappearance of TERRA foci, suggesting the importance of R-loops in TERRA foci formation (Supplementary Figure S5A). RNase H1 (hereafter RNase H) has been implicated in maintaining telomeres in ALT+ cells by regulating TERRA R-loop formation. Depletion of RNase H in U-2 OS cells promotes TERRA R-loop accumulation and higher C-Circles (59). We reasoned that reduced TERRA foci in METTL3-KD cells were due to compromised R-loop and whether TERRA foci can be rescued by depleting RNase H. We were able to rescue TERRA foci in RNase H depleted METTL3-KD cells, suggesting the role of m^6^A in R-loop-driven TERRA foci formation (Figure 5D; Supplementary Figure S5B).

Furthermore, we developed a robust and fast proximity ligation assay (PLA)-based method to examine telomere R-loops using TRF2 and S9.6 antibodies (R-loop PLA). In our experiment, we used wild-type RNase H (RNase H^WT^) and a catalytically inactive mutant RNase H (RNase H^MUT^) as controls. We observed R-loop PLA signal in cells expressing RNase H^MUT^ but not RNase H^WT^ suggesting the specificity of the assay (Figure 5E). As expected, overexpression of RNase H^WT^ led to a decrease in overall R-loop content (Supplementary Figure S5C). RNase H^WT^ cells also exhibited fewer TERRA foci compared to RNase H^MUT^ cells (Supplementary S5D). Importantly, we did not detect any R-loop PLA signal in the PLA controls (Supplementary Figure S5E). Additionally, we tested the R-loop PLA assay in RNase H siRNA transfected cells and observed a significant increase in R-loop PLA signal in RNase H-depleted cells with a concomitant increase in DNA C-Circle content (Supplementary Figure S5F, G). This is consistent with our TERRA foci recovery phenotype of the RNase H-depleted METTL3-KD cells (Figure 5D).

We applied the R-loop PLA assay to Control-sh and METTL3-KD cells and observed a loss of PLA signal over the telomere in METTL3-KD cells, which is consistent with the data obtained using DRImR (Figure 5C, F). To verify the direct role of METTL3 in TERRA R-loop formation, we employed two approaches. Firstly, we overexpressed METTL3^WT^ or METTL3^MUT^ in U-2 OS cells and found that overexpression of METTL3^WT^ could enhance TERRA R-loop formation (Supplementary Figure S5H). In the second approach, we utilized the METTL3-dCasRx system to recruit shRNA-resistant METTL3 to TERRA in endogenous METTL3-depleted cells, where we previously observed rescue of the TERRA foci phenotype (Figure 3E). We found that recruitment of shRNA-resistant METTL3^WT^, but not METTL3^MUT^, could drive TERRA R-loop formation in METTL3-KD cells, as detected by R-loop PLA (Supplementary Figure S5I).

Moreover, the depletion of hnRNPA2B1 and the removal of TERRA m^6^A using dCasRx-FTO^WT^ also led to a loss of R-loop PLA signal (Figure 5G, H), suggesting that TERRA m^6^A and its reader hnRNPA2B1 are required for R-loop formation over telomeres.

### TERRA RNA is m^6^A modified in ALT+ NB cells and tumors

Our observations from U-2 OS cells suggest that METTL3 is required for telomere maintenance in ALT cells, and inhibiting METTL3 could be a therapeutic approach against ALT+ tumors. A significant portion of the primary NBs are ALT+ and patients with ALT+ tumors have a poor prognosis. Additionally, relapsed NBs are frequently found to be ALT+ (27,29,60,61). However, the role of TERRA in ALT+ NBs has not been extensively investigated. Understanding the mechanisms underlying telomere maintenance in ALT+ NB is crucial for developing novel treatment strategies for these tumors. In this study, we aimed to characterize the role of m^6^A modification of TERRA in telomere maintenance in ALT+ NB.

First, we checked TERRA expression in ALT+ and ALT-negative: ALT- [MYCN amplified (NMA) and non-amplified (Non-NMA)] NB cell lines. We observed higher TERRA expression in ALT+ (CHLA-90 and SK-N-FI) compared to ALT-NB cell lines (Figure 6A). ALT positivity of the NB cell lines was verified using the C-Circle assay (Supplementary Figure S6A). Furthermore, we performed m^6^A RIP using fragmented RNA from ALT+ and ALT-NB cell lines and observed robust enrichment of m^6^A-modified TERRA in ALT+ cells (Figure 6B). Consistently, prominent TERRA foci were observed in ALT+ but not ALT-NB cell lines (Figure 6C). The higher TERRA expression and m^6^A enrichment in ALT+ cells may contribute to TERRA foci formation. Additionally, the TERRA foci in ALT+ NB cells were sensitive to knockdown using locked nucleic acid (LNA) probes and frequently co-localized with m^6^A IF signals in the nucleus (Supplementary Figure S6B; Figure 6D). To further characterize TERRA expression and m^6^A enrichment in ALT+ (SK-N-FI) and ALT- [SK-N-BE(2)] NB cells we performed RNA-seq and m^6^A RIP-seq. Consistent with the slot blot assay, we observed lower TERRA expression in ALT-cells compared to ALT+ cells (Figure 6A; Supplementary Figure S6C). We also found enrichment of the "DRACH" motif in the m^6^A peaks obtained from both SK-N-FI and SK-N-BE(2) m^6^A RIP-seq data (Supplementary Figure S6D). Next, we checked for subtelomeric m^6^A enrichment in these two cell lines, mappability and spike-in normalized subtelomeric m^6^A enrichment (log_2_ m^6^A/input) was significantly higher in ALT+ SK-N-FI cells compared to ALT-SK-N-BE(2) cells (Figure 6E, F).

**Figure 6:**
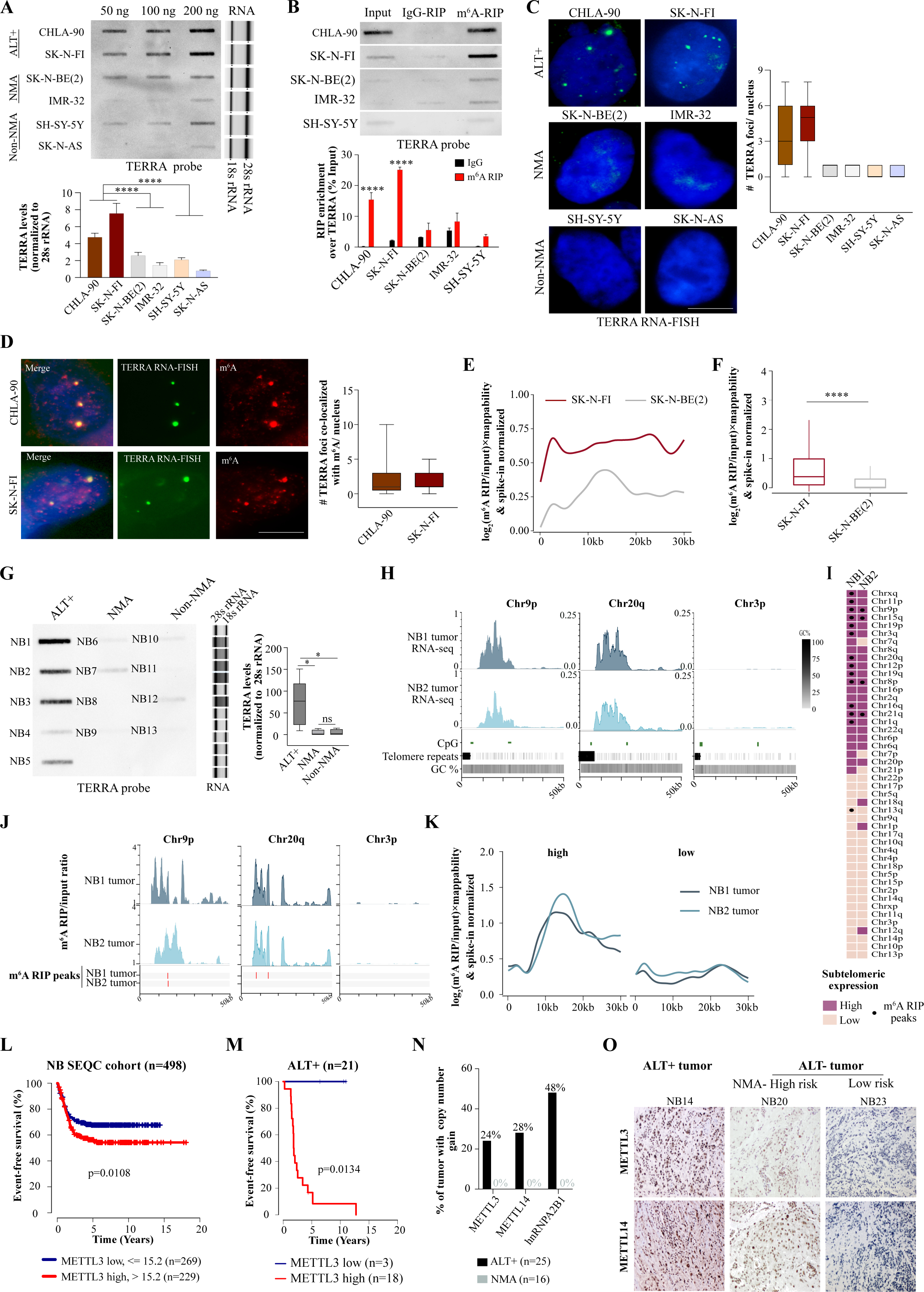
(**A**) Profiling TERRA expression in ALT+ and ALT-NB cell lines (with MYCN amplification [NMA] and without MYCN amplification [Non-NMA]) by slot blot assay performed with 50, 100, and 200 ng of RNA isolated from cell lines indicated. Blot probed with a DIG-labeled TERRA probe. TapeStation profile on the right shows 18s and 28s rRNA served as a loading control. Bar graph shows the TERRA level quantified from blots and normalized to 28s rRNA loading control. Data are shown as mean ± SD from three biological replicates. One-way ANOVA with Tukey’s *post hoc* test was used, **** *p <* 0.0001. (**B**) m^6^A RIP followed by TERRA slot blot on RNA isolated from ALT+ and ALT-cell lines. RIP with IgG severed as a negative control. Bar graph shows the quantification of the blots. Data are shown as mean ± SD from two biological replicates. Sidak’s multiple comparisons test was used, **** *p <* 0.0001. (**C**) TERRA foci (green) visualized by TERRA RNA-FISH in ALT+ (**upper panel**), NMA (**middle panel**), and Non-NMA (**lower panel**) NB cell lines. Box plot shows the quantification of the TERRA foci per nucleus. At least 50 cells were counted from two to three independent biological replicates. (**D**) TERRA foci (green) combined with m^6^A (red) IF in CHLA-90 (top) and SK-N-FI (bottom) cells. Box plot shows the number of overlaps of TERRA and m^6^A per nucleus. At least 50 cells were counted from two independent biological replicates. Scale bar is 10 μm. (**E**) The m^6^A enrichment profiles across all subtelomeres using T2T assembly in SK-N-FI and SK-N-BE(2) cells. The m^6^A RIP/input signals were normalized to spike-in and to the mappability likelihood of each chromosome ends. (**F**) Box plot of spike-in, mappability-normalized m^6^A RIP/input enrichment profiles of all subtelomeres in SK-N-FI and SK-N-BE(2). Statistical significance was calculated using the Wilcoxon test, **** *p <* 0.0001. (**G, left panel**) TERRA expression in ALT+, NMA, and Non-NMA NB patient tumor RNA samples were detected by slot blot assay performed with 100 ng of total RNA. (**Middle panel**) TapeStation profile shows 18s and 28s rRNA served as a loading control, samples are loaded as NB1 to NB13 from top to bottom. (**Right panel**) Box plot shows the TERRA level quantified from blots and normalized to 28s rRNA loading control. (**H**) Representative browser screenshot of 50 kb region around the telomere ends, showing TERRA RNA expression (CPM) from two of the active chromosome ends (Chr9p, Chr16q) and one inactive chromosome end (Chr3p) using Illumina short read RNA-seq performed on NB1 and NB2 tumor samples. CpG islands are marked with green bars and the telomeric repeats in this region are marked with black bars. (**I**) Heatmap summarizing RNA-seq and m^6^A RIP-seq data of two NB tumor samples (NB1 and NB2). Chromosomes are sorted based on high (dark-colored) and low (light-colored) subtelomeric TERRA transcription. Black dots denote m^6^A RIP-seq peaks identified using MACS peak caller in these NB tumor samples. (**J**) Genome browser screenshots showing m^6^A RIP/input ratio tracks for NB1 and NB2 tumor samples at two active chromosome ends (Chr12p, Chr20q) and one inactive chromosome end (Chr3p). Red bars mark m^6^A RIP peaks identified using MACS peak caller. (**K**) The m^6^A enrichment profiles across subtelomeres using T2T assembly in NB1 and NB2 tumors. The m^6^A RIP/input signals were normalized to spike-in and to the mappability likelihood of each chromosome ends. Subtelomeres (high and low TERRA transcription) were classified according to the normalized median expression shown. (**L**) Event-free survival of NB patients (n=498, SEQC cohort) with either low (blue) or high (red) expression of METTL3. (**M**) Event-free survival of ALT+ NB patients (n=21) with either low (blue) or high (red) expression of METTL3. (**N**) Bar graph showing the percentage of ALT+ or NMA tumors with copy number gain in *METTL3*, *METTL14*, or *hnRNPA2B1* genes. (**O**) Immunohistochemistry (IHC) analysis of METTL3 and METTL14 in human neuroblastoma tumors belonging to either of the subgroup (ALT+, NMA-high risk, or low risk). Sections were counterstained with hematoxylin. One representative image is presented from each subgroup of NB patients.

To gain a better understanding of TERRA expression in NB tumor samples, we compared TERRA expression in representative ALT+ and ALT-tumors (NMA and Non-NMA) for which we had access to RNA. Two of the NB tumors were previously reported by us as ALT+ (27) and for the remaining tumors investigated, we confirmed the ALT status using the C-Circle assay (Supplementary Figure S6E; Supplementary Table S2). We observed elevated levels of TERRA expression in ALT+ NBs, which is consistent with earlier observations (Figure 6G) (61). Next, we aimed to characterize TERRA expression and m^6^A positivity using RNA-seq and m^6^A RIP-seq in two ALT+ NBs (NB1 and NB2) that exhibited high TERRA expression in the slot blot assay (Figure 6G). The RNA-seq data from ALT+ NBs were used to define subtelomeres expressing high and low levels of TERRA, similar to the approach used in U-2 OS cells. We observed consistent transcriptional activity from the subtelomeres in both tumors, with some subtelomeres being silent or moderately transcribed (Figure 6H; Supplementary Figure S6F). Overall, the subtelomeres that actively transcribe TERRA were similar in both the NBs and U-2 OS cell lines, suggesting a commonality in TERRA transcription (Figure 6I, 2C). Additionally, we found that "DRACH"-like motifs were enriched in the m^6^A peaks obtained from both NBs (Supplementary Figure S6G). TERRA RNAs from several active subtelomeric regions exhibited enrichment of m^6^A modifications in NBs and frequently had m^6^A peaks as well (Figure 6I, J). The number of m^6^A peaks in NB1 was higher compared to NB2 in the subtelomeres, possibly due to tumor-specific variations (Figure 6I). Furthermore, the subtelomeric metagene profile of mappability and spike-in normalized m^6^A enrichment (log_2_ m^6^A/input) indicated that subtelomeres expressing elevated levels of TERRA were more robustly m^6^A-modified compared to subtelomeres with low expression in both ALT+ NBs (Figure 6K).

Moreover, we found that higher expressions of METTL3, METTL14, and hnRNPA2B1 in the NB cohort (n=498) were significantly associated with poor event-free survival (EFS) (Figure 6L; Supplementary Figure S6H). In a smaller cohort of ALT+ NBs (n=21), we observed that higher expression of METTL3 and METTL14 was significantly correlated with worse EFS, while hnRNPA2B1 showed a trend towards worse EFS, although it did not reach statistical significance (Figure 6M; Supplementary Figure S6I). Additionally, we observed frequent copy number gains of METTL3, METTL14, and hnRNPA2B1, in ALT+ tumors compared to NMA NBs (Figure 6N). Immunohistochemistry (IHC) on a panel of NBs revealed that ALT+ tumors expressed higher levels of METTL3 and METTL14 compared to low-risk NBs, while NMA-high-risk NBs exhibited moderate levels of METTL3 and METTL14 (Figure 6O; Supplementary Figure S6J; Supplementary Table S2).

Taken together, our data suggest the vital role of METTL3/METTL14 and other m^6^A-related genes in ALT+ NB and their association with poor clinical outcomes in NB patients.

### m^6^A modification in TERRA RNA is critical for telomere maintenance in ALT+ NB cells

To explore the role of METTL3-mediated m^6^A modification in ALT+ NB, we depleted METTL3 in ALT+ NB cells. We observed that siRNA-mediated transient depletion of METTL3 in ALT+ CHLA-90 and SK-N-FI cells led to a reduction in the number of TERRA foci, similar to U-2 OS cells (Supplementary Figure S7A). Next, we attempted to stably deplete METTL3 in SK-N-FI and CHLA-90 cells, but we were unsuccessful in generating stable METTL3-depleted cells after several attempts. These cells did not survive, suggesting that METTL3 is essential for the survival of ALT+ NB cells. Therefore, we generated doxycycline (DOX) inducible TetO METTL3-KD cells (Figure 7A). METTL3-depleted ALT+ cells showed a decrease in cell viability and reduction in colony formation compared to TetO Control cells (Supplementary Figure S7B; Figure 7B). Also, TERRA foci loss was evident in TetO METTL3-KD CHLA-90 and SK-N-FI cells (Figure 7C), but TERRA expression remained unchanged as visualized in the slot blot assay (Supplementary Figure S7C). METTL3 depletion resulted in increased accumulation of γ-H2AX over telomeres in both SK-N-FI and CHLA-90 cells, indicative of telomere damage (Figure 7D; Supplementary Figure S7D). METTL3 depletion led to a decrease in C-Circle formation in these ALT+ NB cells, suggesting compromised ALT activity (Supplementary Figure S7E). METTL3-depleted ALT+ NB cells also showed a decrease in RL-TERRA enrichment, similar to U-2 OS cells, suggesting compromised TERRA R-loop formation (Supplementary Figure S7F). We xenografted TetO Control and METTL3-KD SK-N-FI and CHLA-90 cells into immunocompromised mice and measured the effect on tumor formation following DOX induction. We observed that METTL3-KD led to reduced xenografted tumor growth in both ALT+ cell lines, with a more drastic effect in SK-N-FI cells (Figure 7E, F; Supplementary Figure S7G). METTL3-depleted SK-N-FI xenografts showed a decreased level of C-Circles and a reduction in telomere length (Figure 7G; Supplementary Figure S7H), suggesting a lack of telomere maintenance in the absence of METTL3. We observed an increase in the expression of senescence and apoptotic markers (*CDKN2A* and *CASP3*) with a concomitant decrease in the anti-apoptotic marker *BCL2* in METTL3-depleted SK-N-FI xenografts (Supplementary Figure S7I).

**Figure 7:**
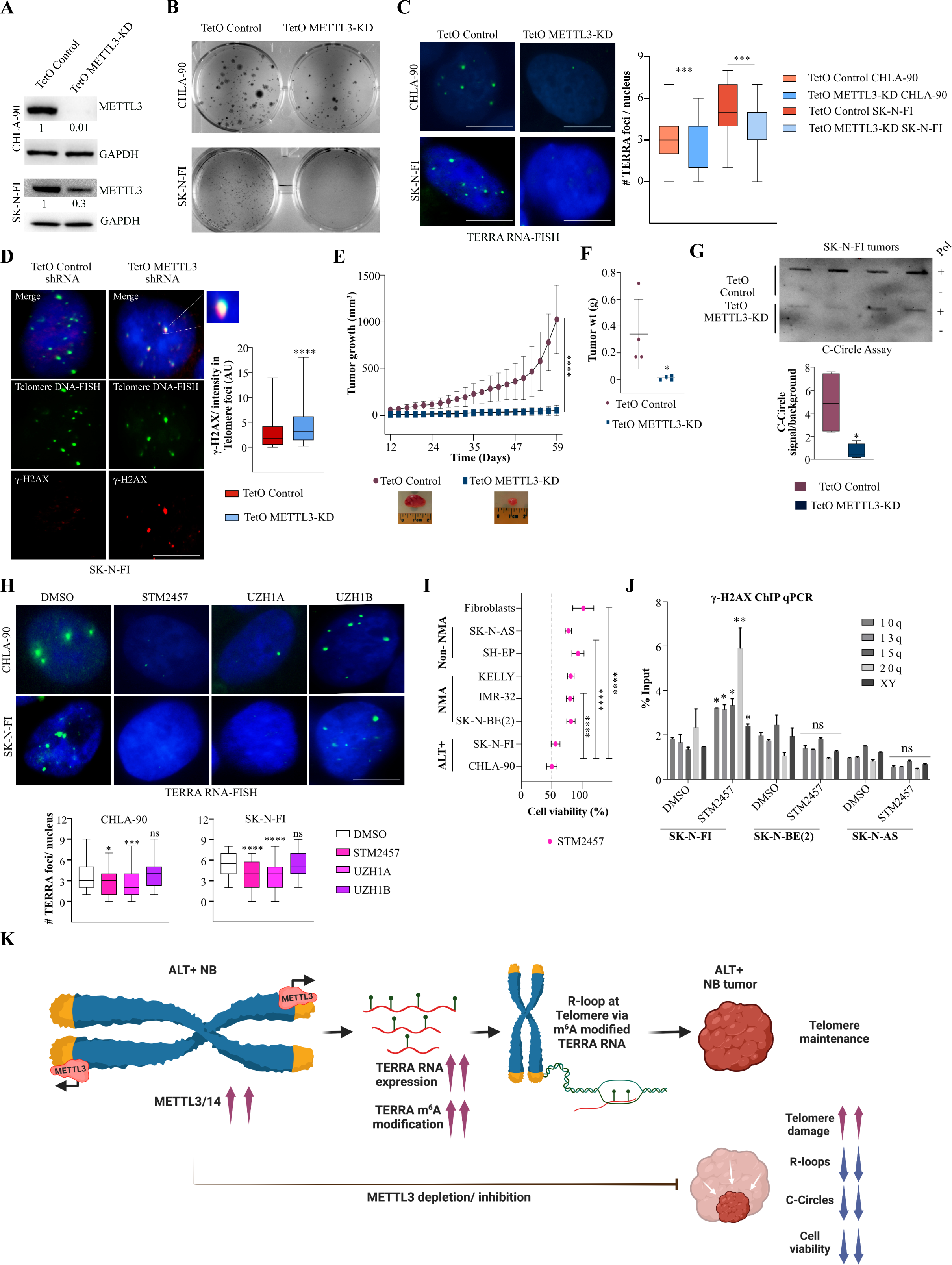
(**A**) Western blot showing METTL3 level in TetO Control and METTL3-KD ALT+ NB cells following DOX induction for 48 h. GAPDH was loading control. The values below indicate the fold change in levels of METTL3. (**B**) Representative images from colony formation assay performed on CHLA-90 (**top**) and SK-N-FI (**bottom**) cells with TetO Control or METTL3-KD. (**C**) TERRA RNA-FISH in TetO Control and METTL3-KD SK-N-FI and CHLA-90 cells. Box plot shows the quantification of the TERRA foci in the indicated conditions. At least 70 cells were counted from three independent biological replicates. Unpaired t-test was used, *** p < 0.001. (**D**) Accumulation of γ-H2AX (red) over telomere after 48 h of DOX induction in SK-N-FI TetO METTL3-KD cells. Telomeres were detected by telomere DNA-FISH (green). Box plot shows the γ-H2AX intensity in telomeric foci (Arbitrary units, AU). At least 50 cells were counted from two independent biological replicates. Unpaired *t*-test was used, **** *p <* 0.0001. (**E**) Line graph showing average tumor growth (mm^3^) of SK-N-FI cells (TetO Control and METTL3-KD) derived xenografts over time. Data are shown as mean ± SD. Sidak’s multiple comparisons test was used, **** *p <* 0.0001. (**F**) Scatter plot showing tumor weight post necropsies of xenografts derived from SK-N-FI cells with either TetO Control or METTL3- KD. Data are shown as mean ± SD. Unpaired *t*-test was used, * *p <* 0.05 (n=4). One representative image of the tumor is shown per condition. (**G**) C-Circle assay results visualized on slot blot. C-Circle assay with/without Phi29 polymerase (Pol+/Pol-) performed with DNA isolated from SK-N-FI cells derived xenografts with either TetO Control or METTL3-KD. Box plots show the quantification of the blots, data are presented as signal/background. Unpaired *t*-test was used, * *p <* 0.05 (n=4). (**H**) TERRA RNA-FISH detecting TERRA foci (green) in CHLA-90 (**top panel**) and SK-N-FI (**bottom panel**) cells treated with METTL3 inhibitors (10 μM STM2457, 50 μM UZH1A, and 50 μM UZH1B) for 6 h. DMSO treatment served as a control. Box plot showing quantification of the TERRA foci as indicated. At least 70 cells were counted from three independent biological replicates. Dunnett’s multiple comparisons test was used, * *p <* 0.05; *** *p <* 0.001; **** *p <* 0.0001. (**I**) Cell viability was measured by MTT assay after treating ALT+, ALT- (NMA and Non-NMA), and fibroblast cells with 10 μM of METTL3 inhibitor (STM2457) for 72 h. Data for each cell line were normalized to DMSO control. Data are shown as mean ± SD from three biological replicates. One-way ANOVA with Tukey’s *post hoc* test was used, **** *p <* 0.0001. (**J**) Bar plot shows the percentage input values of γ-H2AX enrichment over selected telomere ends from ChIP qPCR data in DMSO and METTL3 inhibitor STM2457 treated SK-N-FI, SK-N-BE(2), and SK-N-AS cells. STM247 treatment was done for 72 h. Data are shown as mean ± SD from two independent biological replicates. Unpaired *t*-test was used, \*\**p <* 0.01, \**p <* 0.05 ns - nonsignificant *p* > 0.05. (**K**) The model illustrates the role of METTL3-mediated TERRA m^6^A modification in telomere maintenance of ALT+ NB. Elevated levels of METTL3/14 and TERRA m^6^A promote telomere maintenance in ALT+ NBs through R-loop formation. In contrast, METTL3 depletion or inhibition in NBs results in reduced TERRA m^6^A levels, leading to compromised R-loop formation, decreased DNA C-Circle content, and an accumulation of telomere damage. The deficiency in telomere maintenance after METTL3 depletion leads to tumor regression.

To test if TERRA foci formation in ALT+ NB cells is hnRNPA2B1-dependent, we depleted hnRNPA2B1, which led to the loss of TERRA foci (Supplementary Figure S7J). We also observed that TERRA foci co-localized with hnRNPA2B1 in ALT+ NB cells (Supplementary Figure S7K). Finally, we investigated the therapeutic potential of inhibiting the METTL3 enzyme in ALT+ NB cell lines. Treatment of ALT+ NB cells with METTL3 inhibitors STM2457 and UZH1A for 6 h led to a dramatic decrease in TERRA foci formation compared to cells treated with UZH1B (a less potent inhibitor) and DMSO control (Figure 7H). Next, we treated a panel of NB cell lines (ALT+, ALT-[NMA and Non-NMA]) and normal fibroblast cells with the METTL3 inhibitor STM2457 for 72 h. We observed that ALT+ NB cell lines were more sensitive to METTL3 inhibition compared to ALT- (NMA, Non-NMA) NB cell lines, while fibroblast cells were unaffected (Figure 7I). Treatment with STM2457 resulted in the enhanced accumulation of γ-H2AX over telomere specifically in ALT+ but not ALT-NB cell lines, suggesting that METTL3 inhibition induced telomere damage was more specific to ALT+ NB cells (Figure 7J; Supplementary Figure S7L).

In conclusion, our data demonstrate that TERRA m^6^A and METTL3 play a vital role in telomere maintenance in ALT+ NB. Inhibiting METTL3 could be a potential therapeutic target for ALT+ NB (Figure 7K).

## Discussion

Our study has identified the role of m^6^A RNA modification in TERRA, a telomere-derived noncoding RNA that plays a crucial role in telomere maintenance in ALT+ cells. Our data suggest that the METTL3/14 complex is recruited to telomeres in ALT+ U-2 OS cells. It is intriguing to speculate that TERRA G-quadruplex, derived from telomeric repeats, may guide the METTL3/14 complex to the telomeres, as recent studies have shown the binding of METTL14 to TERRA G-quadruplexes (25). Depletion of METTL3 leads to a decrease in m^6^A modification in the TERRA transcript and loss of TERRA foci in ALT+ cells. By recruiting shRNA-resistant dCasRx-METTL3 to TERRA in endogenous METTL3-depleted cells, we were able to rescue the loss of TERRA phenotype, suggesting a direct role of METTL3-mediated m^6^A modification in TERRA function. Modulating the m^6^A level through METTL3-KD results in the compromised repair of telomeric DNA, as visualized by increased accumulation of γ-H2AX over the telomeres. Our data indicate that m^6^A-containing TERRA foci are required for repairing bleomycin-induced telomere damage. Additionally, depletion of METTL3 in U-2 OS cells leads to reduced telomere content, with frequent detection of telomere-free ends in metaphase spreads. Consistent with our observations, a recent study also reported the role of m^6^A modification in TERRA in telomere maintenance in ALT+ cells, and METTL3-KD was found to induce telomere shortening in ALT+ cells (46). The authors also reported the presence of m^6^A modification in the subtelomeric sequences of TERRA. In line with this, our m^6^A RIP-seq and xPore-identified m^6^A sites from long-read RNA-seq data also demonstrate enrichment of m^6^A in TERRA transcripts derived from subtelomeric regions. Our study, using m^6^A RIP with fragmented RNA (Figure 1B, C), *in vitro* m^6^A methyltransferase assay of telomeric repeats (Supplementary Figure S1B), and TERRA RNA-MS analysis (Figure 1D-G), suggests the presence of m^6^A modification over UUAGGG telomeric repeats in TERRA. Although the efficiency of m^6^A deposition by METTL3/METTL14 on telomeric repeats was comparatively low compared to the DRACH sequence *in vitro*. However, the presence of multiple copies of telomeric repeats within a single TERRA molecule could drive effective m^6^A modification *in vivo*. This m^6^A modification of TERRA in the repeats could be further enhanced by the presence of G-quadruplex-like structures in TERRA. Modulating m^6^A specifically in the telomeric repeats using dCasRx-FTO recapitulated the METTL3-KD phenotype, further suggesting the importance of m^6^A modification at telomeric repeat sequences in TERRA. TERRA subtelomeric sequences containing the "DRACH" motif showed differential enrichment in m^6^A RIP-seq data obtained from Control-sh and METTL3-KD cells. The m^6^A sites detected by xPore also included the "DRACH" motif. Since our long-read RNA-seq data was generated using poly-A selected RNA, it may not include m^6^A sites present in the non-poly-A fraction of TERRA (62). Our m^6^A RIP-seq and xPore data are consistent with the notion, as reported by Chen et al (46), that "DRACH"-like sequences in TERRA could also be targeted by METTL3 for m^6^A modification. Taken together, our findings provide evidence that both telomeric repeats and subtelomeric sequences contribute to the m^6^A enrichment of TERRA. Given the complexity of TERRA RNAs, which are derived from multiple chromosome ends and the presence of telomeric repeats not only at the ends of the chromosomes but also interspersed in the subtelomeric region (Figure 2B), it is challenging to precisely determine the exact location of m^6^A modification in TERRA.

Our mechanistic study revealed that m^6^A modification in TERRA promotes the association of the m^6^A reader protein hnRNPA2B1. Another m^6^A reader protein, YTHDC1, has also been shown to interact with TERRA in an m^6^A-dependent manner to regulate TERRA stability (46). Although we observed a moderate decrease in TERRA in METTL3-KD U-2 OS cells, we did not see any change in TERRA levels in SK-N-FI cells with inducible METTL3-KD. In U-2 OS cells, hnRNPA2B1-KD, and the specific removal of TERRA m^6^A using an inducible dCasRx-FTO^WT^ did not impact TERRA levels. Short-term treatment (6 h) with the METTL3 inhibitor resulted in the loss of TERRA foci, but TERRA levels remained unchanged. This suggests that m^6^A modification is crucial for the telomere targeting of TERRA. We observed that in the absence of telomere localization in METTL3 and hnRNPA2B1 depleted cells, a fraction of TERRA showed cytoplasmic localization. We believe that the major event in the absence of TERRA m^6^A or m^6^A reader proteins, such as hnRNPA2B1, is the delocalization of TERRA from the telomere. Changes in the nuclear/cytoplasmic localization and stability of TERRA (46) are subsequent events that may not always affect the steady-state level of TERRA. Other factors such as the THO complex and SMG proteins, known to regulate telomere association of TERRA, also do not significantly affect the steady-state level of TERRA either (21,63).

The m^6^A reader protein hnRNPA2B1 does not possess the YTH domain for m^6^A recognition but has been proposed to identify RNA through an "m^6^A switch" mechanism (64). Interestingly, the motif recognized by hnRNPA2B1 is UAGG-like sequences (65), whereas the major motif for m^6^A is "DRACH"-like sequences (66). UUAGGG-containing TERRA repeats have been shown to bind hnRNPA2B1 *in vitro*, and hnRNPA2B1 also recognizes UAGGG-containing motifs *in vivo* (67,68). Our *in vitro* and *in vivo* observations suggest that m^6^A modification in TERRA is critical for hnRNPA2B1 association. We identified the presence of m^6^A in the non-canonical UUAGGG context in TERRA. Similar observations of m^6^A in non-canonical contexts have been made in other lncRNAs such as HSATIII (AAUG repeats) and Xist (AUCG context in the tetra loop of Repeat-A) lncRNAs (16,17). In the case of Xist, m^6^A sites in the AUCG context in Repeat-A create local unfolding that favors recognition by YTHDC1 (16). The "m^6^A switch" mechanism also suggests that the presence of m^6^A increases RNA accessibility, thereby enhancing the association of RNA-binding proteins like hnRNPC and RBMX with RNA molecules (9,69).

We observed that both m^6^A modification and hnRNPA2B1 interaction are required for TERRA targeting to telomeres through R-loop formation. R-loop-associated TERRA (RL-TERRA) was found to be abundantly m^6^A modified, as detected by DRImR, which allows the combined detection of the RNA strand of the R-loop and m^6^A modification. We speculate that m^6^A modification in telomeric repeats containing UUAGGG might promote the local unfolding of TERRA during transcription, leading to the efficient recruitment of hnRNPA2B1 and other repair proteins, such as RAD51, to facilitate R-loop formation. R-loop formation is essential for the telomere targeting of TERRA after transcription (20). Chen et al. reported compromised RAD51 recruitment to telomeres in METTL3-KD cells, and defective RAD51 recruitment might affect TERRA-dependent R-loop formation (20,46). The mechanism of chromatin targeting by lncRNAs is still not well understood. Increasing evidence suggests that m^6^A modification in lncRNAs might promote chromatin association, and m^6^A-modified nascent RNAs accumulate at the site of transcription (36). Our observations on TERRA provide evidence that m^6^A-mediated R-loop formation might be a general mechanism utilized by other lncRNAs for chromatin targeting (36).

METTL3 has been shown to modify m^6^A DNA damage-associated RNA (49). METTL3- mediated m^6^A modification of chromatin-associated RNA at double-strand breaks (DSBs) facilitates the recruitment of RAD51 and BRCA1 to promote efficient homologous recombination (HR)-mediated repair of DSBs in an R-loop-dependent manner (50). Similarly, our data also suggest that METTL3-dependent m^6^A modification of TERRA can promote R-loop formation and thereby contribute to telomere maintenance in ALT+ cells through HR (59). Our study suggests that the m^6^A-dependent localization of TERRA to telomere, enabling HR-mediated telomere maintenance in ALT+ cells, represents a specialized and locus-specific mechanism. This mechanism can be seen as a refined adaptation of the broader m^6^A/RNA-dependent DNA repair pathway (49). The compromised telomere targeting of TERRA in the absence of m^6^A might affect the recruitment of DNA repair proteins like BRCA2, which are involved in recombination-dependent telomere maintenance in ALT+ cells (20). m^6^A-dependent TERRA R-loop formation and telomeric recombination in ALT+ cells are tightly regulated by RNase H and the THO complex, which can restrain R-loop formation over telomeres (59,63). TERRA has also been shown to frequently interact with epigenetic modifiers that are required for telomere maintenance (57,70). In METTL3-KD cells, the lack of telomere targeting in TERRA might result in altered chromatin structure at ALT+ telomeres, thereby resulting in a defect in telomere maintenance as well. However, the extent of chromatin changes over telomeres in METTL3- depleted/inhibited ALT+ cells will require further studies.

A significant portion of different cancer types, including NB and sarcoma, exhibit ALT as a telomere maintenance mechanism. ALT+ tumors often do not respond well to available treatment strategies (30). Leveraging our detailed mechanistic insights regarding TERRA m^6^A modification in telomere maintenance of ALT+ cells, we further investigated the role of TERRA m^6^A in NB. We provide evidence that TERRA in ALT+ NB cells exhibits high levels of m^6^A modification compared to ALT-NB cells. This is the first report of TERRA m^6^A modification in RNA derived from ALT+ NB tumors. TERRA is upregulated in ALT+ NB tumors, and it is worth exploring whether the loss of the chromatin remodeler protein ATRX (61) plays a role in driving TERRA expression and m^6^A modifications in ALT+ NB tumors. Our analysis indicates that higher expression levels of METTL3/14 and hnRNPA2B1 predict poor EFS in NB patients, and high expression of METTL3/14 in an ALT+ background correlates with the worst disease prognosis. *In vitro* and mouse xenograft experiments showed that METTL3-KD in ALT+ NB cells resulted in reduced proliferation. Moreover, ALT+ NB cells exhibited greater vulnerability to METTL3 inhibition compared to ALT-NB cells. Inhibition of METTL3 led to the accumulation of telomere damage specifically in ALT+ NB cells. Further investigation is needed to understand how ALT-cells are sensitive to METTL3 inhibition. As METTL3 is required to repair DNA damage in general, one possibility could be the ALT-NB cells will have problems repairing DNA damage (in non-telomeric regions) when METTL3 is inhibited.

Collectively, our data suggest that inhibiting METTL3 could be a novel therapeutic approach for the unfavorable high-risk subset of ALT+ NBs. Furthermore, METTL3 inhibition may hold potential as a therapeutic strategy for other ALT+ cancers.

## Acknowledgments

We would like to thank the core facility at Novum, BEA, Bioinformatics and Expression Analysis, which is supported by the board of research at the Karolinska Institute and the research committee at the Karolinska hospital for help with sequencing. Proteomics core facility at GU for their support with LC-MS/MS. The computations and data handling were enabled by resources in project SNIC-2022-22-85 provided by the Swedish National Infrastructure for Computing (SNIC) at UPPMAX, partially funded by the Swedish Council through grant agreement no. 2018-05973. We acknowledge Anders Sjölander for assistance concerning technical and implementational aspects of the UPPMAX resources.

## Funding

This work was funded by grants from the Swedish Research Council (Vetenskapsrådet, 2018- 02224) and project grants from Svenska Läkaresällskapet, Barncancerfonden, Cancerfonden, Åke Wibergs Stiftelse, and Kungl Vetenskaps-och Vitterhets-Samhället (KVVS) grant to TM, 4 years research position grant from Barncancerfonden to TM, Cancerfonden, Tore Nilsons Stiftelse and Bollan scholarship to RV, Assar Gabrielsson Fond to RV and KT.

## Notes

### Competing Interest Statement

The authors have declared no competing interest.

### Summary of Updates

Revision across all figures. We have identified the direct role of METTL3 in TERRA function.

