## Supplementary data for "METTL3 drives telomere targeting of TERRA lncRNA through m^6^A-dependent R-loop formation: a therapeutic target for ALT-positive neuroblastoma"

Vaid, Thombare, et al

Department of Laboratory Medicine, Institute of Biomedicine, University of Gothenburg, Gothenburg, Sweden.

#### Contents

Supplementary Figures S1-S7

Supplementary methods

Reference

#### Other Supplementary Materials for this manuscript

Supplementary Table S1: Sequence of siRNAs, shRNAs, qPCR primers, probes, and other oligos used in the study. Also contains information regarding all the antibodies used in the study.

Supplementary Table S2: Information about neuroblastoma tumor samples.

### Supplementary Figure S1

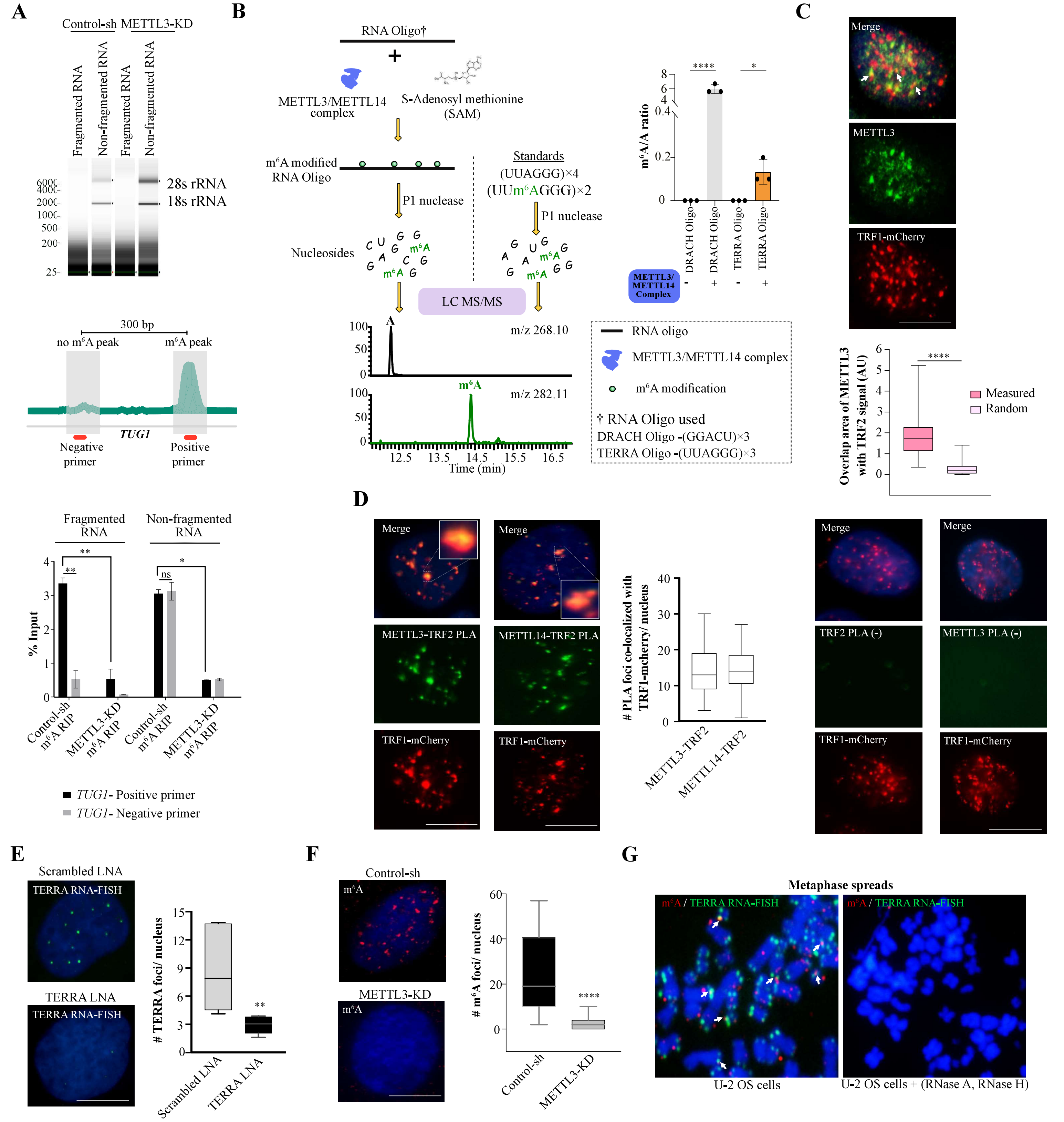

**Supplementary Figure S1:**

(**A, top panel**) TapeStation profile of RNA isolated from Control-sh or METTL3-KD U-2 OS cells which is either fragmented or non-fragmented. (**Middle panel**) Schematic diagram showing the location of the primers over $m^6A$ positive (positive primer) and $m^6A$ negative (negative primer) region over *TUG1* gene. (**Lower panel**) $m^6A$ RIP-qPCR with *TUG1*- positive and negative primers. Data are represented as a percentage of input and shown as mean ± SD from two biological replicates. Two-way ANOVA with Sidak's *post hoc* test was used, ** $p < 0.01$, * $p < 0.05$. These data served as a control for Figure 1C.

(**B**) Schematic diagram shows *in vitro* METTL3/METTL14 methyltransferase assay performed with DRACH oligo or TERRA repeat containing oligo. “A” and “$m^6A$” were detected and quantified by LC-MS/MS-based method. RNA oligo standard with or without $m^6A$ was used as a control. Bar graph shows the ratio of $m^6A/A$ in the indicated conditions. Unpaired *t*-tests were used, * $p < 0.05$; **** $p < 0.0001$.

(**C**) METTL3 (green) IF was performed in U-2 OS cells expressing TRF1-mCherry (red). White arrows indicate the co-localization of METTL3 and TRF1-mCherry. Box plot shows the overlap area of METTL3 with TRF1 (Arbitrary units, AU). The area of the overlapping METTL3 and TRF1-mCherry signal was quantified using the Interaction Factor package in ImageJ. To assess the significance of the observed overlapped area, the METTL3 signal was randomized for each image. The means of 50 randomizations were then plotted alongside the experimentally observed values. This analysis allowed us to evaluate the statistical significance of the observed overlap between METTL3 and TRF1-mCherry signals. At least 70 cells were counted from two independent biological replicates. Unpaired *t*-test was used, **** $p < 0.0001$. Scale bar is 10 μm.

(**D**) Proximity ligation assay (PLA) in U-2 OS cells expressing TRF1-mCherry depicting the interaction of TRF2 with METTL3 and METTL14. TRF1-mCherry (red), PLA foci (green) in the nucleus (marked by DAPI in blue). Box plot shows the number of co-localizations between TRF1-mCherry and PLA signal. Background control for PLA with only TRF2 and only METTL3 antibody in U-2 OS cells expressing TRF1-mCherry in the nucleus (marked by DAPI). At least 100 cells were counted from three independent biological replicates.

(**E**) TERRA foci (green) in U-2 OS cells treated with either Scrambled or TERRA LNA. Box plot shows the quantification of number of TERRA foci per nucleus. At least 70 cells were counted from three independent biological replicates. Unpaired *t*-test was used, ** $p < 0.01$. Scale bar is 10 μm.

(**F**) m$^6$A (red) IF was performed in Control-sh or METTL3-KD U-2 OS cells. Box plot shows the number of m$^6$A foci per nucleus. At least 80 cells were counted from three independent biological replicates. Unpaired *t*-test was used, **** $p < 0.0001$.

(**G**) m$^6$A (red) IF along with TERRA (green) RNA-FISH was performed on metaphase spreads of U-2 OS cells. White arrows indicate the co-localization of m$^6$A and TERRA.

#### Supplementary Figure S2

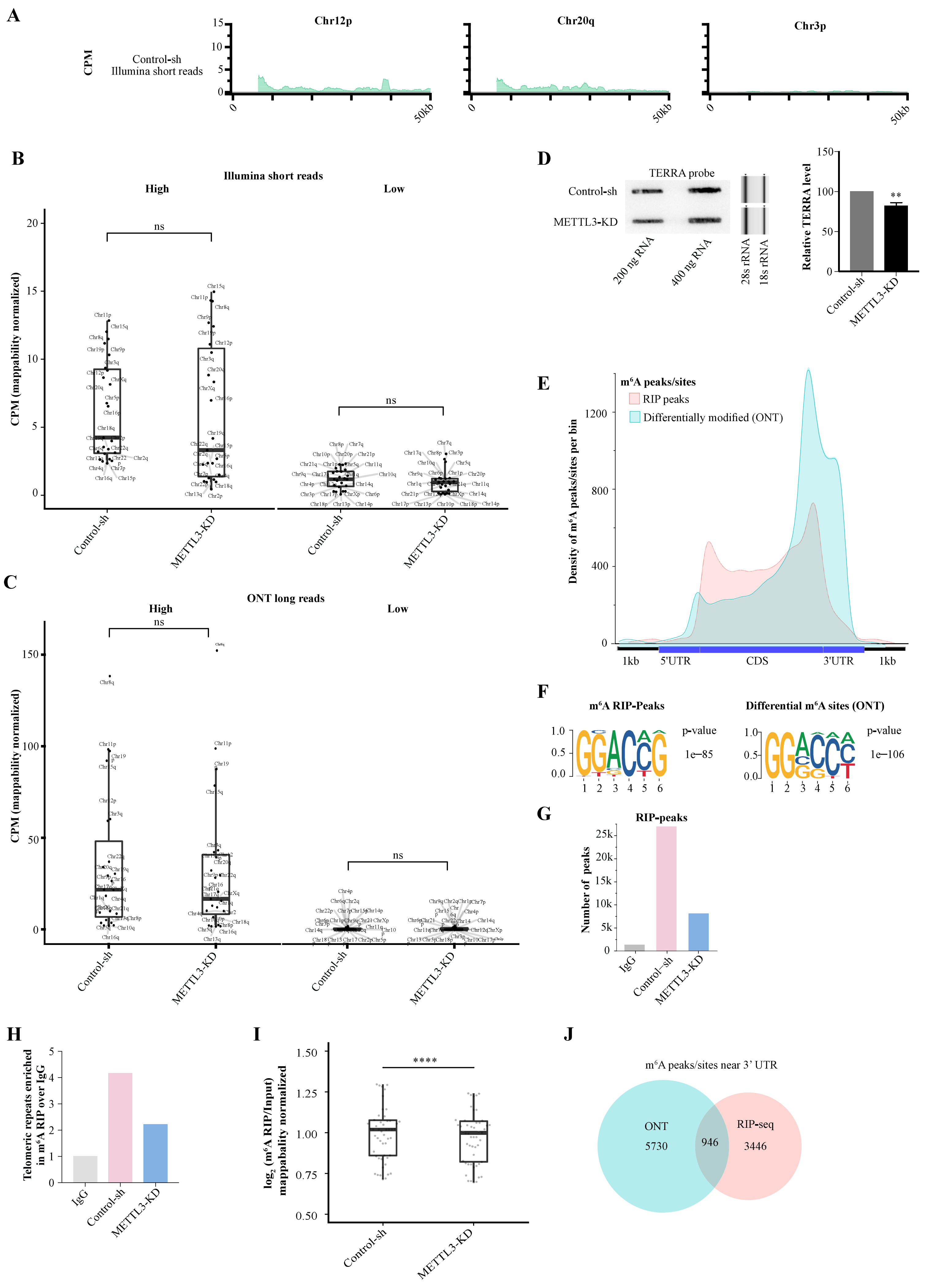

### Supplementary Figure S2 continue

**K**

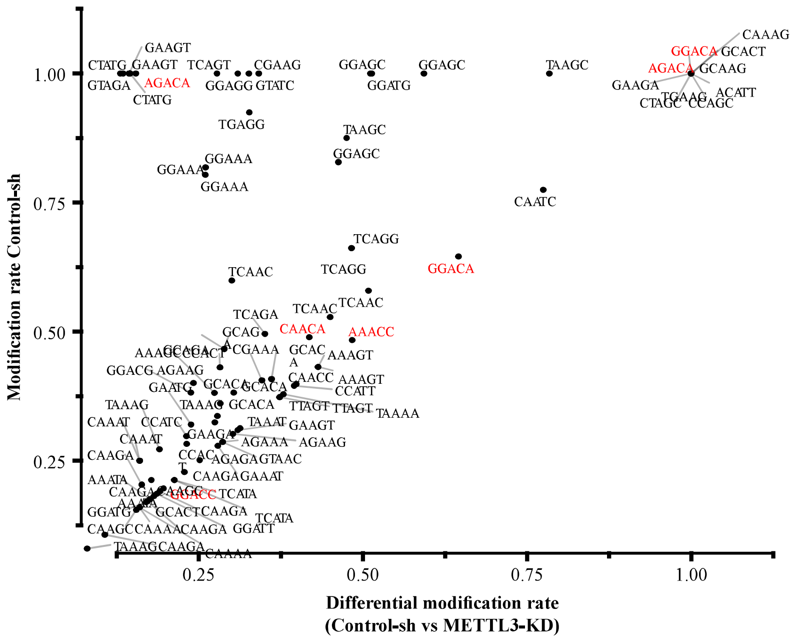

**Supplementary Figure S2:**

(**A**) Genome browser tracks of normalized CPM coverage obtained from Strand-specific Illumina short-reads in U-2 OS cells showing transcription away from telomere ends (direction of transcription is opposite to TERRA). Reads were mapped to the positive strand on p arms, and the negative strand on q arms, to match the direction of transcription, are shown for two active chromosome ends (Chr12p, Chr20q) and one inactive chromosome end (Chr3p).

(**B-C**) Characterizing the high and low TERRA expressing subtelomeres using normalized read counts from either (**B**) Illumina short reads or (**C**) ONT-long reads. Read counts were normalized using CPM and to the mappability likelihood of each chromosome ends. Statistical significance was calculated using the two-sided paired *t*-test.

(**D**) Slot blot with total RNA isolated from Control-sh or METTL3-KD U-2 OS cells. Blot probed with a DIG-labeled TERRA probe. TapeStation profile showing 18s and 28s rRNA served as a loading control. Bar graph shows the quantification of the blot normalized to the respective loading control. Unpaired *t*-test was used, ** $p < 0.01$.

(**E**) Metagene plot showing density distribution of $m^6A$ peak or $m^6A$ sites per bin from both $m^6A$ RIP peaks (red) and $m^6A$ xPore sites (blue). (**F**) Identified top enriched motif logo from *de novo* motif analysis of $m^6A$ peaks from $m^6A$ RIP-seq and $m^6A$ xPore sites. (**G**) Bar plot summarizing peaks identified genome-wide in RIP-seq experiment. (**H**) Telomeric repeat reads enriched in $m^6A$/IgG RIP-seq using TelomereHunter.

(**I**) Enrichment of $m^6A$ RIP/input signal at DRACH motif coordinates located within the subtelomeric $m^6A$ RIP-seq peaks in Control-sh and METTL3-KD U-2 OS cells. Statistical significance was calculated using the two-sided paired *t*-test. **** $p < 0.0001$.

(**J**) Venn diagram showing the overlap between $m^6A$ RIP peaks (red) and $m^6A$ xPore sites (blue) at the 3' UTR.

(**K**) Scatter plot depicting differentially modified sites identified using xPore from Control-sh and METTL3-KD U-2 OS long-reads. Predicted modified k-mers within the 30 kb subtelomeres regions showing a positive modification rate in Control-sh within the NNANN context being differentially modified as compared to METTL3-KD are shown. DRACH motifs identified are marked in red.

#### Supplementary Figure S3

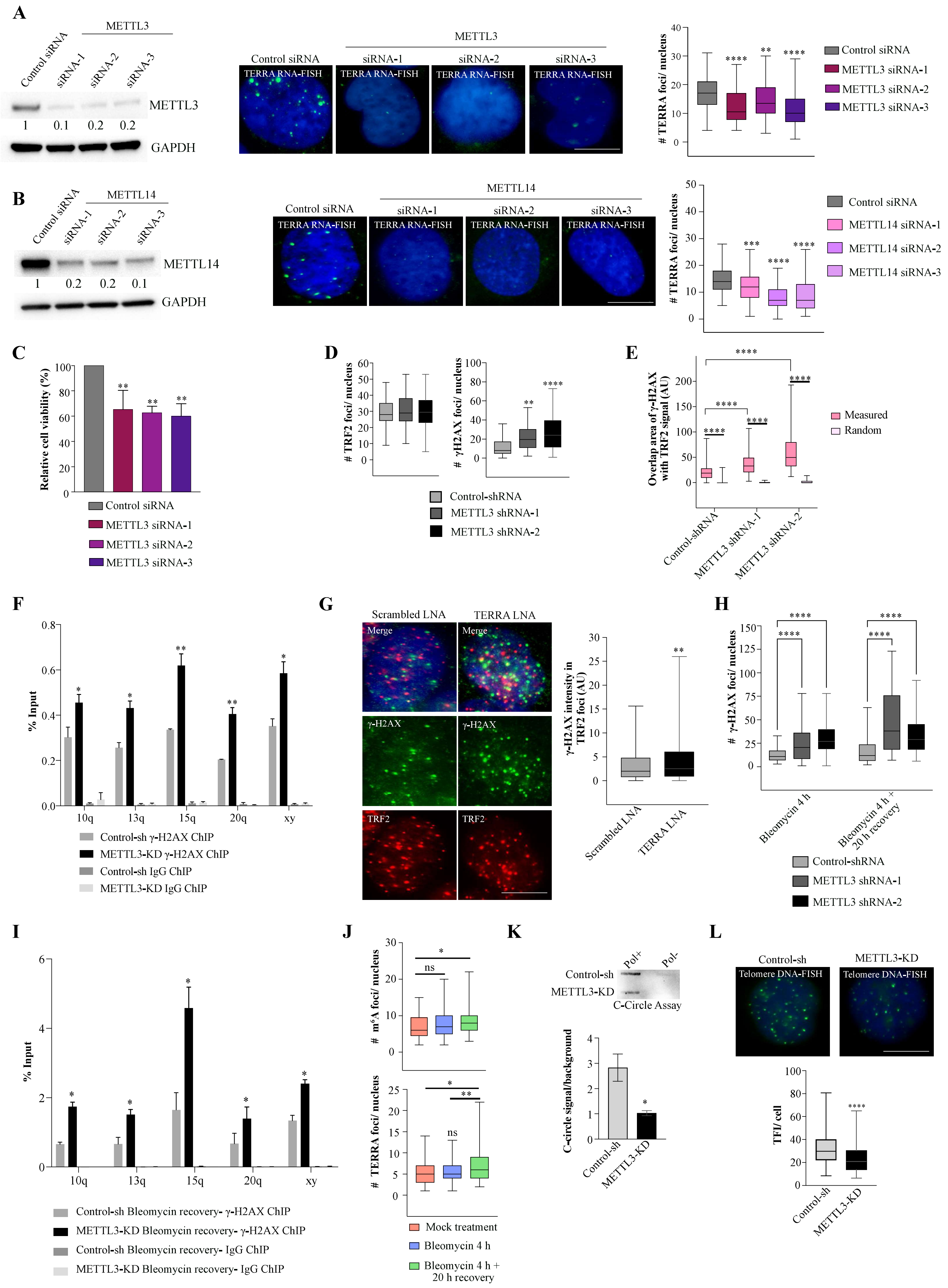

#### Supplementary Figure S3 continue

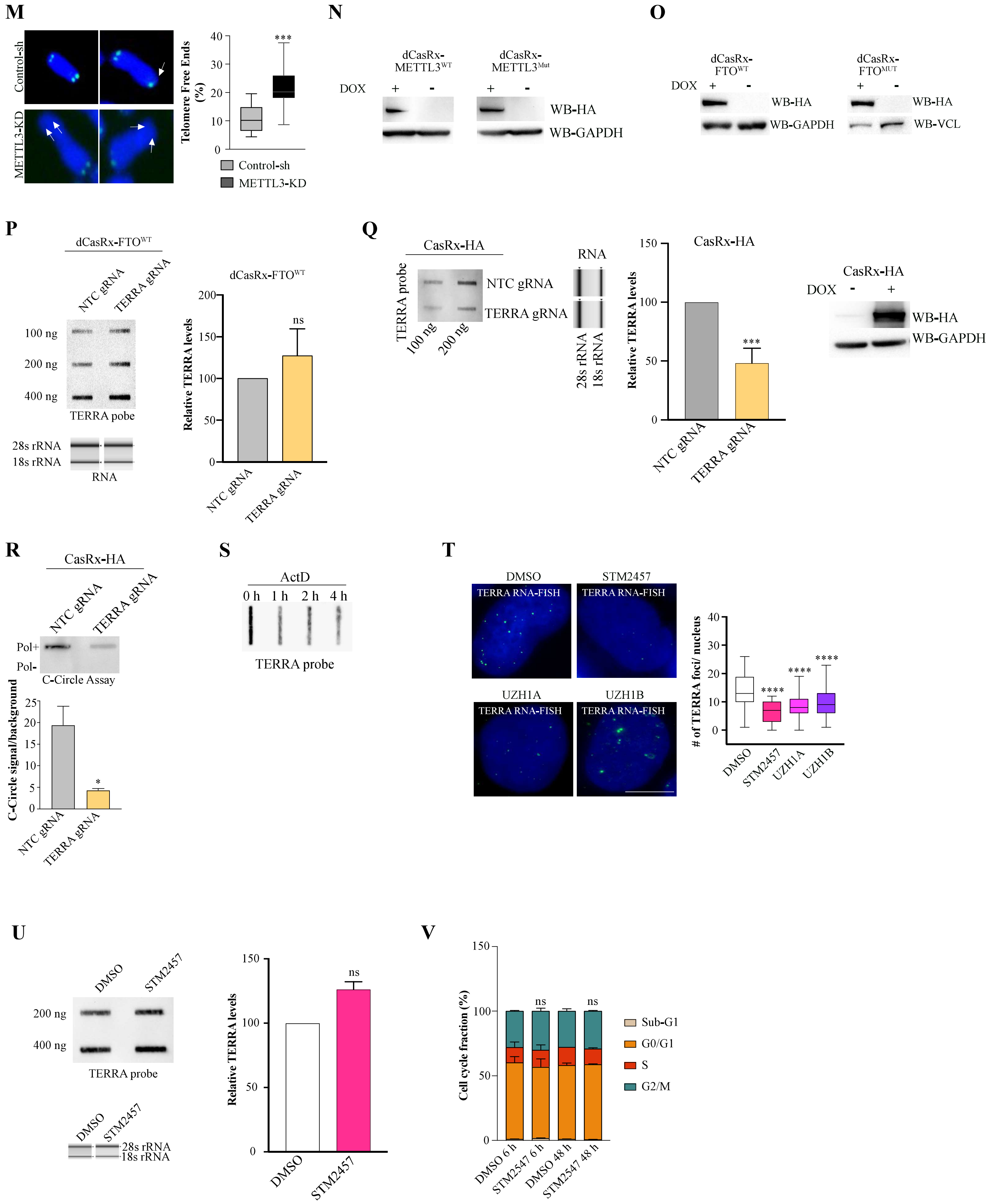

**Supplementary Figure S3:**

(**A, left panel**) Western blot to verify METTL3 siRNA-mediated KD in U-2 OS cells. Vinculin was used as a loading control. (**Middle panel**) TERRA foci (green) in U-2 OS cells after siRNA-mediated KD of METTL3. (**Right panel**) Box plot showing the number of TERRA foci per nucleus in the conditions indicated conditions. At least 100 cells were counted from three independent biological replicates. Dunnett's multiple comparisons test was used, ** $p < 0.01$; **** $p < 0.0001$.

(**B**, **left panel**) Western blot to verify METTL14 siRNA-mediated KD in U-2 OS cells. Vinculin was used as a loading control. (**Middle panel**) TERRA foci (green) in U-2 OS cells within Control or METTL14 siRNA treated cells. Nucleus marked by DAPI. (**Right panel**) Box plot shows TERRA signal per nucleus in the indicated conditions. At least 100 cells were counted from three independent biological replicates. Dunnett's multiple comparisons test was used, *** $p < 0.001$; **** $p < 0.0001$.

(**C**) Bar graph displaying cell viability after siRNA-mediated METTL3 KD U-2 OS cells relative to Control siRNA treatment. Data are shown as mean ± SD from two independent biological replicates. Dunnett's multiple comparisons test was used, ** $p < 0.01$.

(**D**) Box plots showig TRF2 and γ-H2AX foci number per nucleus in Control-sh and METTL3-KD U-2 OS cells. Data from images presented in Figure 3B. One-way ANOVA with Tukey's *post hoc* test was used, ** $p < 0.001$ **** $p < 0.0001$.

(**E**) For IF performed in Figure 3B, the overlap area of γ-H2AX with TRF2 in Control-sh and METTL3-KD U-2 OS cells is presented in box plots (Arbitrary units, AU). The area of the overlapping γ-H2AX and TRF2 signal was quantified using the Interaction Factor package in ImageJ and the statistical significance between observed versus random overlap was measured as described in Supplementary Figure S1C. Two-way ANOVA with Sidak's *post hoc* test was used, **** $p < 0.0001$.

(**F**) Bar plot shows the percentage input values of γ-H2AX enrichment over selected telomere ends from ChIP qPCR data in Control-sh and METTL3-KD U-2 OS cells. IgG ChIP served as a negative control. Data are shown as mean ± SD from two independent biological replicates. Unpaired *t*-test was used, *$p < 0.05$, **$p < 0.01$.

(**G**) Localization of the γ-H2AX (green) over telomere in Scrambled or TERRA LNA cells. Telomere is detected by TRF2 (red). Box plot shows the γ-H2AX intensity in TRF2 foci (Arbitrary units, AU). At least 80 cells were counted from three independent biological replicates. Unpaired *t*-test was used, **** $p < 0.0001$.

(**H**) γ-H2AX foci number per nucleus in bleomycin-treated Control-sh and METTL3-KD U-2 OS cells 4 h post-treatment and 4 h bleomycin treatment followed by 20 h of recovery without bleomycin. Data from images presented in Figure 3C. Two-way ANOVA with Sidak's *post hoc* test was used, **** $p < 0.0001$.

(**I**) γ-H2AX ChIP qPCR data, represented as percentage input over selected telomere ends in Control-sh and METTL3-KD U-2 OS cells treated with bleomycin for 4 h followed by 20 h of recovery without bleomycin. IgG ChIP served as a negative control. Data are shown as mean ± SD from two independent biological replicates. Unpaired *t*-test was used, *$p < 0.05$.

(**J**) Number of $m^6A$ (**upper panel**) and TERRA foci (**lower panel**) per nucleus in Mock treatment, 4 h bleomycin treatment, and 4 h bleomycin treatment followed by 20 h recovery without bleomycin. Data from images are presented in Figure 3D. One-way ANOVA with Tukey's *post hoc* test was used, *$p < 0.05$, **$p < 0.01$.

(**K**) C-Circle assay results visualized on slot blot. C-Circle assay with/without Phi29 polymerase (Pol+/Pol-) performed with DNA isolated from Control-sh or METTL3-KD U-2 OS cells. Box plots show the quantification of the blots, data are presented as signal/background. Data are shown as mean ± SD from two independent biological replicates. Unpaired *t*-test was used, * $p < 0.05$.

(**L**) Telomere DNA-FISH, for Control-sh or METTL3-KD U-2 OS cells. Telomere fluorescence intensity per cell is plotted. At least 110 cells were counted from three independent biological replicates. Unpaired *t*-test was used, **** $p < 0.0001$.

(**M**) Telomere DNA-FISH, in metaphase spreads of Control-sh or METTL3-KD U-2 OS cells. The white arrows indicate the telomere free ends. The percentage of Telomere Free Ends per metaphase is plotted. At least 30 metaphase spreads were counted from two independent biological replicates. Unpaired *t*-test was used, *** $p < 0.001$.

(**N**) Western blot to verify induction of HA-tagged dCasRx-METTL3$^{WT}$ and dCasRx-METTL3$^{MUT}$ by doxycycline (DOX). GAPDH is loading control.

(**O**) Western blot to verify induction of HA-tagged dCasRx-FTO$^{WT}$ and dCasRx-FTO$^{MUT}$ by DOX. GAPDH and Vinculin are loading controls.

(**P, left panel**) Slot blot with total RNA isolated from U-2 OS cells expressing dCasRx-FTO$^{WT}$ with either NTC or TERRA guide RNA. RNA was loaded in 3 different amounts 100, 200, and 400 ng. Blot probed with a DIG-labeled TERRA probe. TapeStation profile showing 18s and 28s rRNA served as a loading control. (**Right panel**) Bar graph shows the quantification of the blot normalized to loading control. Data are shown as mean ± SD from two independent biological replicates. Unpaired *t*-test was used.

(**Q, left panel**) Slot blot with total RNA isolated from U-2 OS cells expressing catalytically active CasRx-HA with either NTC or TERRA guide RNA. RNA was loaded in 2 different amounts 100 ng and 200 ng. Blot probed with DIG-labeled TERRA repeat-containing probe. TapeStation profile showing 18s and 28s rRNA served as a loading control. (**Middle panel**) Bar graph shows the quantification of the blot normalized to loading control. Data are shown as mean ± SD from two independent biological replicates. Unpaired *t*-test was used, *** $p < 0.001$. (**Right panel**) Western blot to verify induction of CasRx-HA by DOX. GAPDH is loading control.

(**R**) C-Circle assay results visualized on slot blot. C-Circle assay with/without Phi29 polymerase (Pol+/Pol-) performed with DNA isolated from U-2 OS cells expressing active CasRx-HA with either NTC or TERRA guide RNA.

(**S**) TERRA slot blot with total RNA isolated from U-2 OS cells treated with Actinomycin D for the indicated time points.

(**T**) TERRA foci (green) in U-2 OS cells treated with various METTL3 inhibitors for 6 h. Box plot shows the quantification of the number of TERRA foci per nucleus. At least 110 cells were counted from three independent biological replicates. Dunnett's multiple comparisons test was used, **** $p < 0.0001$. Scale bar is 10 µm.

(**U**) TERRA slot blot with total RNA isolated from U-2 OS cells treated with 10 µM of METTL3 inhibitor STM2457 for 6 h. DMSO served as a control for STM2457. TapeStation profile showing 18s and 28s rRNA served as a loading control. Bar graph shows the quantification of the blot normalized to loading control. Data are shown as mean ± SD from two independent biological replicates. Unpaired *t*-test was used.

(**V**) Cell cycle analysis. Histogram shows the cell cycle profile of cells treated with 10 µM of METTL3 inhibitor STM2457 for either 6 h or 48 h. DMSO treatment served as control. Data are shown as mean ± SD from two independent biological replicates. Unpaired *t*-test was used, ns - nonsignificant $p > 0.05$.

### Supplementary Figure S4

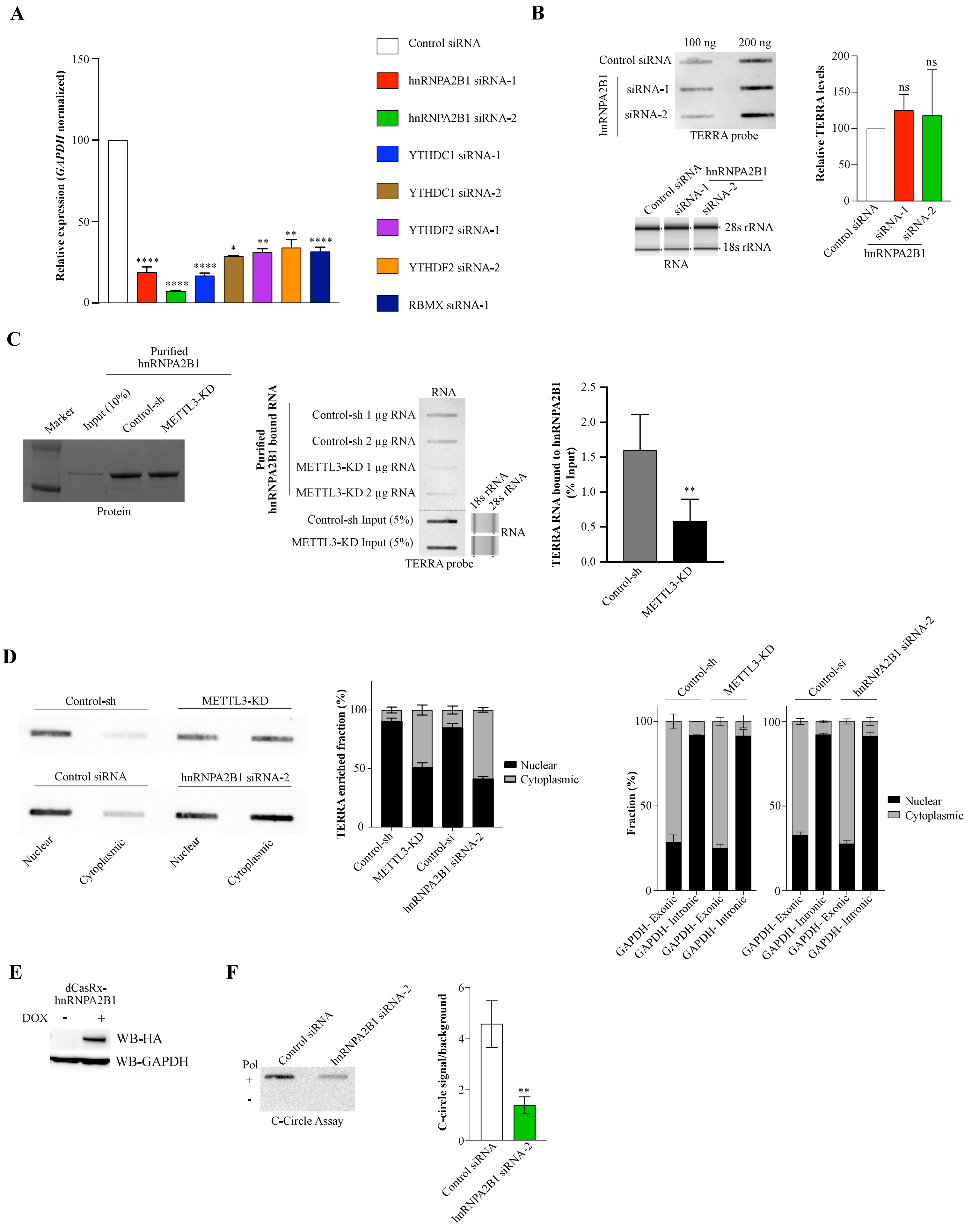

**Supplementary Figure S4:**

(**A**) Relative mRNA expression of m$^6$A reader proteins- *hnRNPA2B1*, *YTHDC1*, *RBMX*, and *YTHDF2* that were knocked down with siRNA. *GAPDH* was used to normalize the qPCR data. Data are shown as mean ± SD from two independent biological replicates. Dunnett's multiple comparisons test was used, * $p < 0.05$; ** $p < 0.01$; **** $p < 0.0001$.

(**B**) Slot blot with total RNA isolated from U-2 OS cells with either control or hnRNPA2B1 siRNA. RNA was loaded in 2 different amounts 100 ng and 200 ng. Blot probed with a DIG-labeled TERRA probe. TapeStation profile showing 18s and 28s rRNA served as a loading control. Bar graph shows the quantification of the blot normalized to loading control. Data are shown as mean ± SD from two independent biological replicates. Dunnett's multiple comparisons test was used.

(**C, left panel**) *In vitro* binding assay between total RNA isolated from Control-sh or METTL3-KD U-2 OS cells (2 different RNA concentrations) with purified His-tagged hnRNPA2B1. Left panel shows Coomassie staining of purified hnRNPA2B1 bound to beads. (**Middle panel**) Input RNA and hnRNPA2B1 bound RNA fraction on slot blot probed with DIG-labeled TERRA probe. Tapestastion showing 18s and 28s rRNA profiles of input RNA serves as loading control. (**Right panel**) Bar graph shows the quantification of the blots. Data are shown as mean ± SD from two independent biological replicates. Unpaired *t*-test was used, ** $p < 0.01$.

(**D, left panel**) Slot blot with RNA isolated post-nuclear-cytoplasmic fractionation from U-2 OS cells with either METTL3-KD or hnRNPA2B1 siRNA. Blot probed with a DIG-labeled TERRA probe. Data are shown as mean ± SD from two independent biological replicates. (**Right panel**) Relative level of intronic (nuclear marker) and exonic (cytoplasmic marker) GAPDH RNA in the nuclear and cytoplasmic fraction, measured by RT-qPCR. Data are shown as mean ± SD from two independent biological replicates. **(E)** Western blot to verify induction of HA-tagged dCasRx-hnRNPA2B1 by DOX. GAPDH is loading control.

(**F**) C-Circle assay results visualized on slot blot. C-Circle assay with/without Phi29 polymerase (Pol+/Pol-) performed with DNA isolated from Control or hnRNPA2B1siRNA treated U-2 OS cells. Box plots show the quantification of the blots, data are presented as signal/background. Data are shown as mean ± SD from two independent biological replicates. Unpaired *t*-test was used, ** $p < 0.01$.

#### Supplementary Figure S5

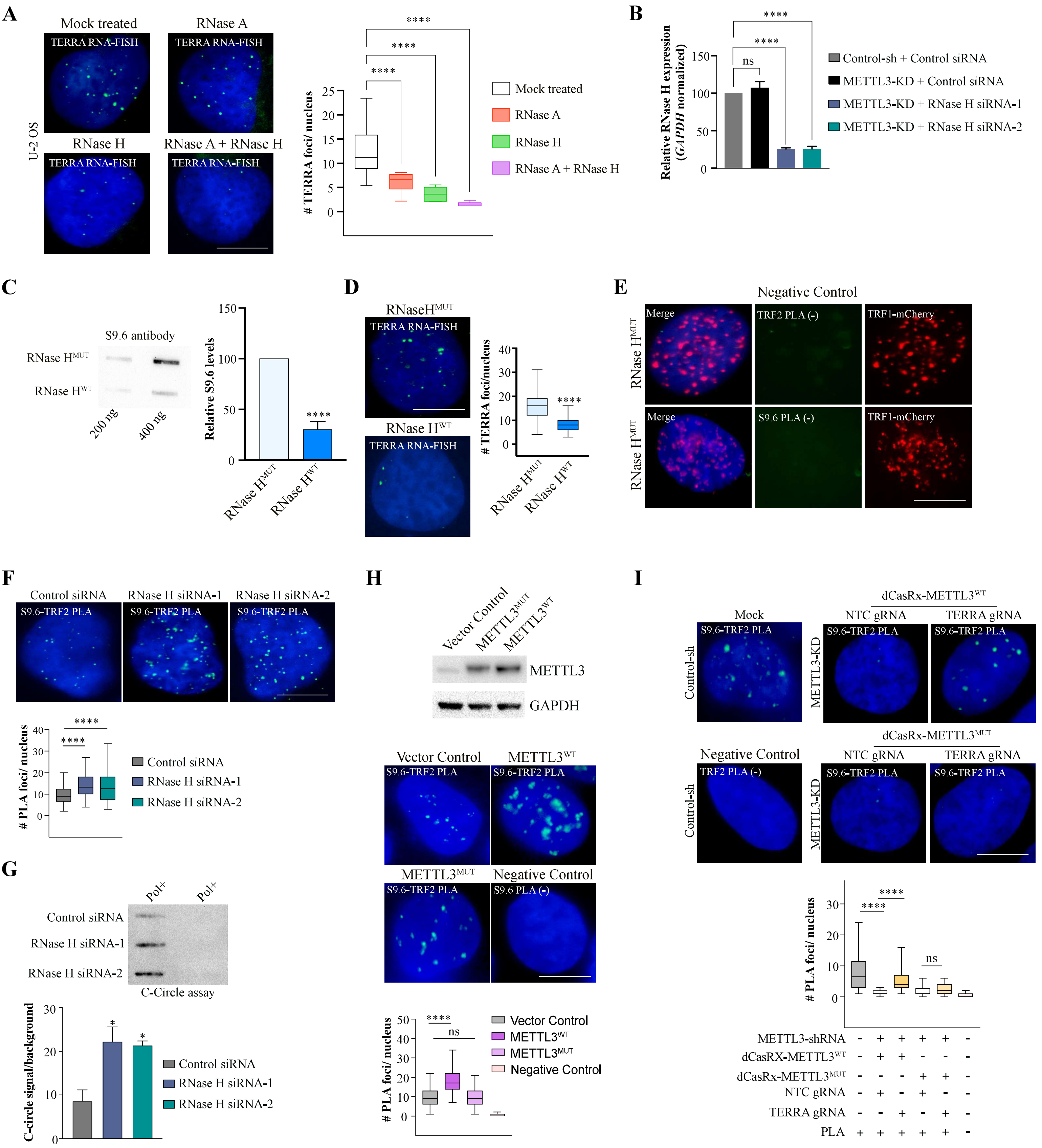

**Supplementary Figure S5:**

(**A**) TERRA RNA-FISH detecting TERRA foci (green) in U-2 OS cells exogenously treated with either mock, RNase A, RNase H, or with both RNase A and RNase H. Box plot shows the number of TERRA foci per nucleus. At least 75 cells were counted from three independent biological replicates. Dunnett's multiple comparisons test was used, **** $p < 0.0001$.

(**B**) Relative RNase H expression following siRNA-mediated KD. GAPDH was used to normalize the qPCR data. Data are shown as mean ± SD from two independent biological replicates. Dunnett's multiple comparisons test was used, **** $p < 0.0001$.

(**C**) Nucleic acid isolated from U-2 OS cells overexpressing RNase $H^{WT}$ or RNase $H^{MUT}$ were immobilized onto a nitrocellulose membrane and immunoblotted with S9.6 antibody. Bar graph is the quantification of the blot. Data are shown as mean ± SD from two independent biological replicates. Unpaired *t*-test was used, **** $p < 0.0001$.

(**D**) TERRA foci (green) in U-2 OS cells overexpressing RNase $H^{MUT}$ (top) or RNase $H^{WT}$ (bottom). Box plot shows the quantification of the number of TERRA foci per nucleus. At least 70 cells were counted from three independent biological replicates. Unpaired *t*-test was used, **** $p < 0.0001$.

(**E**) Background control for PLA with only TRF2 (left); or only S9.6 (right) antibody in RNase $H^{MUT}$ U-2 OS cells expressing TRF1-mCherry.

(**F**) PLA depicting the interaction of TRF2 with R-loop (S9.6 antibody) in Control or RNase H siRNA treated U-2 OS cells. Box plot shows the quantification of the PLA signal. At least 75 cells were counted from three independent biological replicates. Dunnett's multiple comparisons test was used, **** $p < 0.0001$.

(**G**) C-Circle assay results visualized on slot blot. C-Circle assay with/without Phi29 polymerase (Pol+/Pol-) performed with DNA isolated from Control or RNase H siRNA treated U-2 OS cells. Box plots show the quantification of the blots, data are presented as signal/background. Data are shown as mean ± SD from two independent biological replicates. Dunnett's multiple comparisons test was used, * $p < 0.05$.

(**H, top panel**) Western blot to verify overexpression of METTL3 in U-2 OS cells. GAPDH is loading control. (**Bottom panel**) PLA depicting the interaction of TRF2 with R-loop (S9.6 antibody) in U-2 OS cells overexpressing either Vector control, $METTL3^{WT,}$ or $METTL3^{MUT}$. PLA with only the S9.6 antibody served as a negative control. Box plot shows the quantification of the PLA signal. At least 70 cells were counted from three independent biological replicates. Dunnett's multiple comparisons test was used, **** $p < 0.0001$, ns- non-significance $p > 0.05$.

(**I**) PLA depicting the interaction of TRF2 with R-loop (S9.6 antibody) in METTL3-KD U-2 OS cells expressing shRNA resistant dCasRx-METTL3$^{WT}$/dCasRx-METTL3$^{MUT}$ with either NTC or TERRA guide RNA. Control-sh cells were used as a positive control and PLA with only TRF2 antibody served as a negative control. Box plot shows the quantification of the PLA signal in the conditions indicated. At least 75 cells were counted from three independent biological replicates. One-way ANOVA with Tukey's *post hoc* test was used, **** $p < 0.0001$, ns- nonsignificant $p > 0.05$. Scale bar is 10 μm.

### Supplementary Figure S6

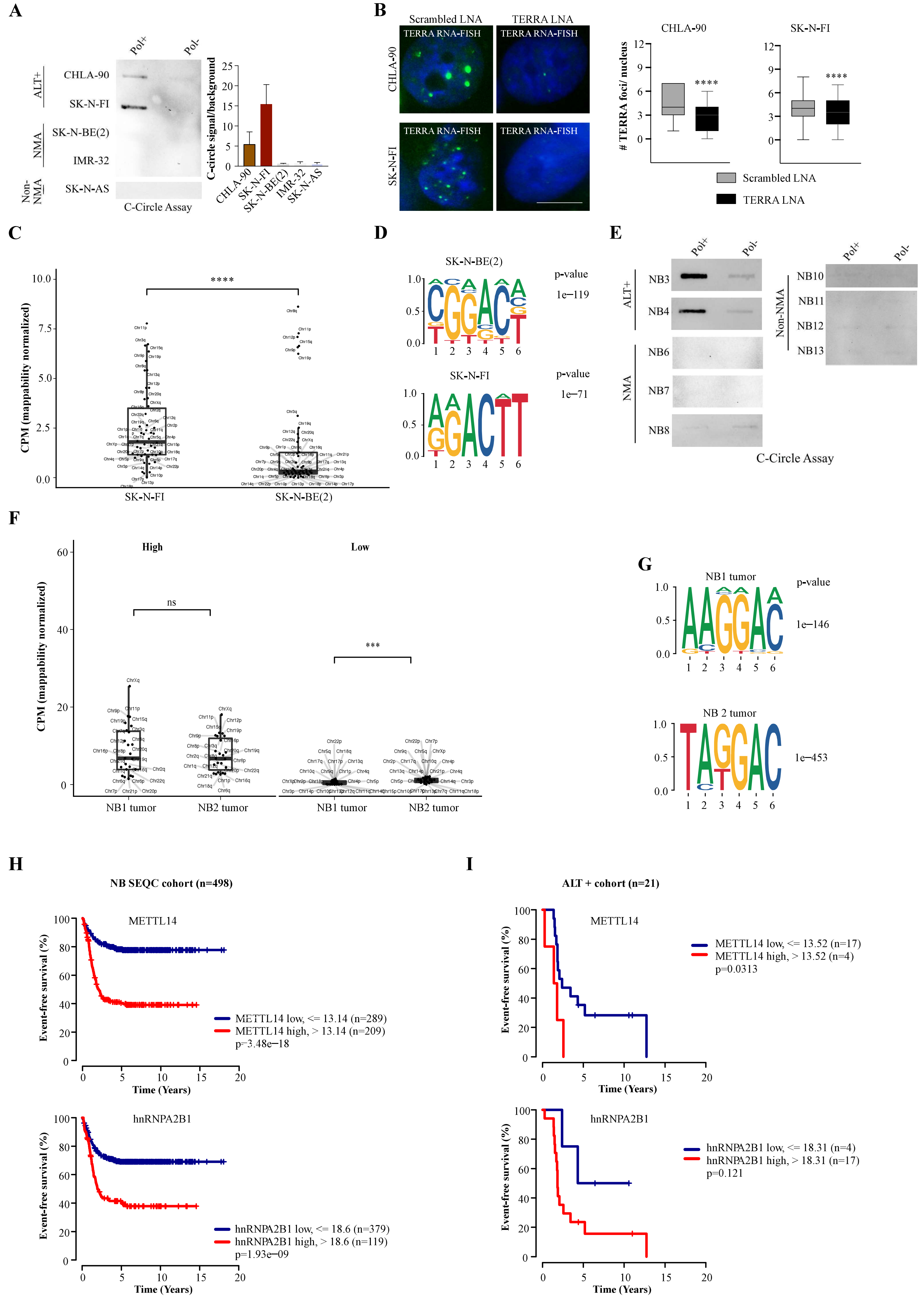

#### Supplementary Figure S6 continue

**J**

**ALT+ tumors**

| | METTL3 | METTL14 |
|---|---|---|
| NB14 | +++ | +++ |
| NB15 | +++ | +++ |
| NB16 | +++ | +++ |

**NMA- high risk tumors**

| | METTL3 | METTL14 |
|---|---|---|
| NB17 | + | + |
| NB18 | + | + |
| NB19 | + | + |
| NB20 | +++ | +++ |

**Low risk tumors**

| | METTL3 | METTL14 |
|---|---|---|
| NB21 | +++ | +++ |
| NB22 | - | - |
| NB23 | - | - |

+++ 100-50% of the tumor cells positive

++ 50-25% of the tumor cells positive

\+ Less than 25% of the tumor cells positive

\- Negative

**Supplementary Figure S6:**

(**A**) C-Circle assay (CCA) was performed on DNA from ALT+, NMA, and Non-NMA NB cells. Bar graph shows the quantification of the blots, data are presented as signal/background (Pol+/Pol-). Data are shown as mean ± SD from two independent biological replicates.

(**B**) TERRA foci (green) in ALT+ CHLA-90 (top) and SK-N-FI (bottom) cells treated with either Scrambled or TERRA LNA. Box plot shows the quantification of TERRA foci per nucleus. At least 70 cells were counted from three independent biological replicates. Unpaired *t*-test was used, **** $p < 0.0001$. Scale bar is 10 µm.

(**C**) Subtelomeric TERRA expression (CPM) in SK-N-FI and SK-N-BE(2) cells using mappability normalized RNA-seq data. Statistical significance was calculated using the two-sided paired *t*-test, **** $p < 0.0001$.

(**D**) Top enriched motif logo from *de novo* motif analysis of $m^6A$ peaks from SK-N-FI and SK-N-BE(2) $m^6A$ RIP-seq data.

(**E**) Representative C-Circle assay visualized on slot blot, performed with DNA from ALT+, NMA, and Non-NMA NB tumors.

(**F**) Subtelomeres are characterized as high and low on TERRA expression (CPM) in NB1 and NB2 tumors using mappability normalized RNA-seq data. Statistical significance was calculated using the two-sided paired *t*-test, *** $p < 0.001$.

(**G**) Top enriched motif logo from *de novo* motif analysis of NB1 and NB2 tumor $m^6A$ peaks.

(**H**) Event-free survival of NB patients (n=498, SEQC cohort) with either low (blue) or high (red) expression of METTL14 and hnRNPA2B1.

(**I**) Event-free survival of ALT+ NB patients (n=21) with either low (blue) or high (red) expression of METTL14 and hnRNPA2B1.

(**J**) Table summarizing results of immunohistochemistry analysis of METTL3 and METTL14 in NB tumors (ALT+, NMA- high risk and low risk).

### Supplementary Figure S7

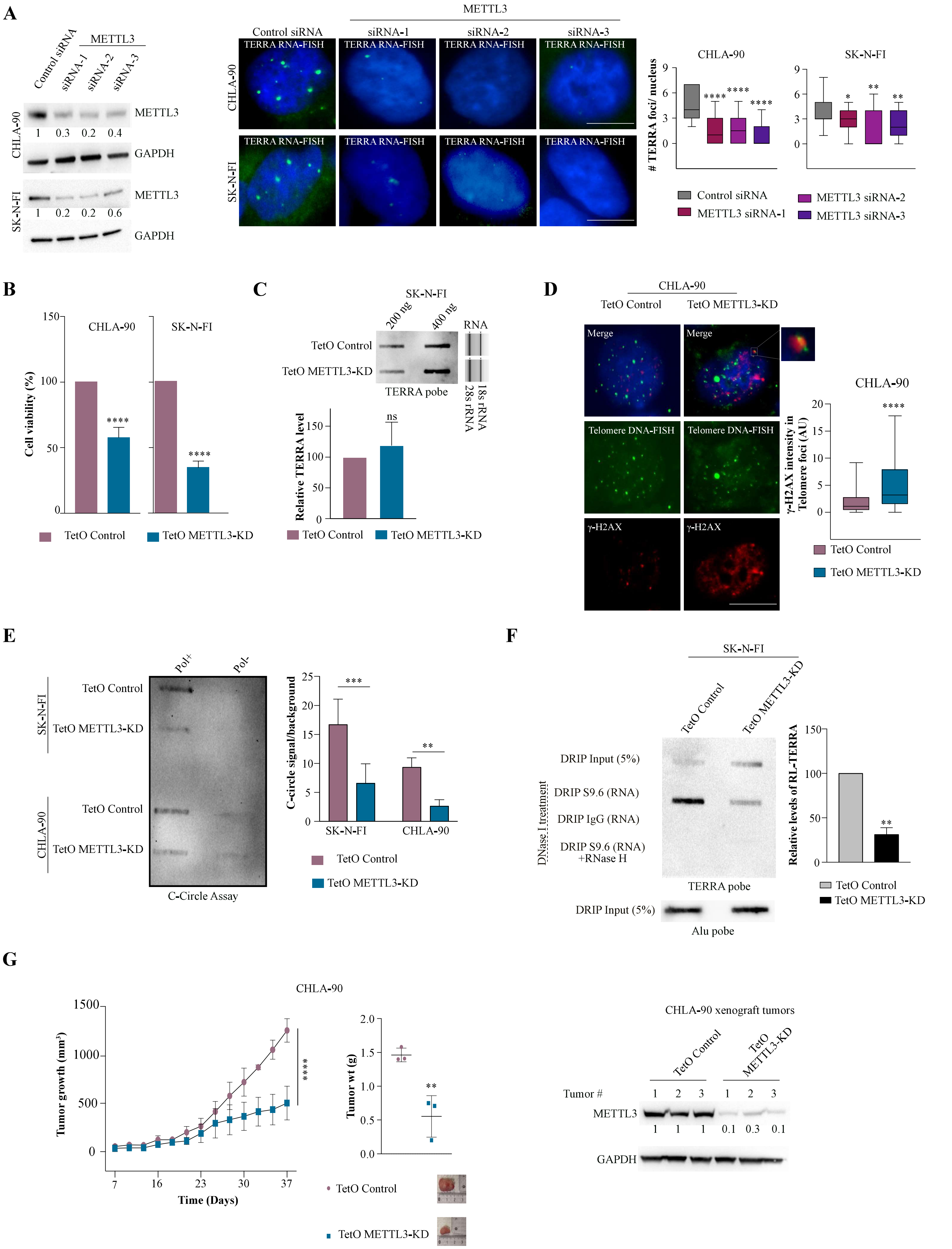

### Supplementary Figure S7 continue

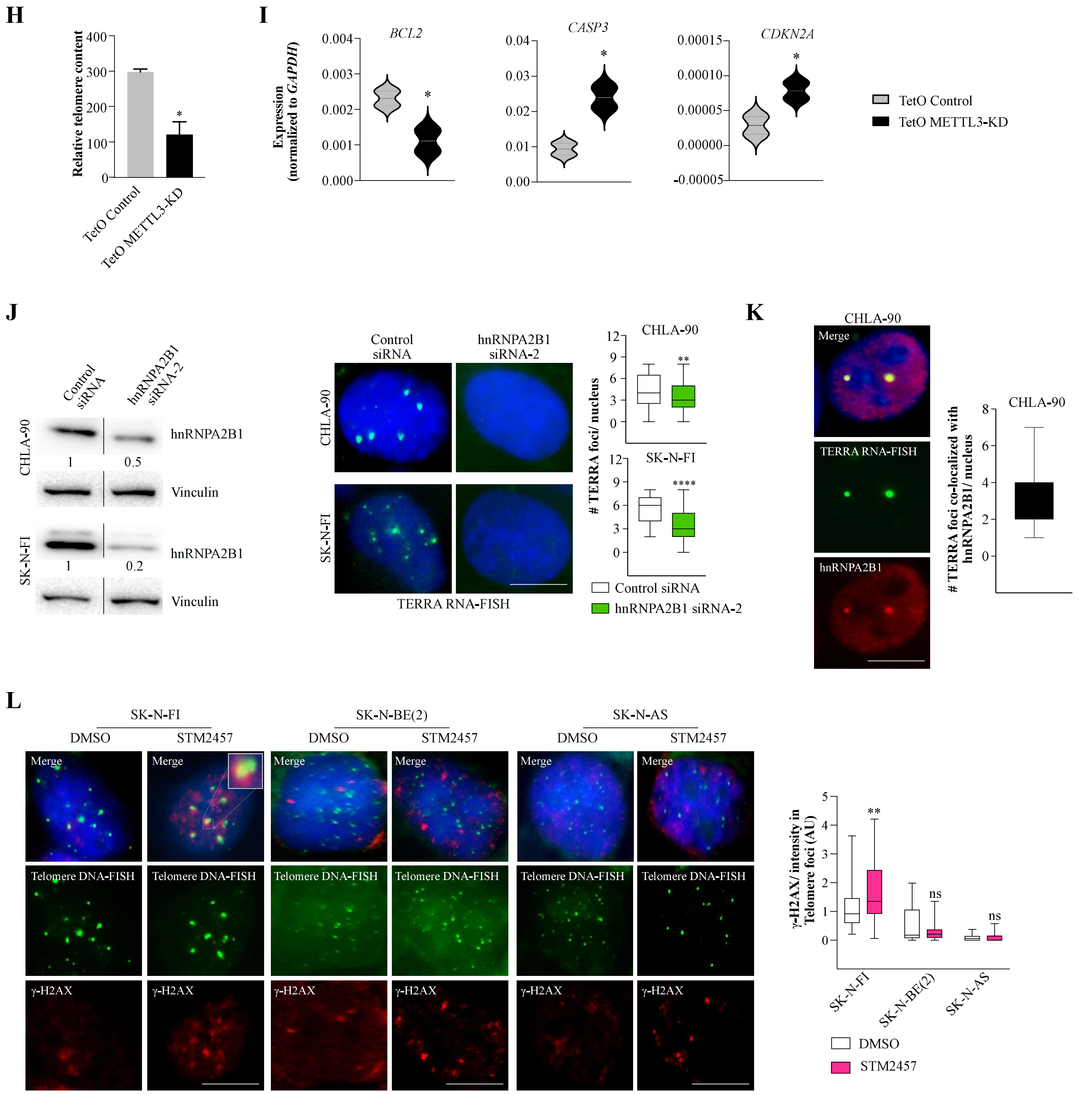

**Supplementary Figure S7:**

(**A**) TERRA foci following siRNA-mediated METTL3 depletion in ALT+ NB cells. (**Left panel**) Western blot for METTL3 level after siRNA transfection in CHLA-90 cells. (**Middle panel**) TERRA RNA-FISH detecting TERRA foci (green) in CHLA-90 and SK-N-FI cells after METTL3-KD as indicated. (**Right panel**) Box plots show the quantification of TERRA foci per nucleus. At least 50 cells were counted from two to three independent biological replicates. Dunnett's multiple comparisons test was used, * $p < 0.05$; ** $p < 0.01$; **** $p < 0.0001$.

(**B**) Bar graph displaying the effect on cell viability after conditional KD of METTL3 (6 days post DOX treatment) in CHLA-90 and SK-N-FI TetO METTL3-KD cells. Data normalized to control and shown as mean ± SD. Unpaired *t*-test was used, **** $p < 0.0001$.

(**C**) Slot blot with total RNA isolated from SK-N-FI cells with TetO Control or METTL3-KD. RNA was loaded in 2 different amounts 200 ng and 400 ng. Blot probed with a DIG-labeled TERRA probe. TapeStation profile showing 18s and 28s RNA served as a loading control. Bar graph showing quantification normalized to loading control. Data are shown as mean ± SD from two independent biological replicates. Unpaired *t*-test was used. ns- nonsignificant $p > 0.05$.

(**D**) Accumulation of γ-H2AX (red) over telomere after 48 h of DOX induction in CHLA-90 cells with TetO METTL3-KD. Telomere (green) was detected by telomere DNA-FISH. Box plot shows the γ-H2AX intensity in the telomere. At least 110 cells were counted from three independent biological replicates. Unpaired *t*-test was used, **** $p < 0.0001$.

(**E**) C-Circle assay results are visualized on a slot blot. DNA isolated from CHLA-90 and SK-N-FI cells with TetO METTL3-KD (48 h DOX) was subjected to a C-Circle assay with/without Phi29 polymerase (Pol+/Pol-). Bar graph shows the quantification of the blots. Data are shown as mean ± SD from two independent biological replicates. Unpaired *t*-test was used, ** $p < 0.01$; *** $p < 0.001$.

(**F**) DRIP followed by slot blot assay and probed with DIG-labeled TERRA probe in TetO Control or METTL3-KD SK-N-FI cells, 72 h post-DOX induction. Experiments performed with IgG antibody or nuclei acid pre-treated with RNase H served as a control. Input samples probed with Alu probe served as a loading control. Bar graph shows the quantification of the blots. Data are shown as mean ± SD from two biological replicates. Unpaired *t*-test was used, ** $p < 0.01$.

(**G, Left panel**) Line graph showing average tumor growth ($mm^3$) of CHLA-90 cells (TetO Control and METTL3-KD) derived xenografts. Data are shown as mean ± SD. Sidak's multiple comparisons test was used, **** $p < 0.0001$. (**Middle panel**) Scatter plot showing tumor weight post necropsies of xenografts derived from CHLA-90 cells with either TetO Control or METTL3-KD. Data are shown as mean ± SD. Unpaired *t*-test was used, ** $p < 0.01$ (n=3). One representative image of the tumor is shown per condition. (**Right panel**) Western blot confirms the METTL3-KD in xenograft tumors and GAPDH acted as a loading control.

(**H**) Bar plot shows the relative telomere length measured by qPCR in SK-N-FI cells derived xenografts with either TetO Control or METTL3-KD. Data normalized to *IFNB1* and shown as mean ± SD (n=4). Unpaired *t*-test was used, * $p < 0.05$.

(**I**) Expression of *BCL2, CASP3,* and *CDKN2A* in xenograft derived from SK-N-FI cells with either TetO Control or METTL3-KD. RT-qPCR data normalized to *GAPDH* are shown as mean ± SD (n=4). Unpaired *t*-test was used, * $p < 0.05$.

(**J, left panel**) Western blot to verify the hnRNPA2B1-KD in ALT+ NB cells. Vinculin was used as a loading control. (**Right panel**) TERRA foci (green) in CHLA-90 (top) and SK-N-FI (bottom) cells after transient KD of hnRNPA2B1. Box plot shows the quantification of TERRA foci per nucleus. At least 60 cells were counted from three independent biological replicates. Unpaired *t*-test was used, ** $p < 0.01$; **** $p < 0.0001$.

(**K**) TERRA RNA-FISH detecting TERRA foci (green) combined with hnRNPA2B1 (red) IF in CHLA-90 cells. Box plot shows the quantification of TERRA and $m^6A$ overlap. At least 35 cells were counted from two independent biological replicates.

(**L**) Telomere (green) DNA-FISH and γ-H2AX (red) IF in ALT+ (SK-N-FI), NMA [SK-N-BE(2)] and Non-NMA (SK-N-AS) cells treated with METTL3 inhibitor, STM2457 for 48 h. Box plot showing γ-H2AX intensity in the telomere. At least 70 cells were counted from two to three independent biological replicates. Unpaired *t*-test was used, ** $p < 0.01$, ns-nonsignificant $p > 0.05$. Scale bar is 10 µm.

#### Supplementary methods

##### *Gene silencing and overexpression:*

Lentiviral based vectors were packaged following instructions from Addgene using packaging plasmids psPAX2 (12260) and pMD2.G (12259). To generate stable cells antibiotic selection was performed depending on the type of vector used following lentiviral transduction. To generate METTL3-KD cells U-2 OS and ALT+ NB cell lines were selected with media containing Puromycin (1 µg/ml). In NB cells DOX (200 ng/ml) was used to induce METTL3-KD. To generate CasRx/dCasRx-FTO$^{WT}$/dCasRx-FTO$^{MUT}$/dCasRx-METTL3$^{WT}$/dCasRx-METTL3$^{MUT}$/dCasRx-hnRNPA2B1 and gRNA expressing U-2 OS cells, firstly CasRx /dCasRx-FTO/METTL3/hnRNPA2B1 containing lentivirus were transduced followed by selection with Puromycin (1 µg/ml). Next, the puromycin selected cells were further transduced with gRNA lentivirus and selected with Blasticidin (5 µg/ml). To induce CasRx/dCasRx-FTO/METTL3/hnRNPA2B1 expression, cells were cultured in DOX (200 ng/ml) containing media. RNase H WT and mutant overexpressing vectors were purchased from Addgene (111906 and 111905). To generate stable RNase H$^{WT}$/RNase H$^{MUT}$ cell lines corresponding plasmids were transfected using Lipofectamine 3000 reagent (Thermo Fisher Scientific) following manufacturer instruction followed by selection with media containing Hygromycin (400 µg/ml). METTL3$^{WT}$ (160250, Addgene) and METTL3$^{MUT}$ (160251, Addgene) plasmids were transfected into U-2 OS cells using Lipofectamine 3000 reagent. METTL3 siRNAs were transfected to induce transient knock-down in both U-2 OS and ALT+ NB cells using RNAiMAX (Thermo Fisher Scientific) reagents following the manufacturer's instructions.

##### *RNA and DNA isolation*

*From Cell-lines*

RNA was isolated from the cell line using TRIzol reagent (15596026, Thermo Fisher Scientific) and Direct-zol RNA Miniprep (R2050, ZYMO research) or using ReliaPrep RNA Cell Miniprep system (Z6011, Promega), according to the manufacturer's instruction. The DNase I treatment step was performed mandatorily, and the RNA was eluted in RNase/DNase-free water. DNA was isolated from cells using the Wizard Genomic DNA purification Kit (Promega) according to the manufacturer's instructions.

*From Tumor material*

Tumor tissue was collected after written or verbal informed consent was obtained from parents/guardians according to ethical permits approved by the local ethics committees

(Karolinska Institute and Karolinska University Hospital Research Ethics Committee, registration number 2009/1369-31/1 and 03-736). RNA was extracted from frozen tumor samples using All Prep DNA/RNA/Protein Mini Kit (Qiagen) according to the manufacturer's instructions. Genomic DNA was extracted from fresh/frozen tumors or blood using the DNeasy Blood & Tissue kit (Qiagen, Hilden, Germany) according to the manufacturer's instructions. This study was conducted per the Declaration of Helsinki.

***Processing of m$^6$A RIP-seq data***

Raw Illumina short-reads data was obtained from SMARTER-Stranded Total RNA-Seq Kit v2 sequencing and data were first processed using Trim Galore v0.6.6 (https://www.bioinformatics.babraham.ac.uk/projects/trim_galore/) with a minimum length threshold of 20bp. Quality control of Illumina short-reads was performed using FastQC v0.11.9 (https://www.bioinformatics.babraham.ac.uk/projects/fastqc/) with default parameters (-q 20). Trimmed reads were mapped with HISAT2 v2.2.1 (1) preserving strand information (-U –rna-strandness R) using the Telomere-to-Telomere (T2T) human reference genome assembly (T2T-CHM13 v1.1) (2). The Escherichia coli K12 genome was concatenated to the T2T reference assembly for further spike-in normalization. To control the systematic variation across m$^6$A RIP experiments, the amount of spiked-in bacterial RNA was estimated by counting the total number of reads uniquely mapping to the E. coli K-12 reference genome using Sambamba v0.7.1 (3). The E. coli spike-in bacterial counts were further used to calculate scaling factors for each batch of m$^6$A RIP-seq samples. Duplicate reads were labeled using MarkDuplicates from Picard v2.23.4 (http://broadinstitute.github.io/picard/). Alignments were downsampled according to calculated spike-in scaling factors using DownsampleSam from Picard (--strategy HighAccuracy). Mapping files were separated by strand after duplicate removal using Sambamba v0.7.1. For U-2 OS cells two replicates were pooled to obtain enough reads and for neuroblastoma RNA samples one replicate was used per condition.

***Analysis of m$^6$A RIP-seq and ChIP-seq data***

Peak calling was performed using MACS v2.2.6 (https://github.com/macs3-project/MACS) on m$^6$A RIP and input alignments normalized to spike-in content, with parameters --no-model, --kee*p*-dup auto, --call-summits, –extsize 75, q-value cutoff of 0.05 and effective genome size according to the T2T-CHM13 v1.1 UCSC table browser. Motif analysis on m$^6$A peaks was carried out using findMotifsGenome.pl from HOMER v4.11 (4). m$^6$A RIP coverage tracks were generated using bamCoverage from deepTools v3.3.2 (5).

Public CTCF and RNA Pol II data reported by Deng et al. 2012 (6) were obtained from GEO (GSE1962, GSE19484). Visualization of $m^6A$ RIP coverage tracks data was done in R using the rtracklayer (7) and ggplot2 packages (https://www.R-project.org). Telomere coordinates, GC% content, and CpG island coordinates were retrieved from the T2T CHM13 v1.1 reference genome hub (http://t2t.gi.ucsc.edu/chm13/hub/t2t-chm13-v1.1). Telomeric repeat tracks were generated using scanMotifGenomeWide.pl from HOMER v4.11 and bedToBigBed v2.8 from UCSC utilities (8).

***Processing of direct-RNA Nanopore sequencing data***

Direct RNA Oxford Nanopore Technology (ONT) sequencing data was base-called using Guppy v5.0.16 (flowcell FLO-MIN106, kit SQK-RNA002). Quality control was performed with PycoQC (https://github.com/a-slide/pycoQC) and NanoPlot (https://github.com/wdecoster/NanoPlot) on base-called data. ONT reads were mapped to the T2T-CHM13 v1.1 reference genome using the latest version of Minimap (v2.24) (-ma*p*-ont -uf -k14) optimized for mapping long reads to highly repetitive regions (9,10).

***Analysis of ONT data***

Identification of $m^6A$ modifications from direct RNA ONT data was performed on one replicate per condition following the xPore pipeline (https://xpore.readthedocs.io/) (11). xPore allows the identification of $m^6A$ under multiple conditions without the need for unmodified control samples, enabling the quantitative estimation of modification rates from Nanopore sequencing data. Briefly, raw Nanopore signal-level events were aligned to the T2T CHM13 v1.1 reference transcriptome using Nanopolish eventalign v0.12.0. Processed ONT data were further analyzed with xPore (default parameters; minimum read count 5, pre-filtering t-test method, 500 maximum iterations) to estimate the genome-wide RNA modification rates for each independent sample and the differentially modified sites between control and METTL3-KD conditions. A transcriptome assembly was performed using StringTie2 (12) on Nanopore reads, and remapped reads were used with xPore to refine the search for $m^6A$ modifications at chromosome ends. Predicted $m^6A$ sites were selected from differentially modified positions with a base A within the NNANN context with a significance $p$-value $< 0.05$ and a differential modification rate (DMR) above zero.

***Analysis of TERRA RNA pulldown***

Reads from TERRA and Luciferase RNA pulldown were mapped to the T2T-CHM13 v1.1 genome assembly and quantified after duplicate removal. To further analyze the reads located within genomic repeat elements and TERRA repeats, the coordinates of genomic repeat elements were retrieved from the UCSC genome browser (http://t2t.gi.ucsc.edu/chm13/hub/t2t-chm13-v1.1/rmsk/rmsk.bigBed) and concatenated with TERRA repeat coordinates obtained using SeqKit locate and BEDTools (13,14). Mapped reads overlapping with TERRA and genomic repeat element coordinates were quantified using BEDTools multicov function and normalized to RPKM values to account for differences in library sizes and repeat lengths. RPKM values were further normalized to their overlapping mappability window scores obtained using BEDTools map and intersect functions, to account for different mappability likelihoods for each repeat coordinate. Further, the ratio of TERRA RNA pulldown versus Luciferase was calculated from the mappability-normalized RPKM values for the LINE, SINE, LTR, Satellite elements and TERRA repeats, respectively, to analyze the specificity of TERRA RNA distribution within each repeat family.

***Sequential RNA-FISH and IF***

For sequential IF staining followed by RNA-FISH as described in the main method. Cells were fixed using 4% formaldehyde for 10 min followed by 2x washes with PBS and stored in 70% ethanol at 4°C until further use. Coverslips were rinsed with PBS and cells were permeabilized for 10 min using 0.25% Triton X-100 in 1x PBS at RT. Cells were then washed using PBS-T (0.1% Tween-20 in 1x PBS) and blocked in 3% BSA in PBS-T for 30 min. Primary antibodies were diluted in 3% BSA in PBS-T and incubated for 2 h after blocking. 3x5 min washes were performed using PBS-T and incubated for 1 h with secondary antibodies diluted in 3% BSA in PBS-T. 3x5 min washes were performed using PBS-T followed by 10 min fixation by 4% formaldehyde. Cells were rinsed with PBS twice, permeabilized with 0.25% Triton X-100 in 1x PBS for 5 min followed by a wash with PBS. TERRA RNA-FISH was performed as described above and proceeded for imaging. Details of antibodies are provided in Supplementary Table S1.

***Simultaneous RNA-FISH and $m^6A$ IF staining***

Simultaneous TERRA FISH and $m^6A$ IF were performed as described before (15) with the following modifications. Cells were fixed in MeOH at -20°C for 15 min and TERRA RNA FISH was performed as described above with some modification. After hybridization, during 2x30 min washing steps, cold wash buffer A (described above) was added to coverslips and incubated at 37°C. This step allows the refolding of rRNAs to reduce the nuclear RNA signal coming from rRNAs as described by Fu et al (15). Coverslips were then washed with wash buffer B (described above) for 2-3 min and then rinsed with 1x PBS. Coverslips were incubated in wash buffer A containing 10% formamide for 5-10 min and followed by blocking for 1 h (3% BSA in PBS-T) and the primary antibody against $m^6A$ was incubated for 2 h at RT in 3% BSA in PBS-T. 3x5 min PBS-T washes were done and the secondary antibody was incubated for 1 h at RT. Coverslips were washed 3x5 min with PBS-T and mounted. Imaging was done using EVOS M7000 microscope.

***Western blot***

Cell lysates were prepared using RIPA buffer (R0278, Sigma-Aldrich,) and total protein content was quantified using Pierce BCA Protein Assay Kit (23225, Thermo Fisher Scientific) as per manufacturer's instructions. Equal amounts and volume of samples were resolved by SDS- PAGE on NuPAGE Bis-Tri Gels (4-12%) (Thermo Fisher Scientific), followed by transfer onto 0.2 µm Nitrocellulose membrane using Trans-Blot Turbo Transfer System (Bio-Rad). The membrane was blocked for 1 h at RT with a 5% blocking solution of non-fat dried milk in PBS-T.

The membrane was incubated with the primary antibody in a 5% blocking solution overnight at 4°C. Immunoblotting was continued the next day starting with 3x PBS-T washes followed by incubation with respective secondary antibodies for 1 h at RT, and then 3X PBS-T washes. Proteins were detected with SuperSignal West Pico PLUS Chemiluminescent Substrate (34579, Thermo Fisher Scientific) using the ChemiDoc XRS+ system (Bio-Rad). ImageLab software was used to quantify the bands. Antibodies used in Western blot analysis are provided in Supplementary Table S1.

***Proliferation assay (MTT)***

5,000 cells/well were seeded in a 96-well plate to assess the proliferation of cells after METTL3 inhibitor STM2457 (10 µM) at indicated time points. CellTiter 96 Non-Radioactive Cell Proliferation Assay kit (G4000, Promega) was used to determine cell growth and the manufacturer's instructions were followed. Absorbance was measured using a microplate reader Infinite 50 (Tecan, Austria).

***Colony formation assay***

10,000 cells/well were seeded in a 6-well plate and allowed to attach overnight. DOX (200 ng/ml) was added to the media for induction of METTL3-KD and cells were left to grow sparsely for 14 days with induction media which was replaced after every 3-4 days. Cells were then washed 1x with PBS, fixed with 4% formaldehyde for 10 min at RT, washed 1x with PBS, and then stained using 1% crystal violet solution at RT for 2 h. The plate was carefully washed and allowed to dry. The image was taken using the ChemiDoc XRS+ system (Bio-Rad).

##### *In vitro METTL3/METTL14 methyltransferase assay*

Methyl-transferase assay with RNA oligos (1 µm DRACH motif containing and TERRA repeat containing oligo, Supplementary Table S1) was performed with recombinant METTL3/METTL14 enzyme complex (31570, Active motif) according to manufacturer's instructions, with 10 nM S-Adenosyl methionine (SAM) as methyl group donor. Oligos without enzyme complex were used as a negative control. To detect if the RNA oligos underwent $m^6A$ modification LC-MS/MS-based quantification of $m^6A$ was done, as previously described (16). In brief oligos, post methyltransferase assay were digested by Nuclease P1 (N8630, Sigma-Aldrich) followed by treatment with phosphatase (M0289S, NEB). The sample was then filtered (0.22 µm pore size) and directly injected into the LC-MS. We made triplicate injections of the RNA oligos (both experimental and negative controls) and estimated the ratio of $A/m^6A$. LC-MS/MS profiles were monitored using the parallel reaction-monitoring (PRM) mode for: m/z 268.0–136.0, and m/z 282.0–150.1 which corresponds to A and $m^6A$ respectively as described before (16). DRACH oligos served as a positive control.

##### *METTL3 Chromatin Immunoprecipitation (ChIP) slot blot/ qPCR*

ChIP with METTL3 antibody was performed as described before (17). U-2 OS cells were fixed with formaldehyde (1% final concentration) for 15 min at RT. Fixed cells were washed with cold PBS, lysed and the chromatin was sheared using a Bioruptor (Diagenode) until an average fragment size range of 200-500 bp was achieved. Immunoprecipitation of solubilized chromatin was carried out with 3 µg of METTL3/ g-H2AX/ IgG antibody overnight at 4°C. The immunoprecipitated complex was captured using a mix of Protein A and G Dynabeads (Invitrogen), washed and RNase A treated before eluting the DNA. The ChIP DNA was denatured at 95°C for 5 min before assaying on slot blot using a telomere probe.

For the ChIP qPCR, the input DNA and the immunoprecipitated DNA were directly subjected to qPCR with primers for telomere ends (Supplementary Table S1).

##### *hnRNPA2B1 RIP and hnRNPA2B1-$m^6A$ Re-RIP*

Cells were harvested and fixed for 10 min with formaldehyde (1% final concentration) and quenched with Glycine (0.125 M, final concentration). Fixed cells were lysed and nuclei were isolated by incubating cells in nuclei prep buffer (100 mM Tris pH 7.4, 10 mM Potassium acetate, 10 mM Magnesium acetate, 1% IGEPAL-CA630, 1mM DTT, Protease inhibitor, and

RNase inhibitor) for 10 min in cold followed by centrifugation at 2500 g. the pellet was resuspended in cold RIPA buffer (50 mM Tris pH7.4, 150 mM NaCl, 0.5% Sodium Deoxycholate, 0.2% SDS, 1% IGEPAL-CA630, Protease inhibitor, and RNase inhibitor) and sonicated for 10 cycles (30 secs on 30 secs off) on Bioruptor (Diagenode) and cleared lysate was used for RIP with 3 μg of hnRNPA2B1 (A73256, Gentek,) and IgG (sc-2027, Santa Cruz) antibodies. The RNA-protein-antibody complex was captured using protein A/G magnetic beads (Thermo Fisher Scientific). Magnetic beads were washed with low salt buffer (1x PBS, 0.1% SDS, and 0.5% IGEPAL-CA630) and high salt buffer (5x PBS, 0.1% SDS, and 0.5% IGEPAL-CA630) before eluting in elution buffer (10 mM Tris pH 7.4, 100 mM NaCl, 1 mM EDTA, 0.5% SDS) with Proteinase K followed by RNA extraction using TRIzol reagent. The RNA isolated after hnRNPA2B1/IgG RIP was either directly used on a slot blot assay or subsequently used as input for Re-RIP with $m^6A$ antibody. $m^6A$ RIP was performed as described in the method section. The final Re-RIP RNA was directly assayed on slot blot.

***Purification of hnRNPA2B1 protein and in vitro binding assay***

His-tagged hnRNPA2_WT (98662, Addgene) was transformed in Rossetta DE3 bacterial strain. Cells were grown overnight and further diluted to 0.1 OD 600 nm and grown to about 0.6 OD 600 nm at 37°C and then cells were induced with 0.3 mM IPTG at 18°C for 14-16 h. Cells were pelleted and dissolved in 25 mM HEPES pH 7.5, 150 mM NaCl, 20 mM KCl, and 20 mM $MgCl_2$ along with 1x Protease inhibitor cocktail. Cells were lysed at 4°C in the presence of lysozyme followed by brief sonication. Cell debris were removed by spinning cell lysate at 13000 rpm for 30 min at 4C and cleared lysate were loaded onto Co+ NTA resin for binding of His-tagged hnRNPA2B1 protein and resin were washed with 20 mM imidazole containing lysis buffer without 1x Protease inhibitor cocktail to remove non-specific proteins and His-tagged protein were eluted in 300 mM imidazole containing lysis buffer and eluted protein were further dialyzed in lysis buffer and proteins were stored in -80°C for further use.

The hnRNPA2B1 binding with RNA was performed following the protocol as described before (18) with few modifications. Total RNA from Control-sh or METTL3-KD U-2 OS cells were used instead of *in vitro* synthesized RNA. The RNA bound protein complex was captured by HisPur Cobalt Resin (89965, Thermo Fisher Scientific) and TERRA enrichment was checked by slot blot assay as described above.

***Analysis associated with detection of Chromosomal aberration and ATRX status in NB Swedish cohort***

Tumor samples were analyzed with Applied Biosystems CytoScan HD array (Thermo Fisher Scientific, Waltham) according to the experimental procedure previously described (19,20). The arrays detect both copy number alterations and allele-specific information at high resolution. For primary data analysis, GDAS software (Thermo Fisher Scientific) was used. Chromosome Analysis Suite (ChAS v.3.3; Thermo Fisher Scientific) was used for the generation of genomic profiles and determination of chromosomal aberration, MYCN amplification, and ATRX status. For Whole Genome Sequencing (WGS) DNA from tumor material and matched constitutional DNA were subjected to sequencing and bioinformatical handling as described previously (20).

***C-Circle assay***

50 ng of genomic DNA was digested with 4 U/µg HinfI and RsaI restriction enzymes and 25 ng/µg RNase A for 1 h at 37°C (prepare every sample twice to include no polymerase as negative control). Digested DNA was incubated with 100ug/ml Albumin, 1 mM each dATP, dGTP, and dTTP, 0.1% Tween, 1x Φ29 Buffer, and 7.5 U Φ29 DNA polymerase (NEB) at 30°C for 8 h then 65°C for 20 min. The reaction products were slot blotted onto a 6x SSC soaked Biodyne B nylon membrane (GE Healthcare) for quantification and UV-crosslinking the DNA onto the membrane at 0.120 Joules. Prehybridization was done using DIG Easy Hyb (11603558001, Roche) for 30 min at 37°C, and after adding 25 ng/ml DIG-labeled telomere probe: 5′-$(TAACCC)_5$-DIG (Sigma-Aldrich) at 37°C overnight with rotation. For hybridization, the DIG Wash and Block Buffer set (11585762001, Roche) was used according to the manufacturer's instructions. The membrane was incubated with an anti-DIG antibody conjugated with alkaline phosphatase (1:20,000, 11093274910, Roche) for 1 h and CDP-Star Chemiluminescent substrate (C0712, Sigma-Aldrich) to detect the C-circles. Images were taken using high-resolution chemiluminescence (ChemiDoc, Bio-Rad)

***FISH and IF in metaphase chromosomes***

To prepare metaphase chromosome spreads, cells were treated with 1 µg/µl KaryoMAX colcemid solution for 3 h to induce metaphase arrest. The cells were then harvested, resuspended in 0.075 M KCl at 37°C for 10 minutes, and fixed using a 3:1 mixture of ice-cold methanol/acetic acid. The fixed cells were spread onto glass slides and air-dried at room temperature.

For DNA-FISH on metaphase slides, a similar procedure was followed as described in the method for Telomere DNA-FISH and IF. Instead of performing IF we directly proceed to the DNA-FISH procedure.

For TERRA FISH followed by $m^6A$ IF on metaphase slides, a similar procedure was followed as described in simultaneous RNA-FISH and $m^6A$ IF staining. Metaphase slides were fixed with 4% formaldehyde for 10 min at RT. After washing twice with PBS, the slides were dehydrated with ethanol gradient for 3 min and air dried for 5 min at RT followed by TERRA FISH. As a control, metaphase slides were treated with a combination of RNase A (100 µg/ml, Sigma-Aldrich) and RNase H (10 units, NEB) or PBS for 1 h at 37°C prior to the procedure.

##### *Nuclear and cytoplasmic RNA preparation*

U-2 OS cells were harvested by trypsinization and washed once with cold PBS. One million cells were resuspended in 175µl of RLN1 buffer (50 mM Tris-HCl; pH 8, 140 mM NaCl, 1.5 mM MgCl2, 0.5%v/v NP-40, 40 U RNasin® Ribonuclease inhibitors) and incubated for 5 mins on ice followed by centrifugation in 300 g (1800 RPM). Transfer the supernatant to the new tube carefully without disturbing the pellet and this is considered as cytoplasmic fraction. Pellet contains a nuclear fraction. RNA was isolated using TRIzol reagent (15596026, Thermo Fisher Scientific) and Direct-zol RNA Miniprep (R2050, ZYMO research). Terra level in the nuclear and cytoplasmic fraction was checked by TERRA slot blot assay. Quality of cytoplasmic and nuclear RNA preparation was verified using *GAPDH* exonic and *GAPDH* intronic primers respectively.

##### *Telomere content measurement by qPCR*

Relative Telomere content was measured by following a previously published protocol (21). In brief genomic DNA was isolated from SK-N-FI xenograft from control and METTL3-KD cells. Genomic DNA was used to perform qPCR with either telomere specific primers or with *IFNB1* primers (reference gene). Relative telomere content was measured across the samples after normalization with reference gene *IFNB1*.

##### *Analysis of neuroblastoma next-generation sequencing data from a German cohort*

Copy number gains were estimated from a combination of published neuroblastoma paired-end WGS (22) paired-end WES (23), and aCGH (24) datasets. ALT and MYCN status of the patients were taken from sample overlaps with Roderweiser et al. (25), sequence alignment to reference GRCh37/hg19 was done using BWA mem (v.0.7.13-r1126; https://github.com/lh3/bwa) and copy number analysis was performed with Sclust (26).

Gene expression analysis was performed with the RNA-Seq dataset published by Zhang et al (27). Expression values had been computed using the Magic-AceView pipeline and were given as normalized log(sFPKM) values [see (25) for details]. Survival information, ALT, and MYCN annotation were again taken from Roderweiser et al. (25). Survival analysis was done with groups with high and low expression of selected genes in survival analysis were generated by maximally selected rank statistics using the R package maxstat (version 0.7-25).

***Immunohistochemistry (IHC)***

Tumor tissue was fixed in formaldehyde (3.7-4.0% w/v, AppliChem), processed in an automated tissue processor (LOGOS, Milestone), and embedded in paraffin. Sections (4 µm) were mounted on glass slides (Superfrost+, Thermo Scientific) and heated for 3 h at 56°C. Following de-paraffinization and rehydration in a series of graded alcohols, heat induced epitope retrieval was performed in citrate buffer (C-9999, Sigma) using a Decloaking Chamber (Biocare Medical) set to 5 min at 110°C. Unspecific antibody binding sites were blocked with 5% goat serum (Sigma-Aldrich) in TBS containing 0.1% Tween 20 (TBST) (Sigma-Aldrich) for 1 h at RT, and endogenous peroxidase activity was quenched by incubation with BLOXALL® (Vector Laboratories) for 10 min followed by 2x5 min washes in TBST. Sections were incubated with primary antibodies against METTL3 and METTL14 in a humid chamber at 4°C overnight. Following 3 washes with TBST, the sections were incubated with HRP-conjugated anti-rabbit secondary antibodies (ImmPRESS Polymer Kit; MP-7451; Vector Laboratories) for 30 min followed by incubation with ImmPACT DAB (SK-4105; Vector Laboratories) for visualization. Sections were counterstained with Mayer's Hematoxylin and mounted (SignalStain® Mounting Medium, Cell Signaling Technology). Images were taken on a BH2 microscope with a UC30 camera (Olympus). Information regarding antibodies and dilutions used in the experiment are provided in the Supplementary Table S1.
